# Multi Epitope Vaccine Prediction Against Aichi Virus using Immunoinformatic Approach

**DOI:** 10.1101/795427

**Authors:** Asma Ali Hassan Ali, Sahar Abdeen Abdalla Mohamed, Marwa Abdulrahman Omer Musa, Amani Faisal Bushra ELtahier, Walaa Mohammed Alsamani Abdelrahman Ali, Elaf Ahmed Mohammed Ali Ibrahim, Ahmed Hamdi, Mohamed A. Hassan

**Affiliations:** Department of bioinformatics, Africa city of technology, Khartoum, Sudan; Faculty of science, University of Khartoum, Khartoum, Sudan; Faculty of medical laboratory science, Riyadh international college, Khartoum, Sudan; Alshifa hospital, Khartoum, Sudan; UMbada maternal and child health hospital, Khartoum, Sudan; Department of Bioinformatics, DETAGEN Genetic Diagnostics Center, Kayseri, Turkey

**Keywords:** immunoinformatic, peptide vaccine, Epitope, Aichi virus, immune epitope database IEDB

## Abstract

Aichi virus, AiV is single stranded negative sense RNA genome belonging to the genus *Kobuviru*, a family of *Picornaviridae* that causes severe gastroenteritis. There is no treatment or vaccine for it, thus the aim of this study is to design a peptide vaccine using immunoinformatic approaches to analyze the viral Protein VP1 of AiV-1 strain, to determine the conserved region which is further studied to predict all possible epitopes that can be used as a peptide vaccine. A total of 38 Aichi virus VP1 retrieved from NCBI database were aligned to determine the conservancy and to predict the epitopes using IEDB analysis resource. Three epitopes predicted as a peptide vaccine for B cell was (PLPPDT, PPLPTP, and LPPLPTP). For T cell, two epitopes showed high affinity to MHC class I (FSIPYTSPL and TMVSFSIPY) and high coverage against the whole world population. Also, in MHC class II, three epitopes that interact with most frequent MHC class II alleles (FTYIAADLR and YMAEVPVSA) with high coverage in the whole world population. For both MHCI and MHCII the T-cell peptide with the strongest affinity to the worldwide population was FSIPYTSPL.

Peptide vaccine against AiV is powerfully displace the normal produced vaccines based on the experimental biochemistry tools, as it designed to handle with a wide range of mutated strains, which will effectively minimize the frequent outbreaks and their massive economical wastage consequences.

## Introduction

Aichi virus 1 (AiV-1) is a small round virus its diameter is about 30 nm, belongs to the *Kobuvirus* genus, *Picornaviridae* family, has been considered as the responsible agent for human gastroenteritis and children hospitalization with acute diarrhea probably passed by fecal-oral ways through polluted water or food(3–1).

In 1989, Aichi virus has been identified for the first time, as the likely cause of oyster-related nonbacterial gastroenteritis in a stool specimen of a Japanese patient. AiV-1 was also identified in many Asian countries, for example, 5 (2.3%) of 222 Pakistani children between 1990-1991, and 5 (0.7%) of 722 Japanese travelers returned from tours to Southeast Asian countries between April 1990 and March 1992, the RNA was detected in 54 (55%) of 99 fecal specimens from the patients in 12 (32%) of 37 gastroenteritis epidemics in Japan.

In Finland, Of the 468 stool samples analyzed from the hospital-based epidemiological study, three samples were -positive for Aichi virus (0-6% incidence), a 485 German serum samples panel was presented for Aichi virus antibody, identifying a seroprevalence of 76%. Aichi virus was also found in 28 of 912 fecal samples which were negative for astrovirus, sapovirus, norovirus, adenovirus, and rotavirus and were assembled in Japan, Bangladesh, Thailand, and Vietnam during 2002 to 2005. A research carried out from January 2003 to June 2005 revealed, Aichi virus was the cause of 3.5% of 632 cases of Tunisian children presenting in hospitalization (252 children) or dispensaries (380children) for acute diarrhea. In Italy, the virus was found in 3/170 (1.8%) of the analyzed specimens. The AiV-1 positive samples were of various geographic origin (1, 4–9).

The genome length of AiV-1 is about 8,400 nucleotides, t is positive sense, single stranded RNA, containing an open reading frame encoding a 2,433-residue long polyprotein. The polyprotein cleaves co-translationally and post-translationally to viral capsid proteins VP0, VP3, and VP1, leader protein (L-protein) and nonstructural proteins that control the AiV-1 replication in the infected cell(10, 11).

On the outer surface, a polyproline helix structure, which was not identified formerly in picornaviruses, exists at the VP1 C terminus, a place where integin binding motifs are found in many other picornaviruses. A peptide linked to this polyproline motif attenuates virus infectivity to some extent, possibly blocking host-cell attachment. This may guide cellular receptor identification(12).

AiV can cause severe gastroenteritis and could be lethal for children below five years old, especially in developing countries. Moreover, there is no available vaccine or effective antiviral treatment has been introduced(12).

Our aim is to design a vaccine for Aichi virus using peptide of its vp1 as an immunogen to stimulate an immune response.

Producing vaccine with the experimental biochemistry tools are high-cost, laborious and sometimes does not work effectively, moreover, the vaccine that formulated from attenuated or inactivated microorganism contains immunity induction proteins of that probably develops allergenic or reactogenic responses. For that reason, in silico proper protein residues epitopes prediction is considered to be helpful in peptide vaccine production with a great impact immunogenic and little amount of allergenic effect (13–16). Numerous researches demonstrated the immunological efficacy of peptide-based vaccines against infectious illnesses. The advancement of peptide-based immunizations has fundamentally progressed with the particular epitope’s identification gotten from infectious pathogens. Comprehension of the antigen recognition molecular basis and HLA binding motifs has brought about the improvement of the designed vaccine depending on motifs prediction to bind to host class I or class II MHC(17). There are several types of research have been conducted considering immunoinformatic predication and in sillico modeling of epitope-based peptide vaccine against many viruses (18–22).

## 2. Materials & Method

### 2.1 Sequence of protein recovery

A total of 38 protein strains sequences of Aichi virus vp1 were retrieved from NCBI (https://www.ncbi.nlm.nih.gov/) in October 2018. Those 38 strains sequences were collected from different parts in the world (Japan, Germany and South Korea), The Achi virus VP1 strains, area of collection and their accession numbers are listed in the table (1).

**Table (1):**
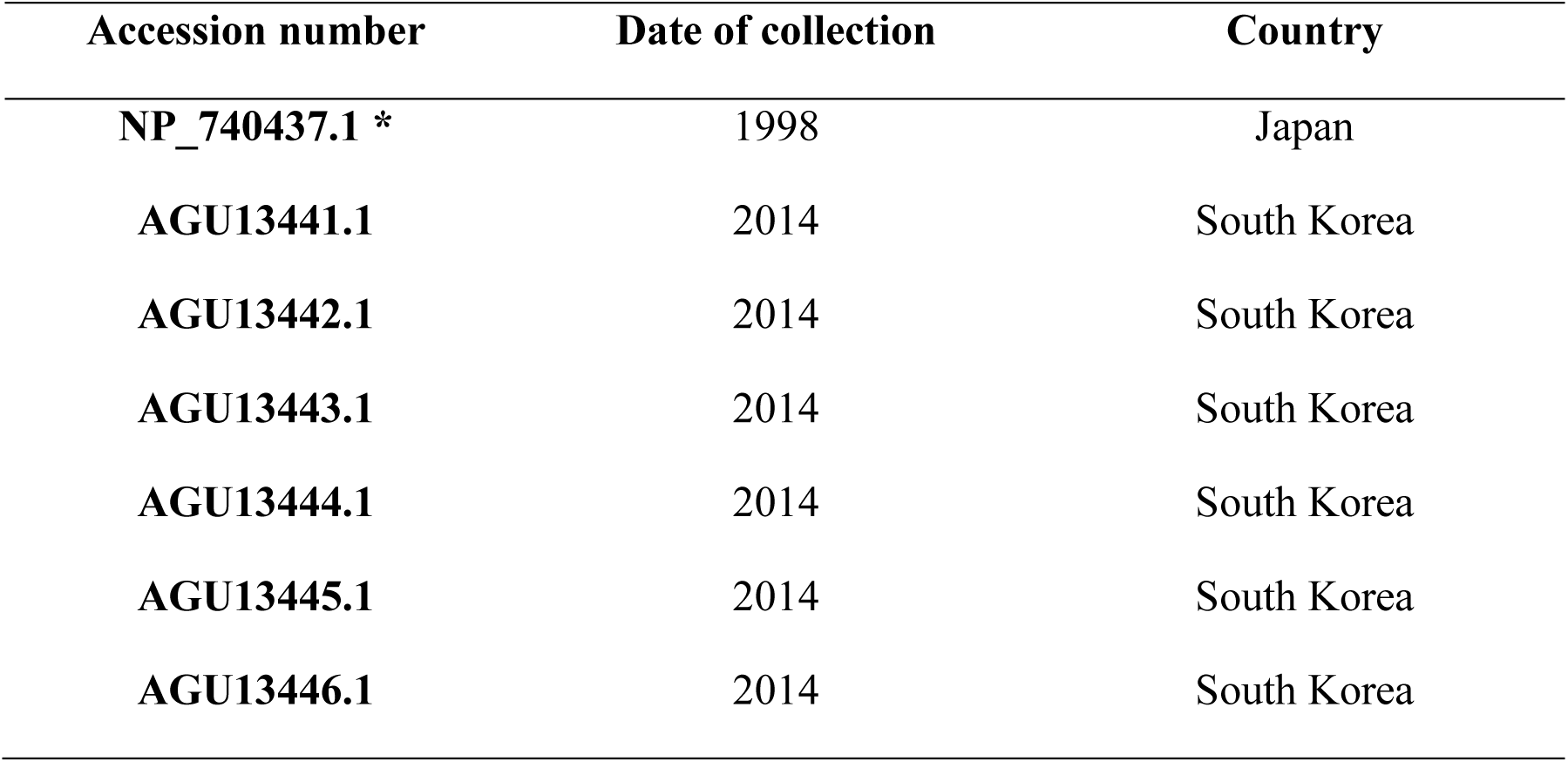
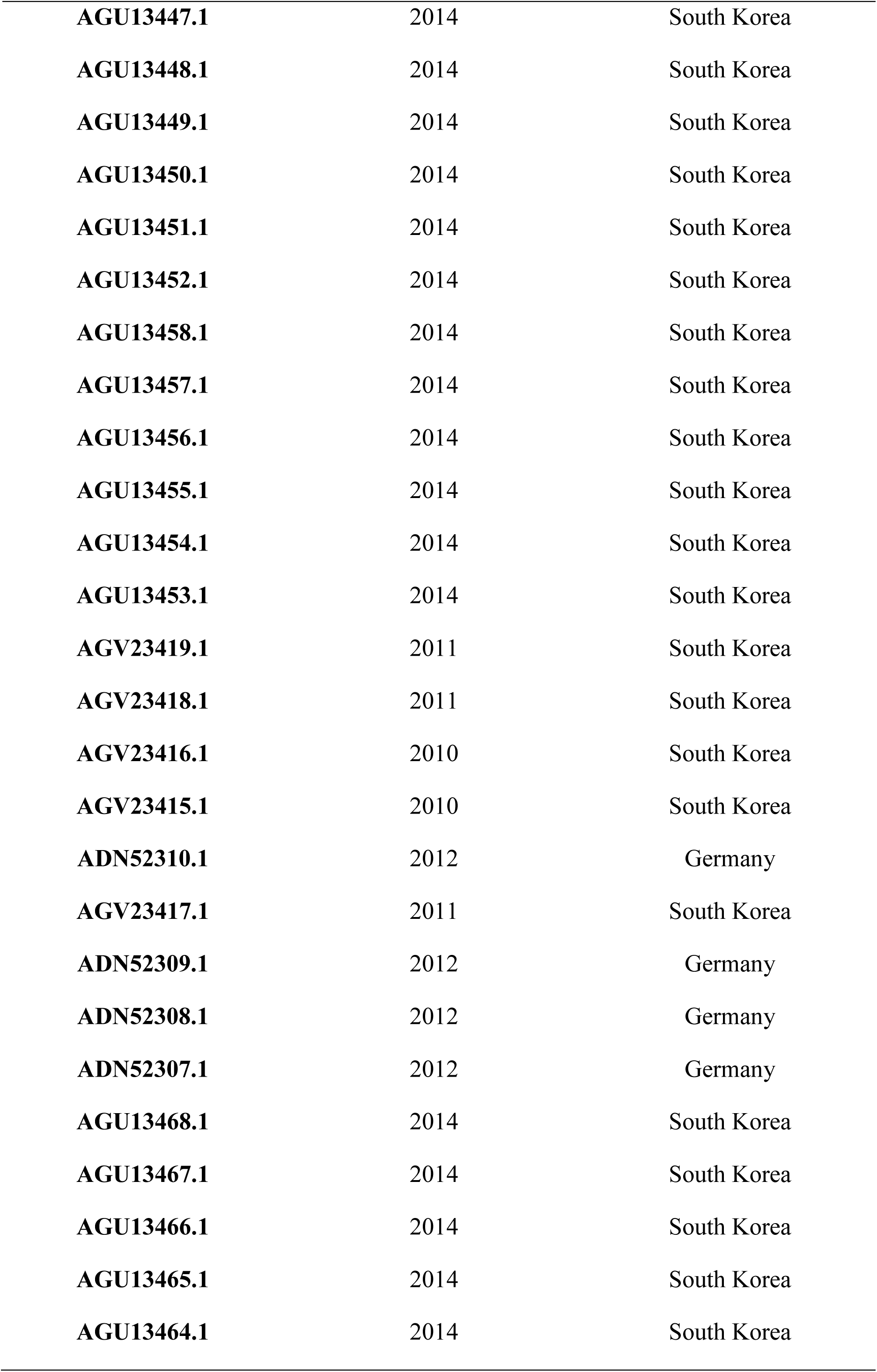
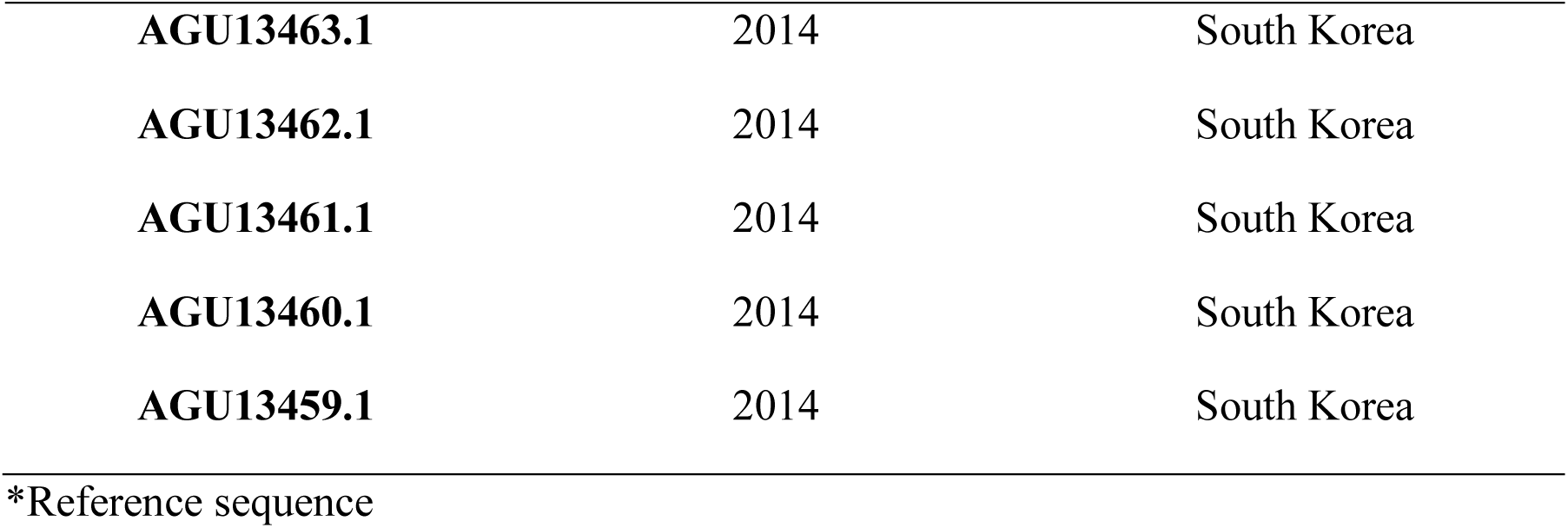
virus strains, accession numbers, areas and years of collection:

### 2.2 Phylogenetic and alignment

The retrieved sequences were submitted to Phylogenetic and alignment tools MEGA7.0 to determine the common ancestor of each strain and the conservancy (23) (https://www.megasoftware.net/). The alignment and phylogenetic tree were presented in Figure (2).

**Figure (1):**
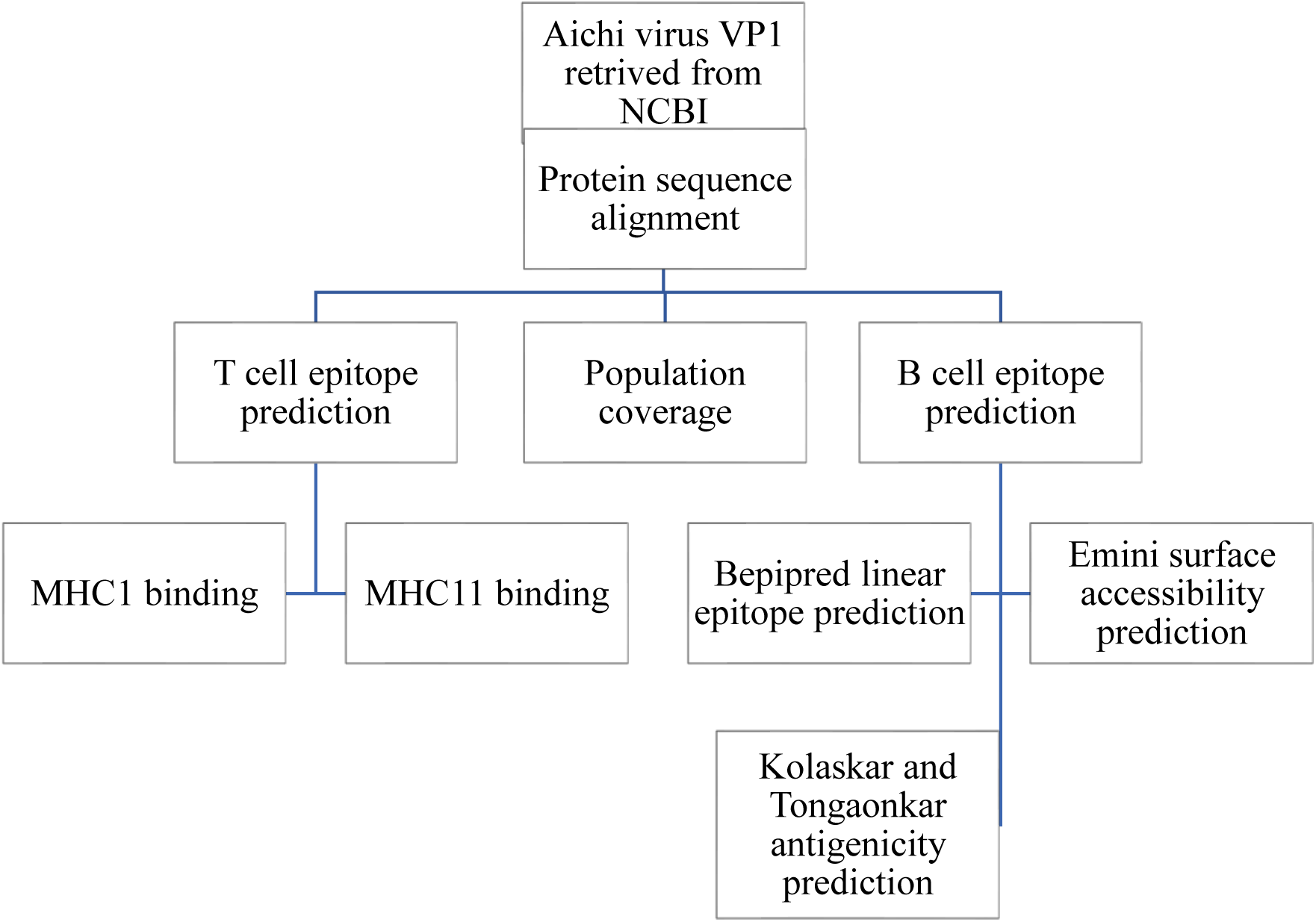
Flowchart of the epitope prediction processes for B cell and T cell.

**Figure (2):**
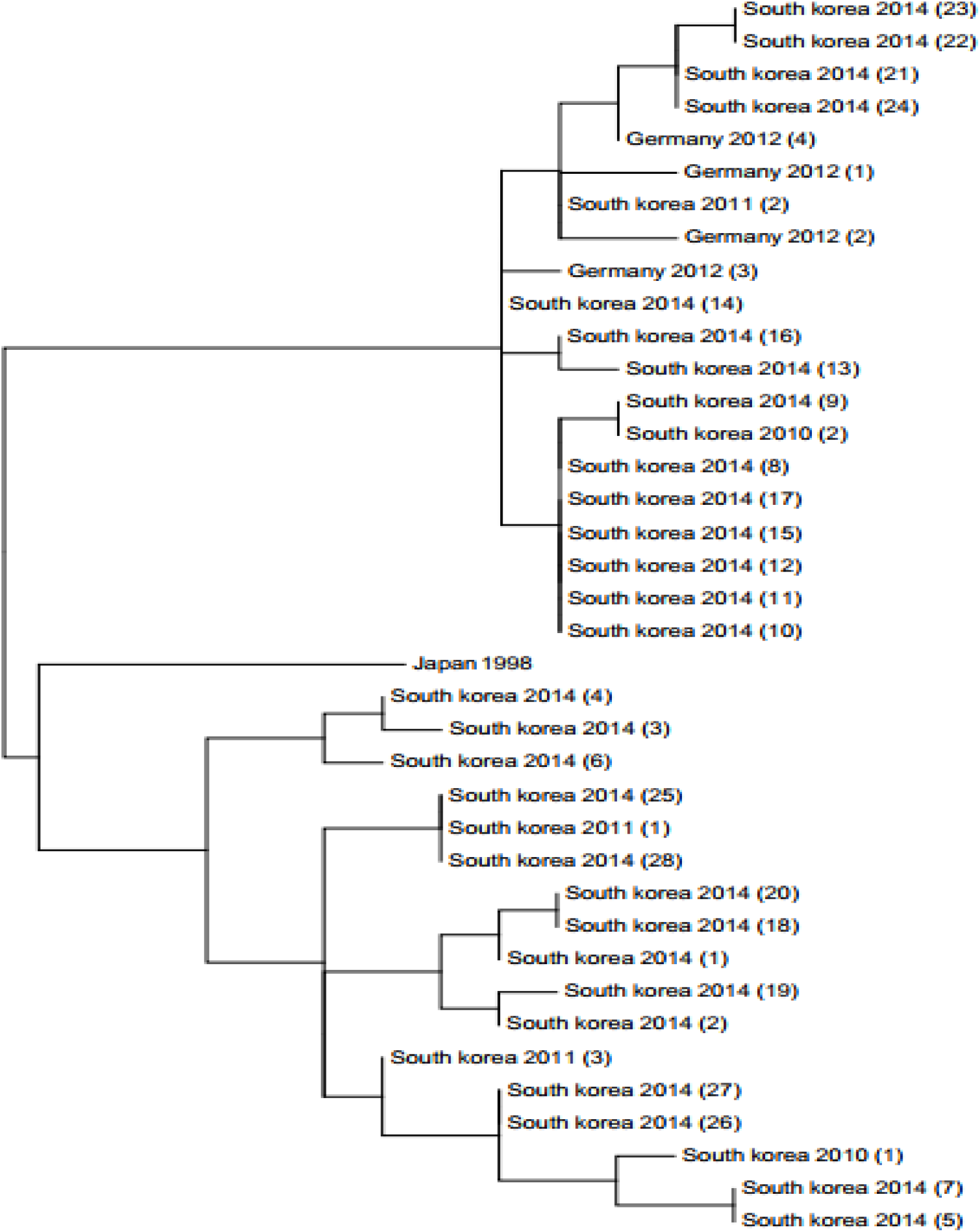
Cladogram shows the relationship between different strains of Aichi virus.

### 2.3 Determination of conserved regions

The chosen sequences were aligned by using multiple sequence alignment (MSA) BioEdit software (version 7.2.5.0) (24) to obtain the sequences of the conserved regions, aligned with Clustal W were used to determine the conserved regions in all Aichi virus VP1, protein sequences shown in figure(3). Peptides chose as epitopes were analyzed by different prediction tools from Immune Epitope Database, IEDB analysis resource (https://www.iedb.org/) (25).

**Figure (3):**
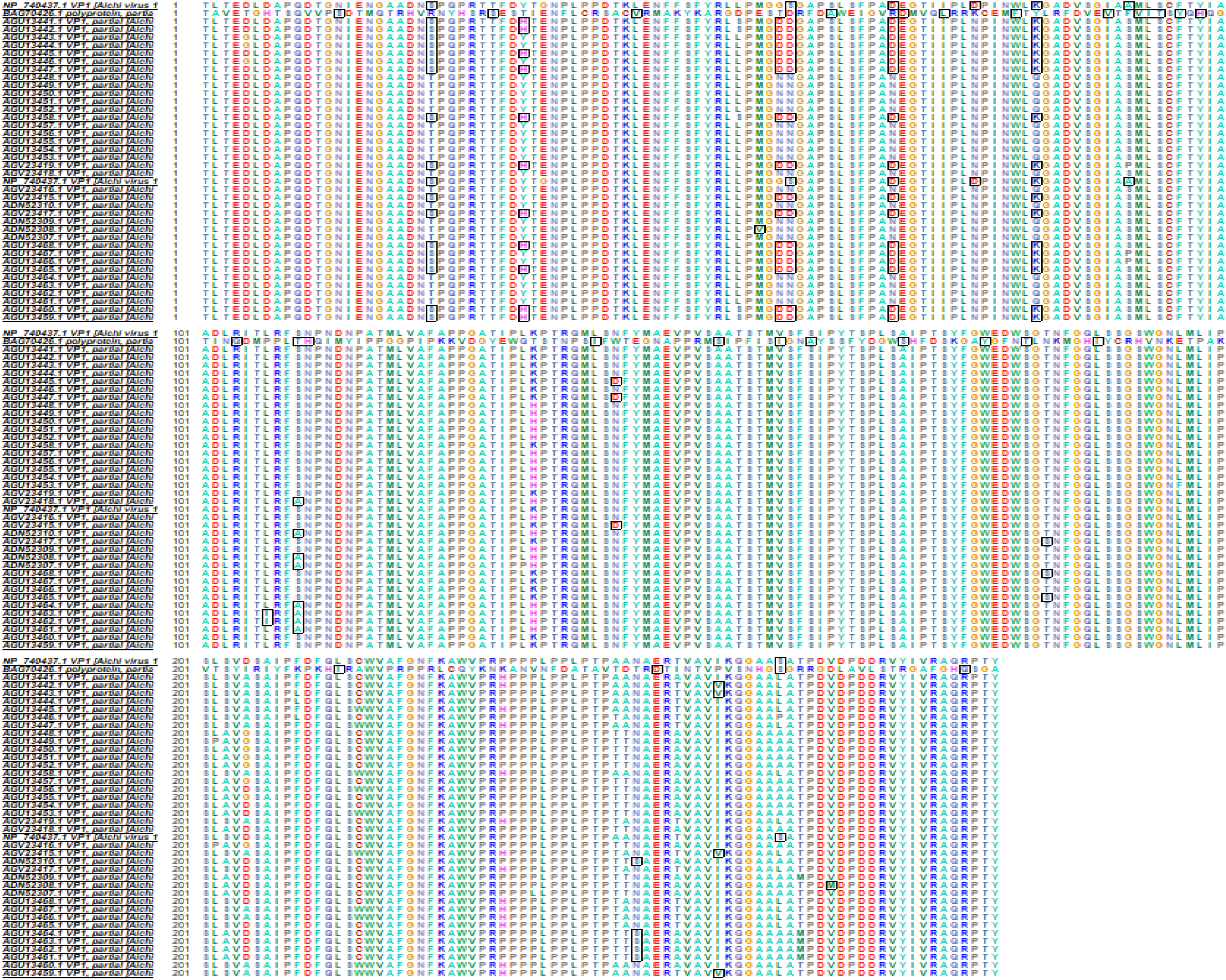
Protein sequence Alignment using Bioedit software for a showing of the conservancy.

### 2.4 Binding prediction of B cell epitope

The reference sequence of Aichi virus VP1 was subjected to many B cell tests in IEDB webpage (http://tools.iedb.org/bcell/) (26).

#### 2.4.1 linear B cell epitopes prediction

The linearity of the peptide was studied using Bepipered Linear Epitope Prediction in the immune epitope database (http://toolsiedb.ofg/bcell/) (27), which had a threshold value of 0.35.

#### 2.4.2 Surface accessibility prediction

Using Emini surface epitope prediction from IEDB (http://tools.immuneepitope.org/tools/bcell/iedb) (28), Epitopes of the surface accessible were predicted from the region in which threshold holding value was 1.

#### 2.4.3 Epitope antigenicity prediction

kolaskar and tongaonker antigenicity in IEDB (http://tools.immuneepitope.org/bcell/) (29) was used to determine the antigenic sites with a default threshold value of 1.010.

### 2.5 Binding prediction of T cell epitope

#### 2.5.1 Binding predictions of MHC class 1

The peptide binding analysis to major histocompatibility class I molecules was evaluated by IEDB MHC I estimated tool at (http://tools.iedb.org/mhci/).

Prediction methods were achieved by Artificial Neural Network (ANN), The analysis was done for alleles with peptides length of 9-mers and which have scored equal or less than 500 Half Maximal Inhibitory Concentration (IC50) (30) which was chosen for further analysis.

#### 2.5.2 Binding Predictions of MHC class 2

Analysis of peptide binding to MHC2 molecules was assessed by the IEDB MHC II prediction tool at (http://tools.immuneepitope.org/mhcii/) (31) For MHCII binding Prediction human allele references set were used (32). We used Artificial Neural Networks (ANN) to identify both the binding affinity and MHCII binding core epitopes. All conserved epitopes that bind to many alleles with a score equal or less than 500 half maximal inhibitory concentration (IC50) were selected for further analysis.

### 2.6 Population Coverage Calculation

All proposed MHC class I & class II epitopes from Aichi virus vp1 protein were used for population coverage to whole world population with selected MHC I and MHC II binding alleles using IEDB population coverage calculation tool at (http://tools.iedb.org/tools/population/iedb_input) (33).

### 2.7 Homology Modeling

The reference sequence of Aichi virus protein was sent to CPH server (http://www.cbs.dtu.dk/services/CPHmodels/index_prf2013.php) (34) to determine the 3D structure. This 3D structure was visualized using chimera version (1.8) from chimera package that accessed from the chimera web site (https://www.cgl.ucsf.edu/chimera/docs/credits.html) (35).

## 3. Results

### 3.1 B-cell epitope prediction

The Bepipred linear epitope prediction, Kolaskar, and Tongaonkar results and Emini surface accessibility prediction results were recorded by subjected reference sequence of Aichi virus (vp1) in IEDB table 2, figures 4, 5 and 6.

**Table (2):**
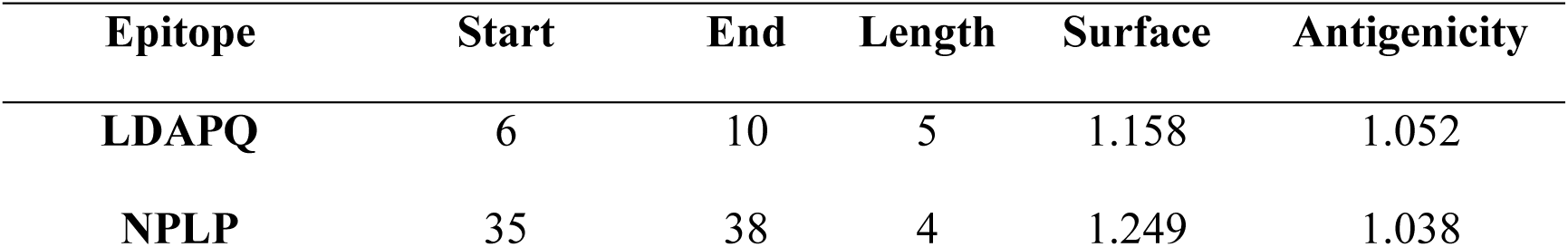
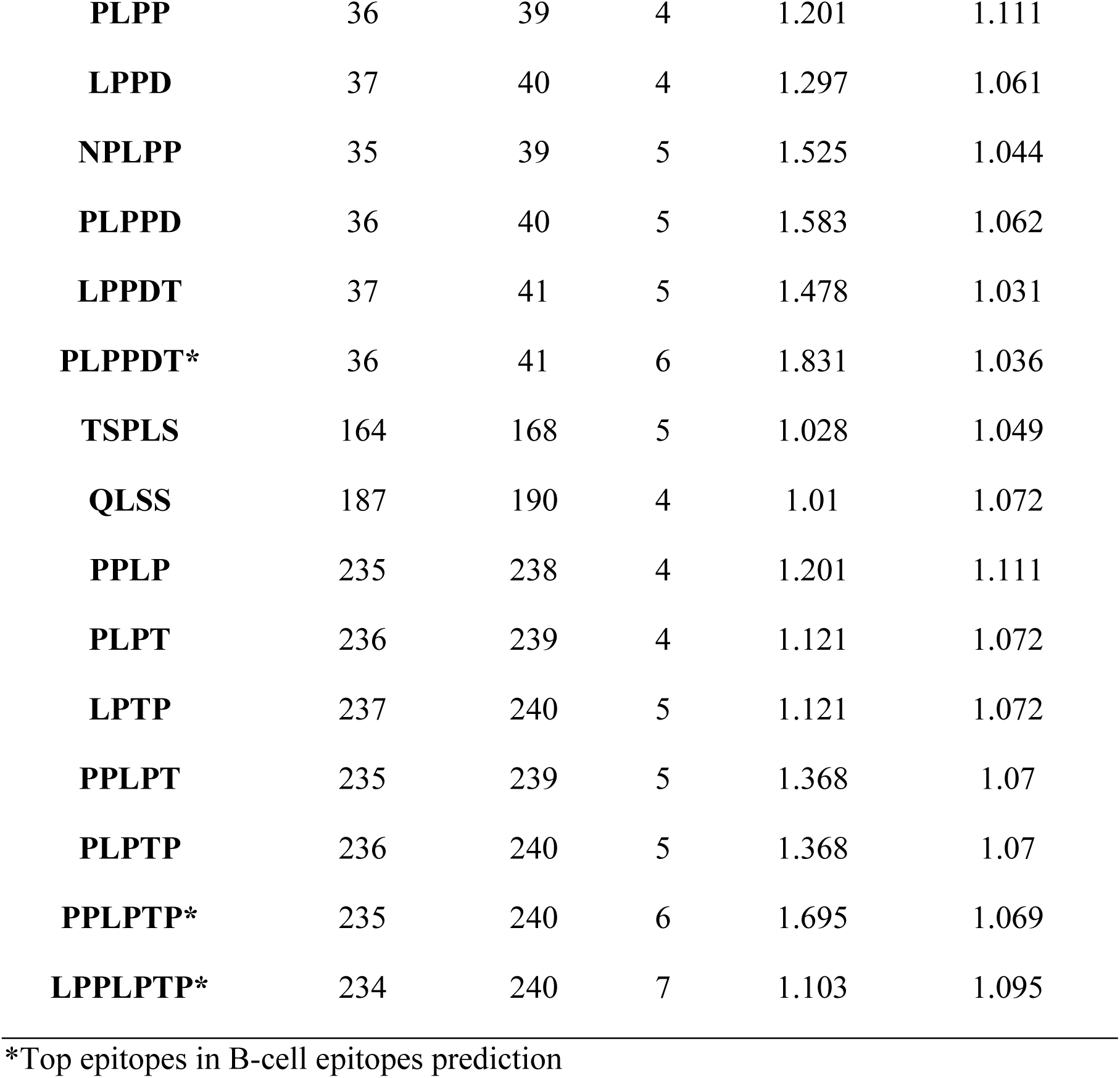
Results of B-cell epitopes prediction.

**Figure (4):**
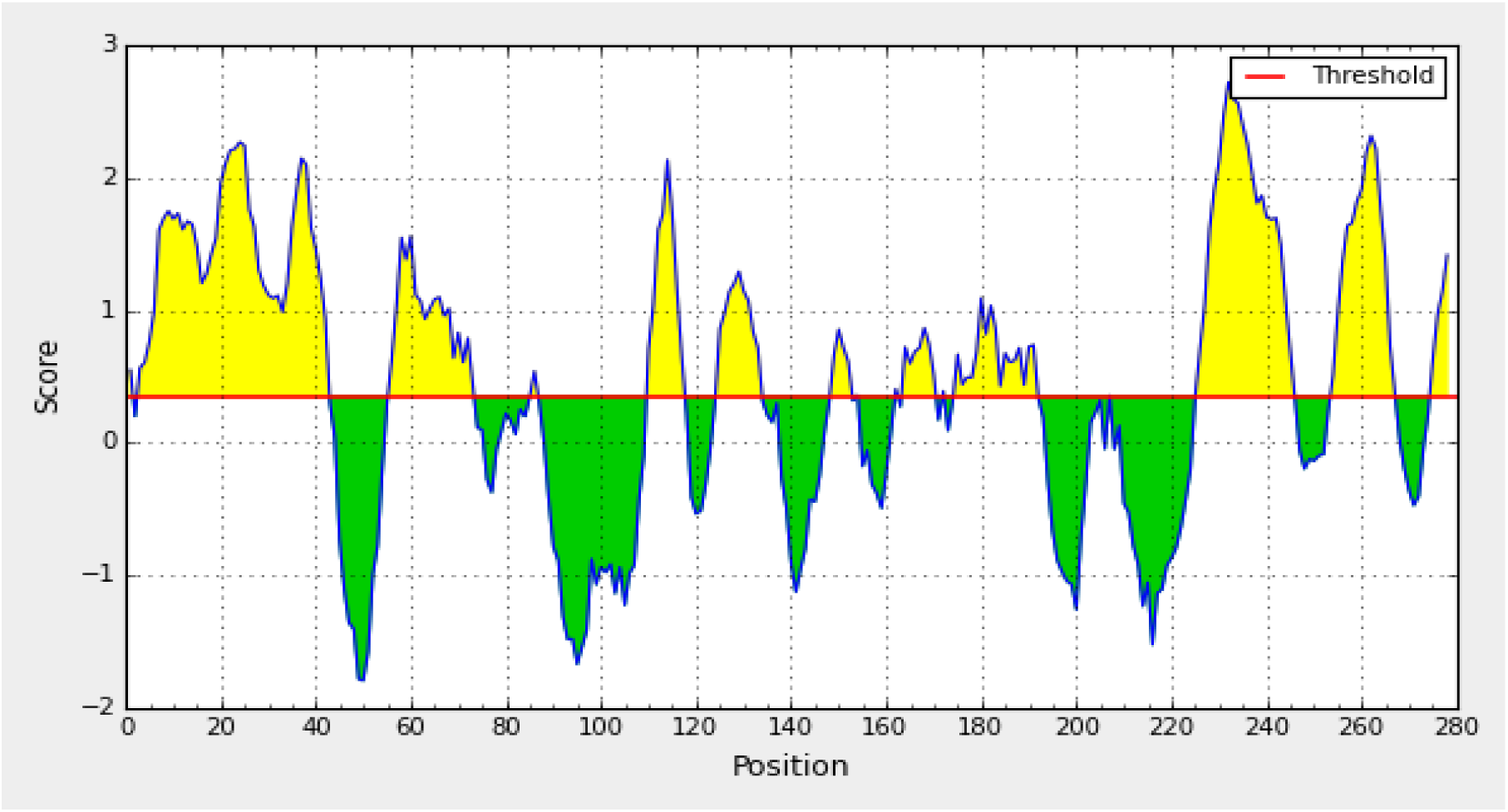
Bepipred linear epitope prediction. Yellow areas above threshold (red line) are proposed to be a part of B cell epitope. While green areas are not.

**Figure (5):**
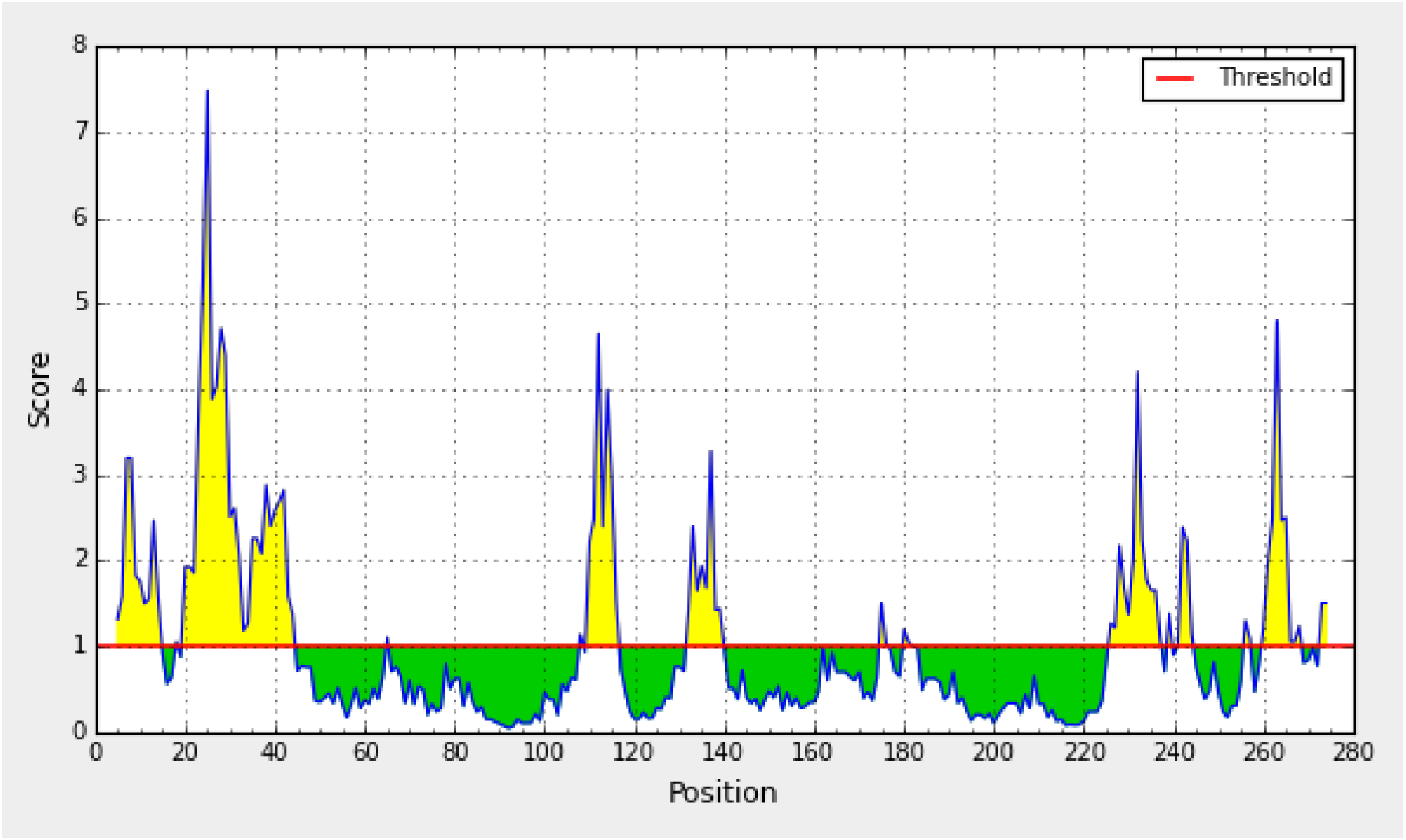
Emini surface accessibility. Yellow areas above threshold (red line) are proposed to be a part of B cell epitope. While green areas are not.

**Figure (6):**
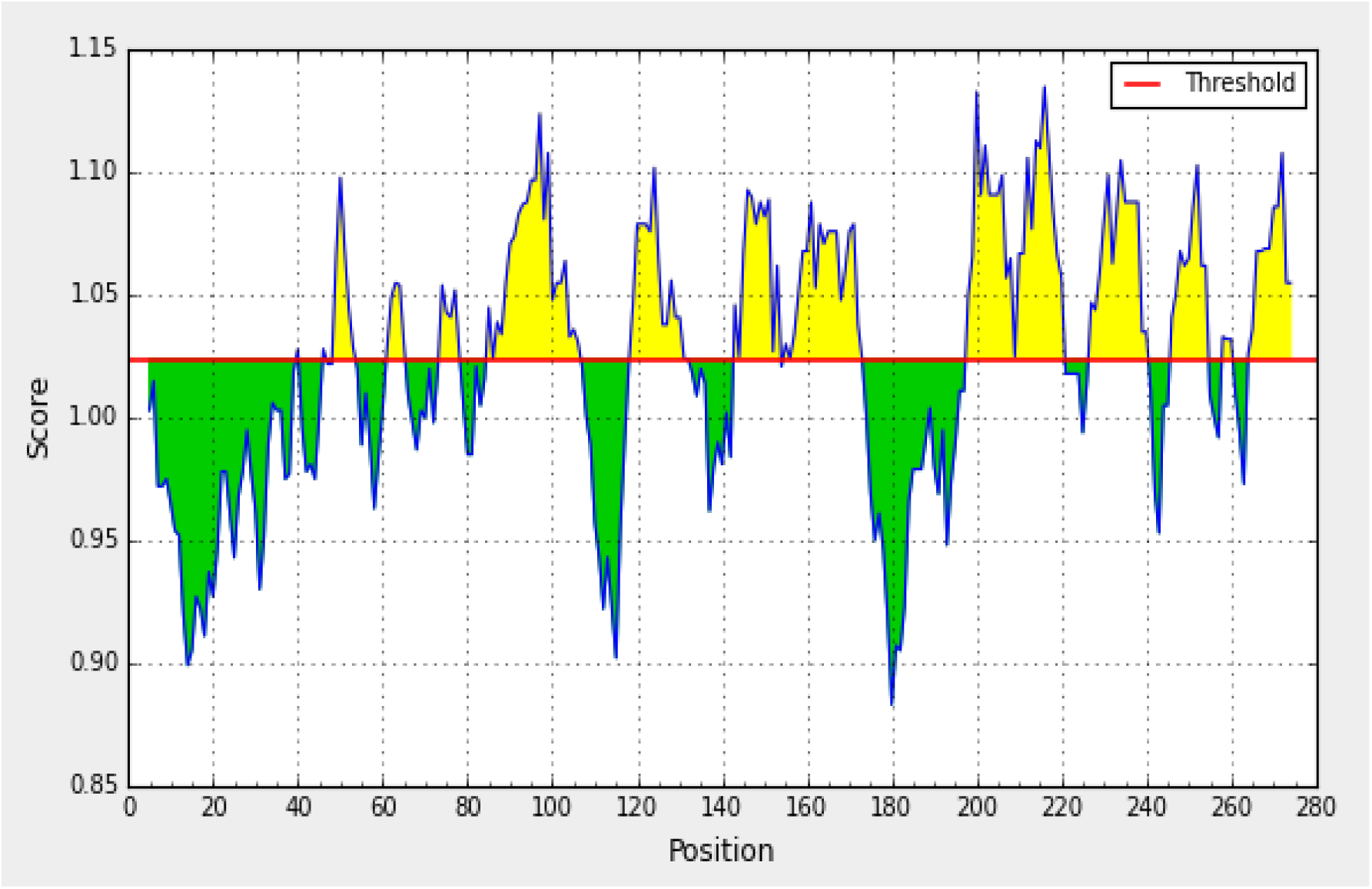
Kolaskar and togaonkar antigenicity. Yellow areas above threshold (red line) are proposed to be a part of B cell epitope. While green areas are not.

**Figure (7):**
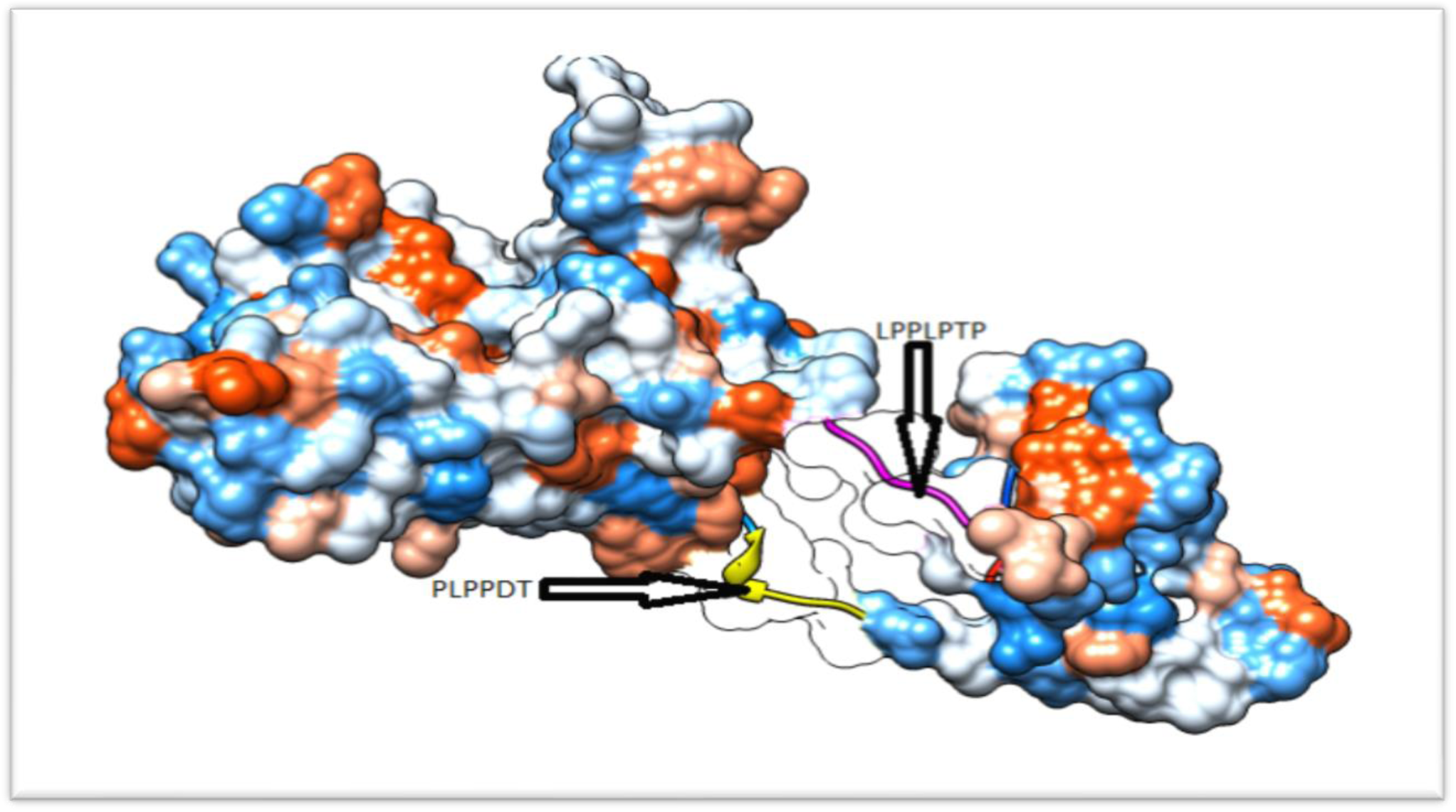
3D structure of predicted B cell epitopes of vp1 Protein in Aichi virus.

### 3.2 Prediction of cytotoxic T-lymphocyte epitope and interaction with MHC 1

The (vp1) reference sequence protein of Aichi virus was submitted in the IEDB MHC-1 binding prediction tool to predict epitopes interact with MHC-1 alleles.

**Table (3):**
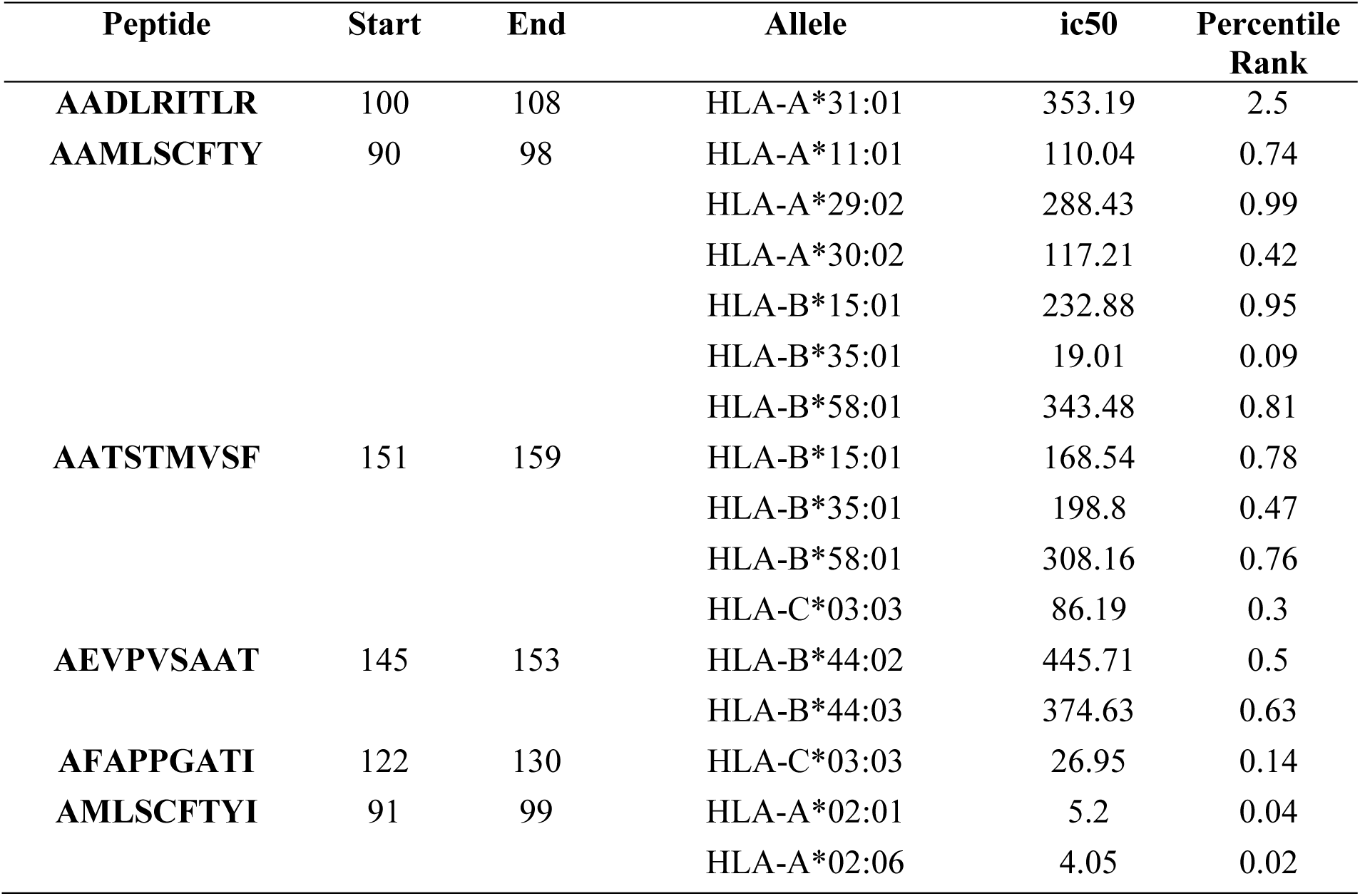
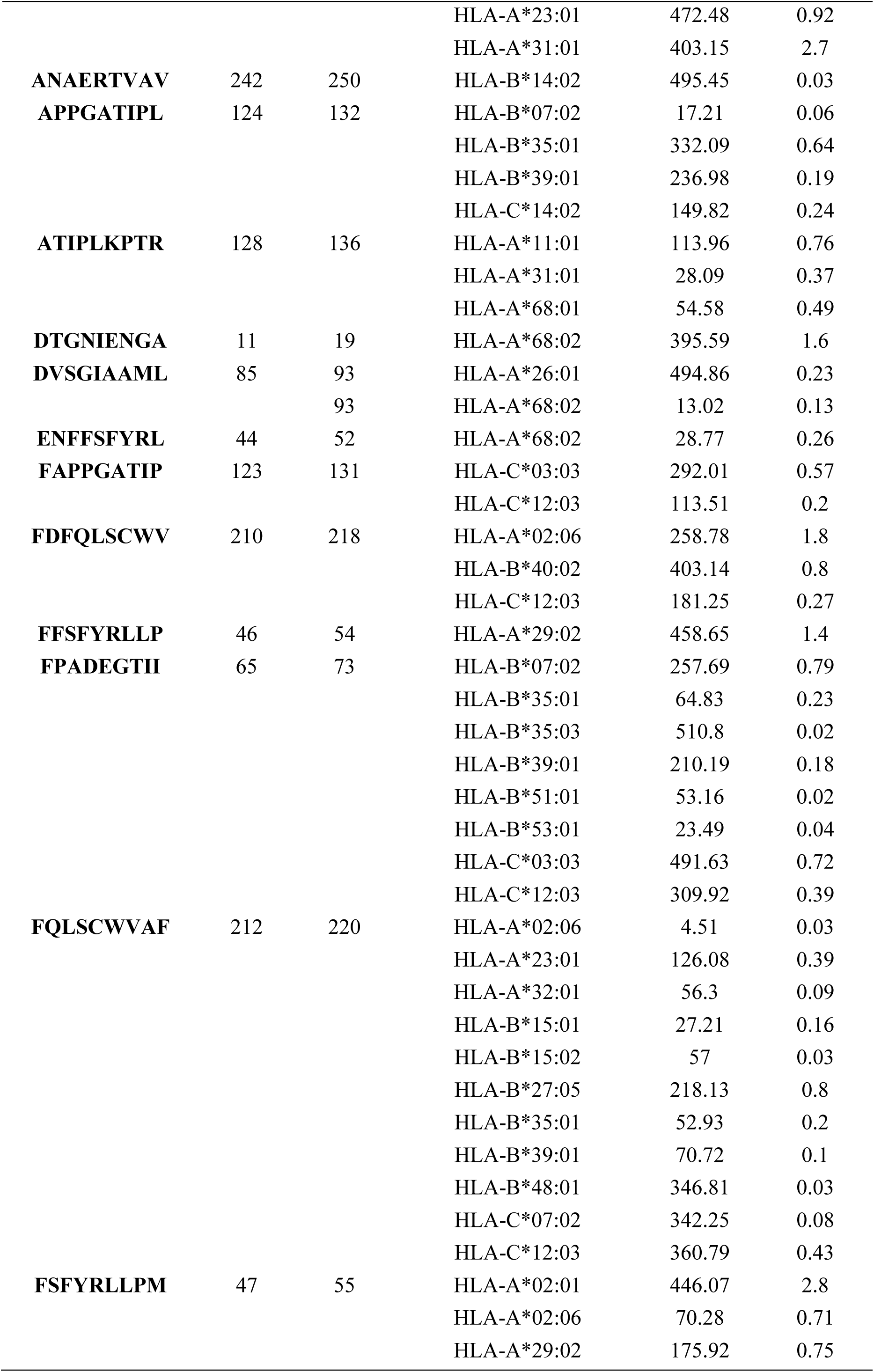
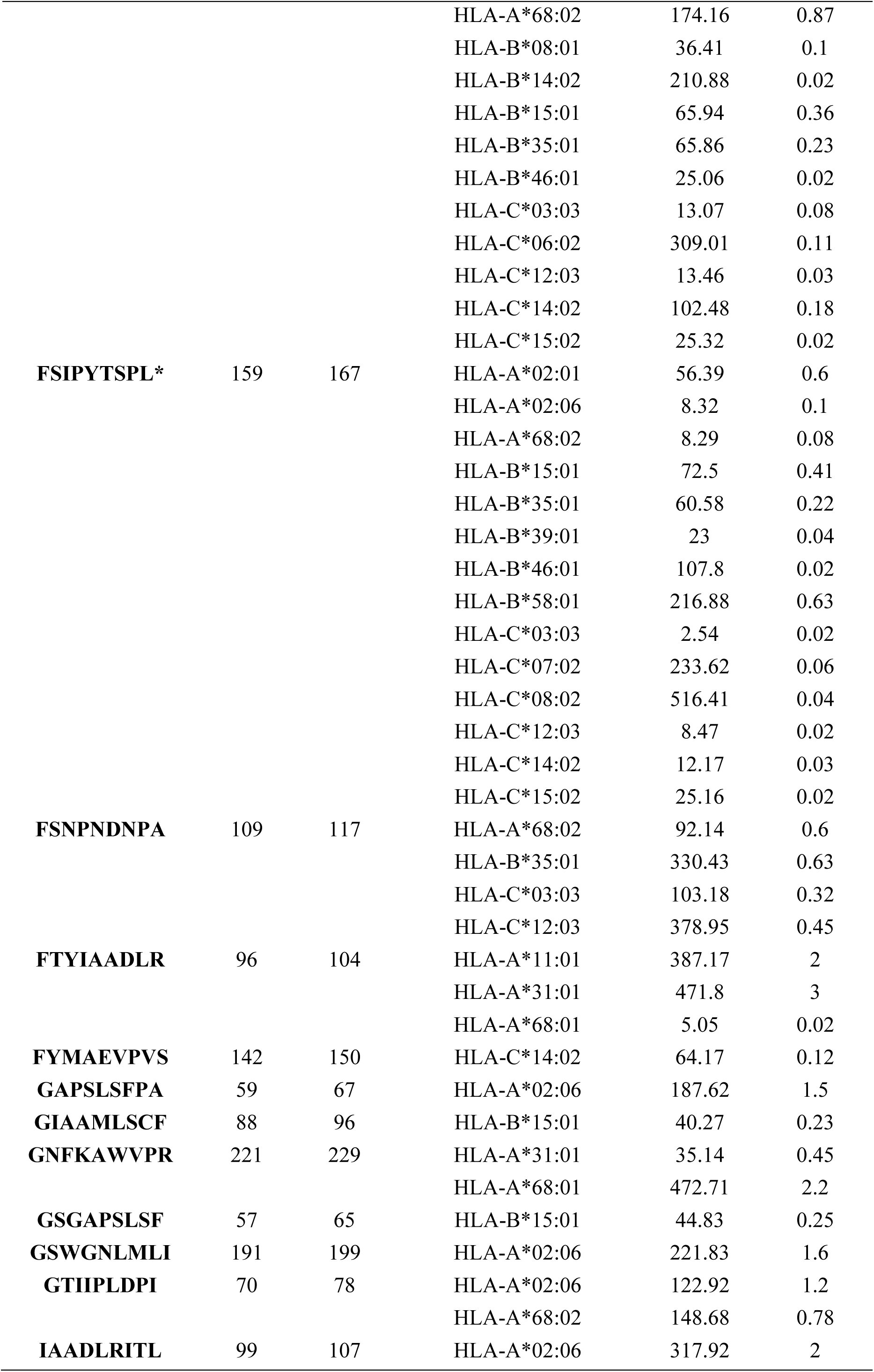
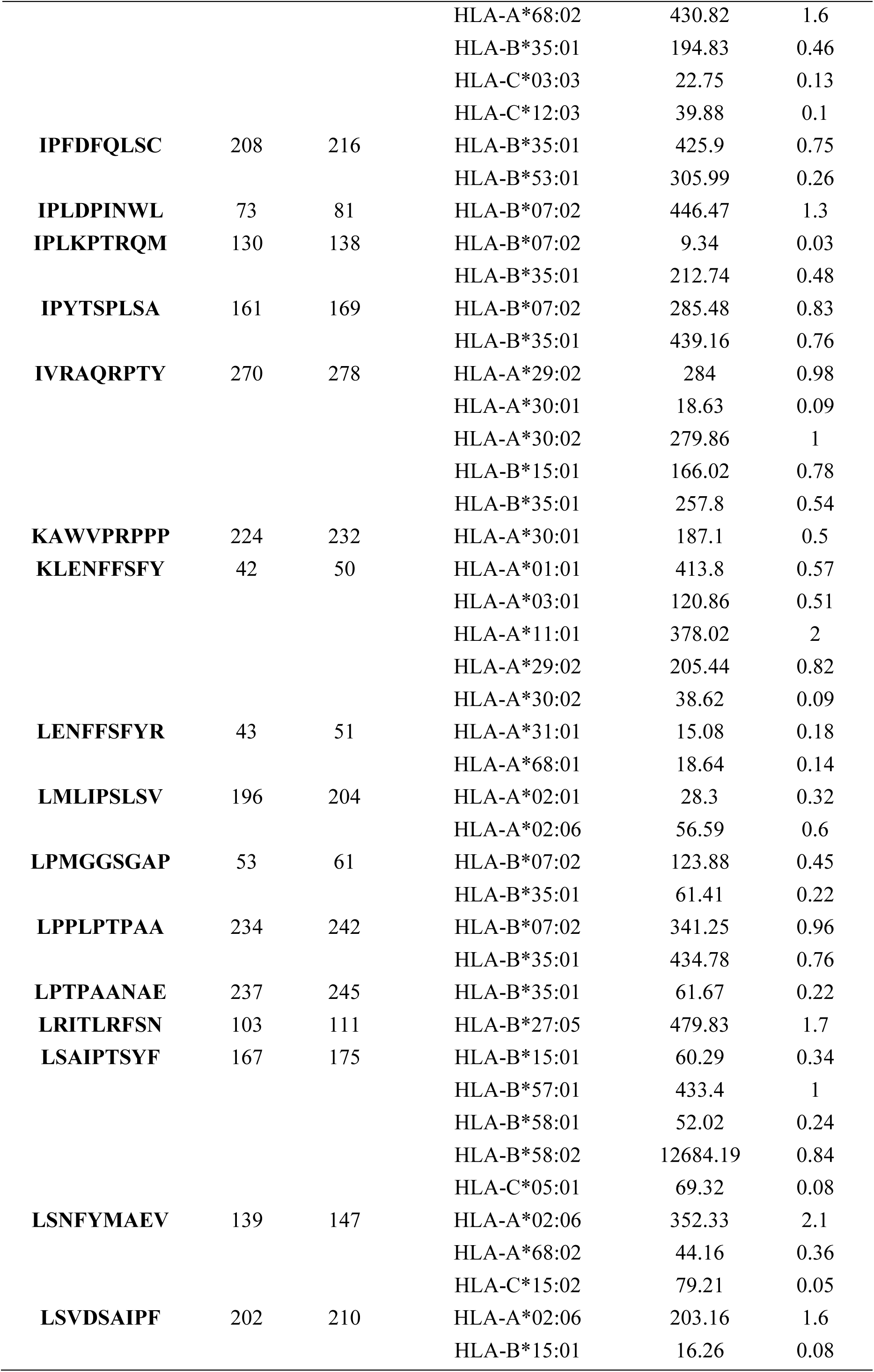
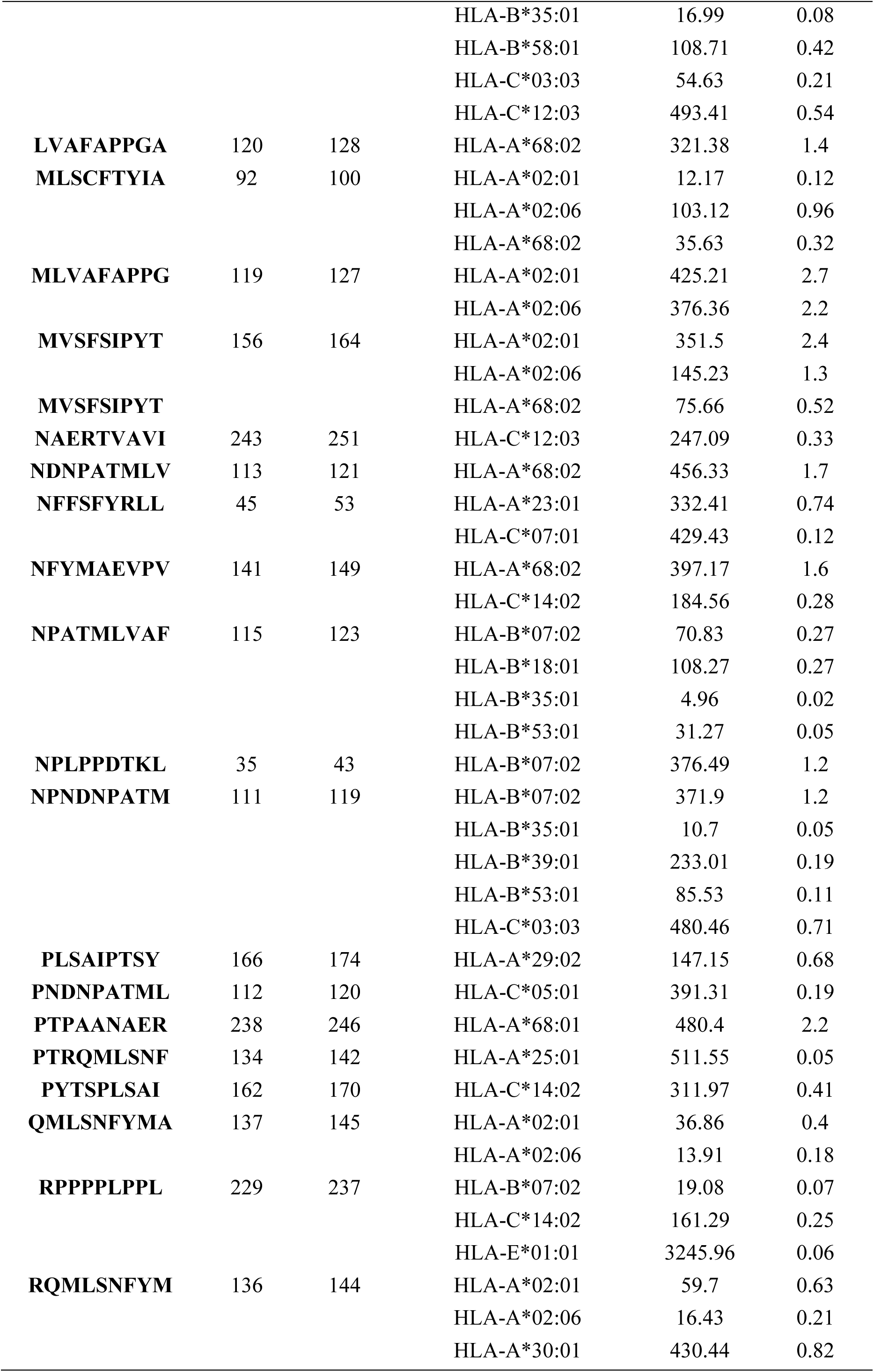
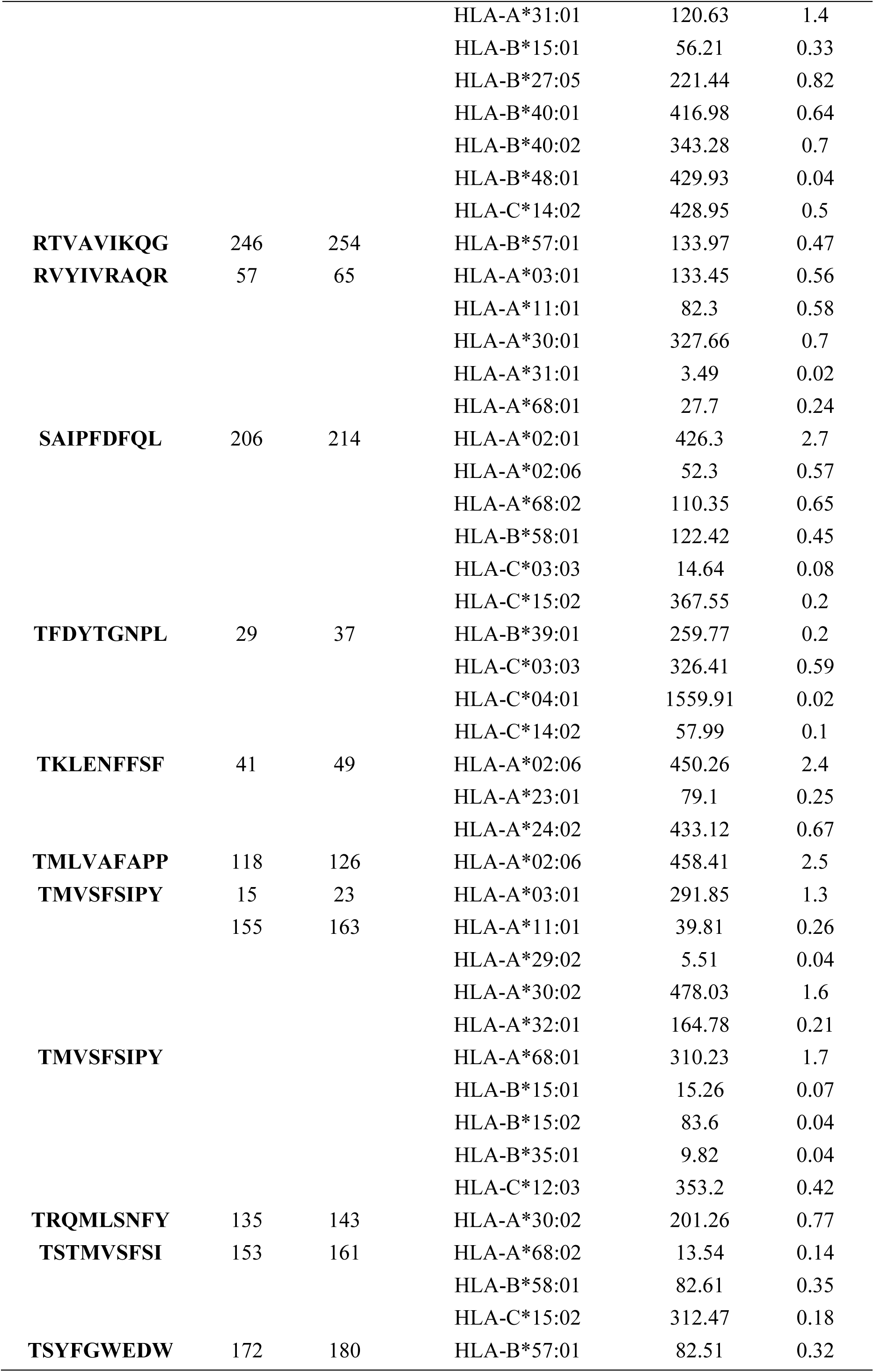
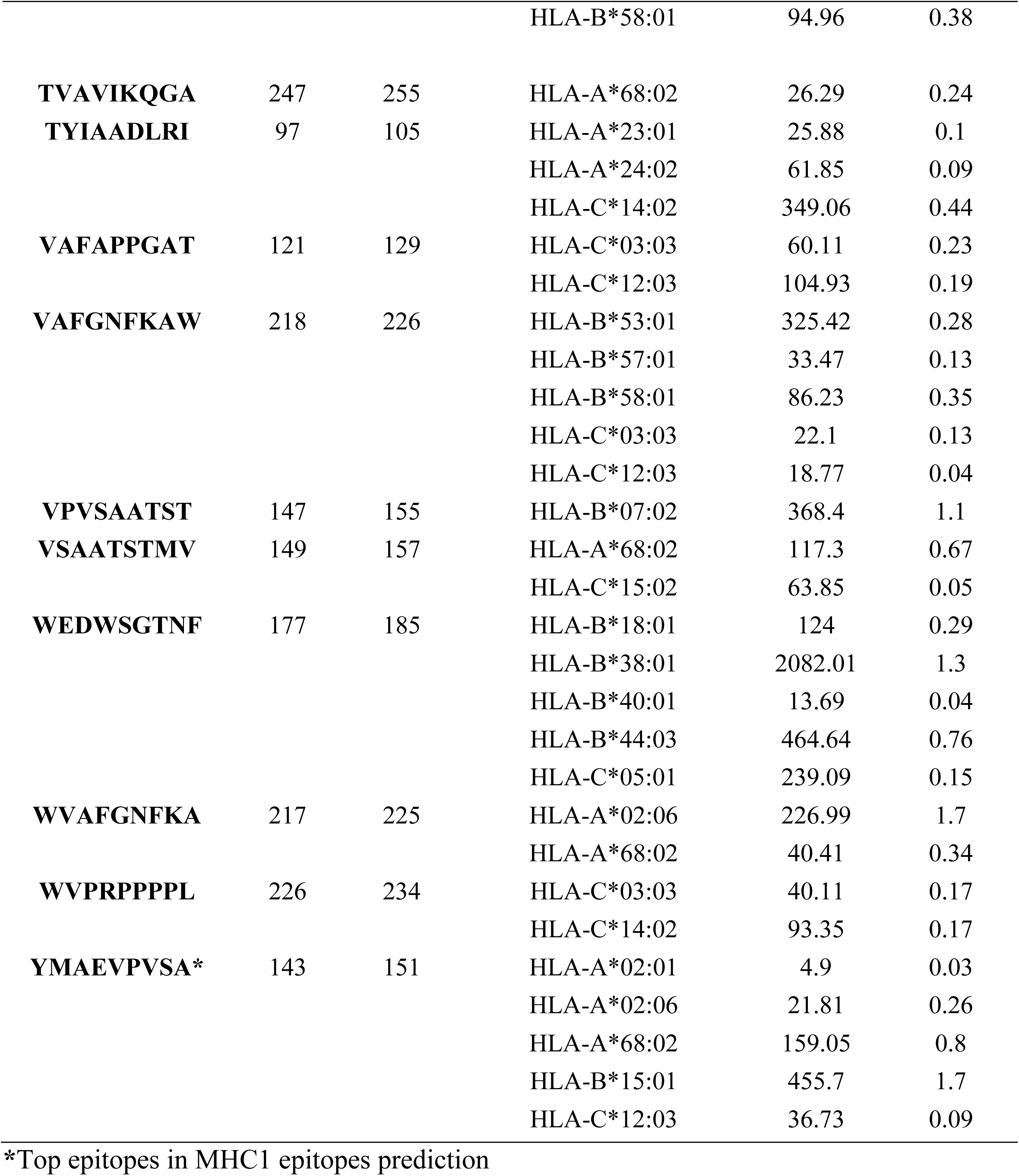
Result of predicted peptides that interact with MHC1:

**Figure (8):**
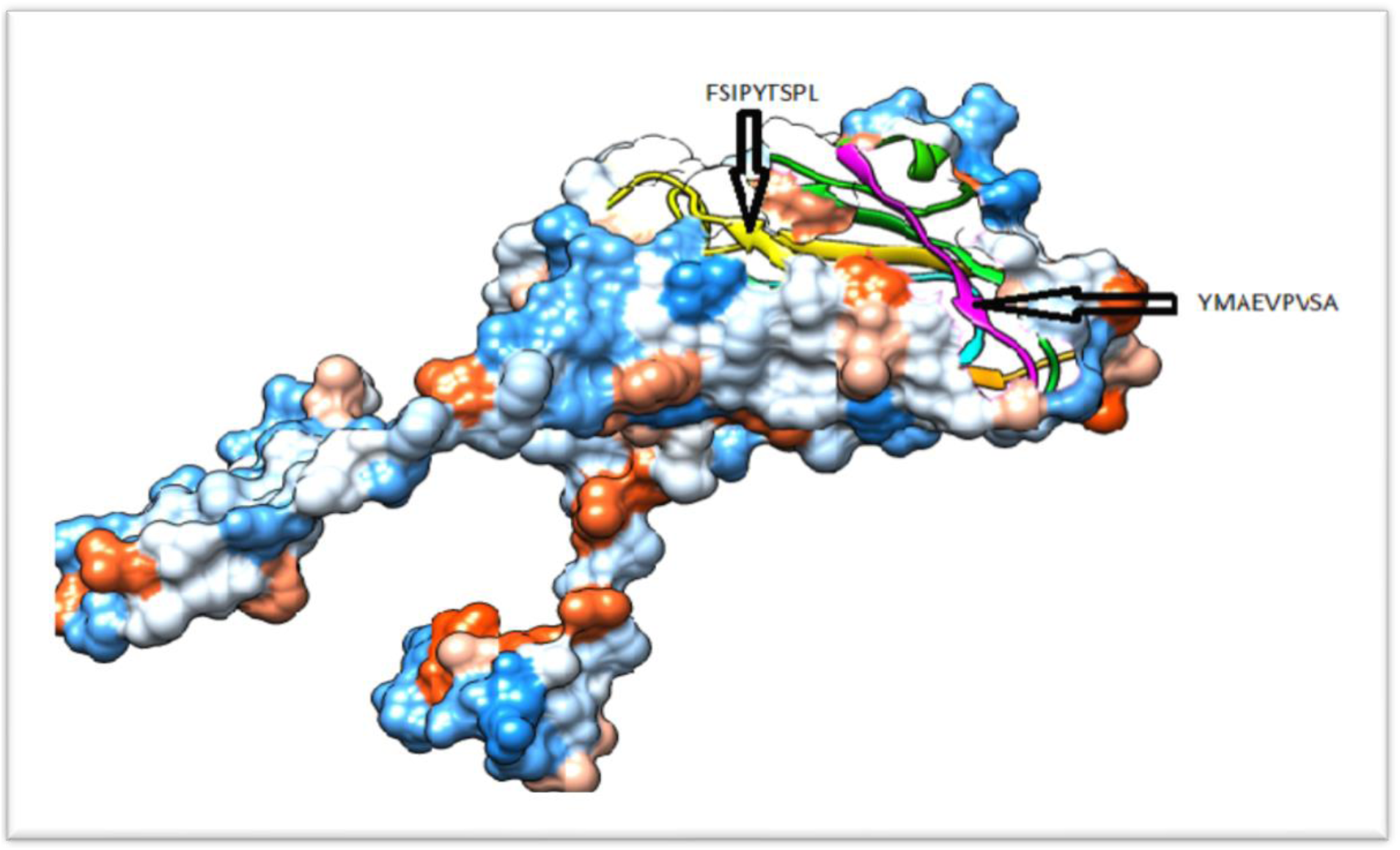
3D structure of cytotoxic T cell epitopes interacts with MCH1.

**Figure (9):**
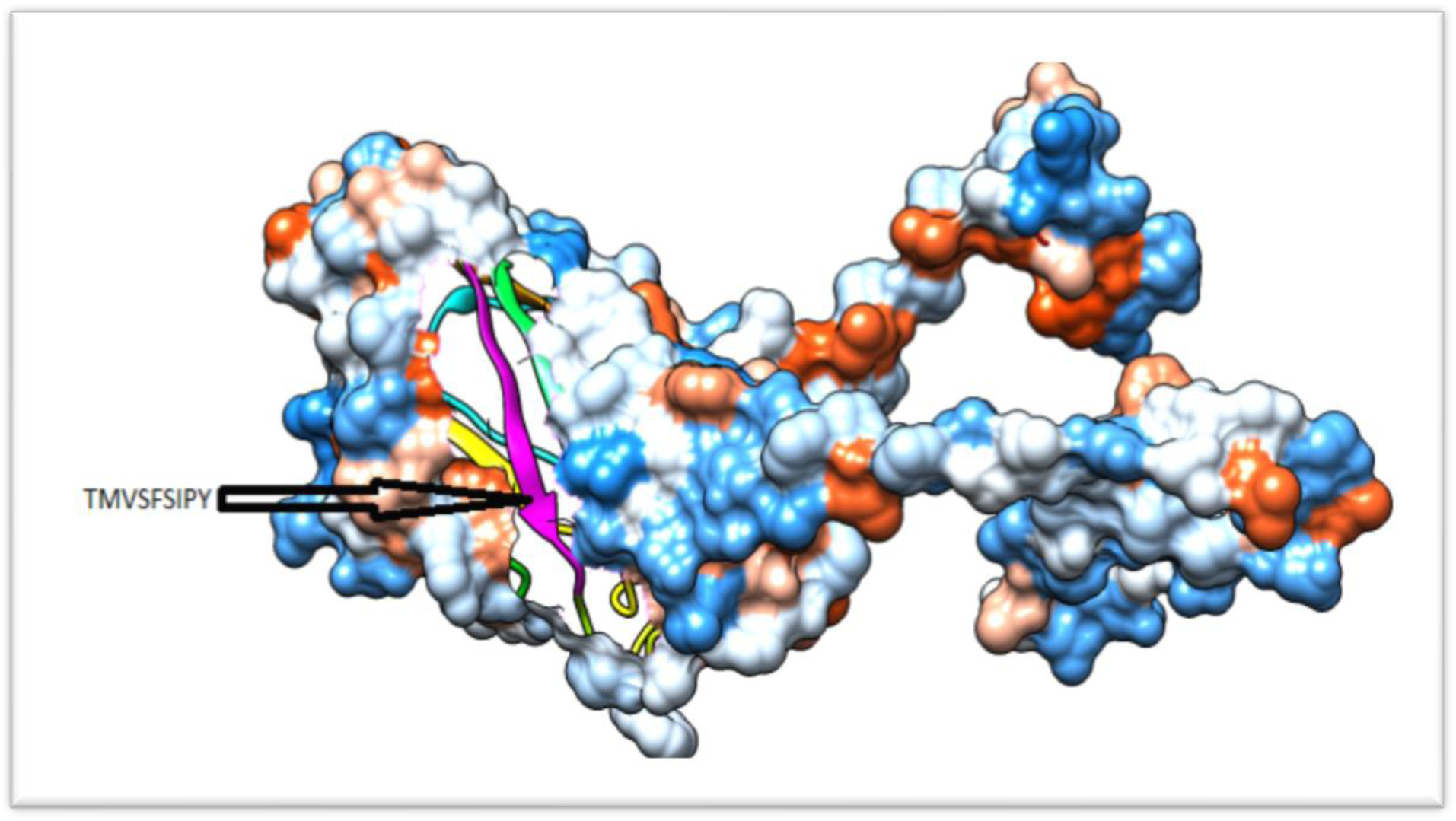
3D structure of cytotoxic T cell epitopes interacts with MCH1.

### 3.3 Prediction of T-cell epitopes and interaction with MHC 11

The (vp1) reference sequence protein of Aichi virus was submitted in the IEDB MHC-11 binding prediction tool to predict epitopes interact with MHC-11 alleles.

**Table (4):**
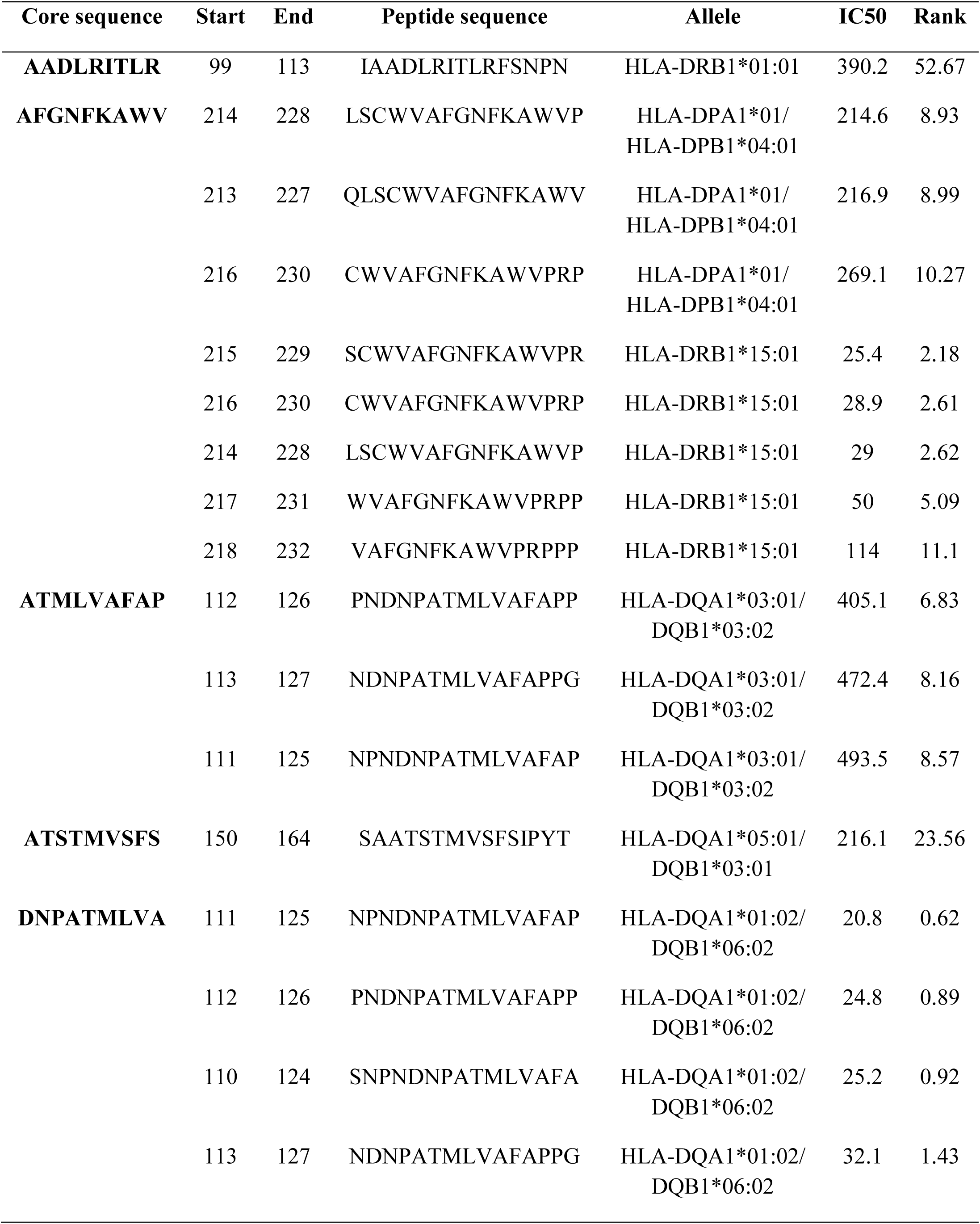
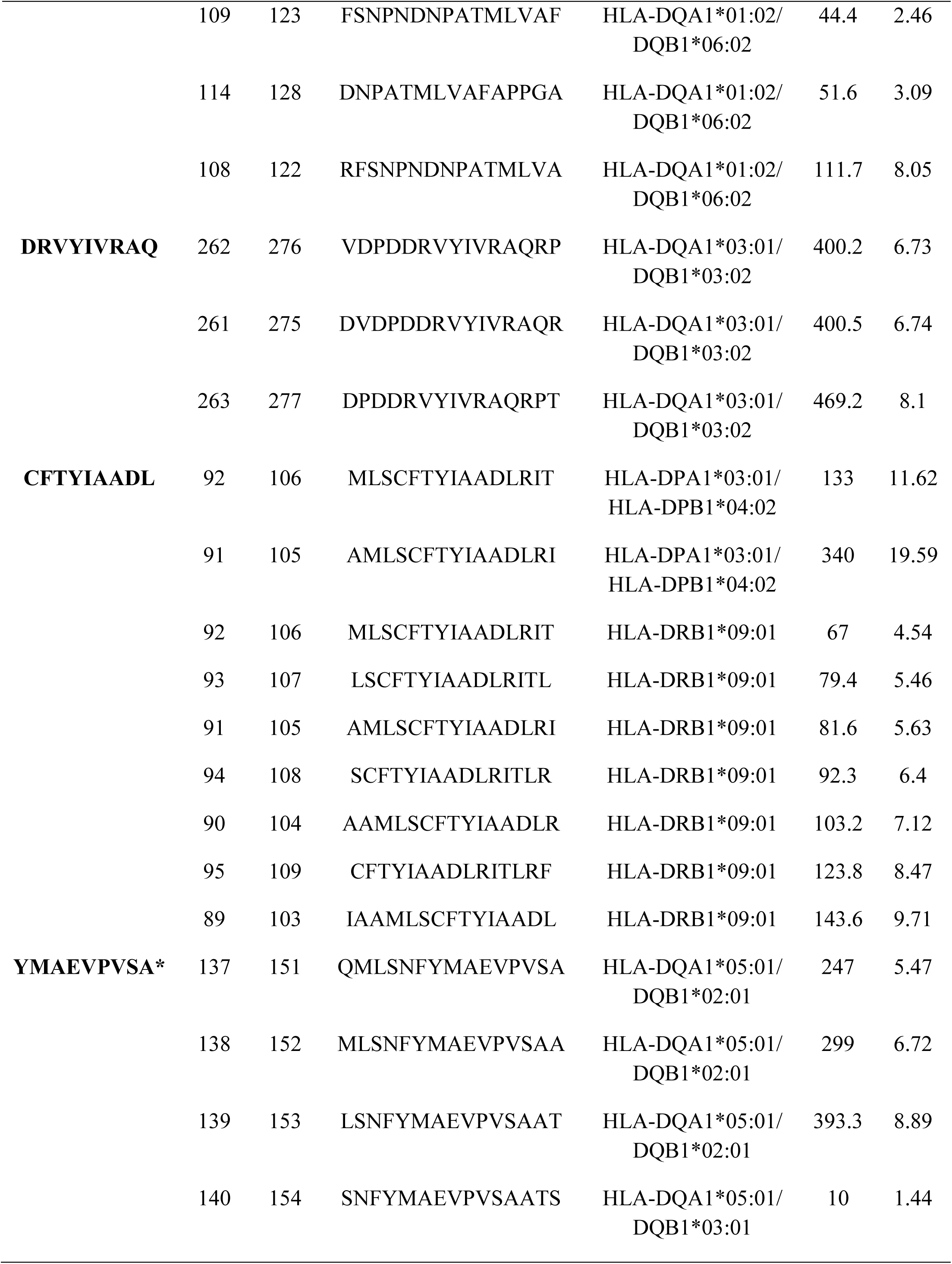
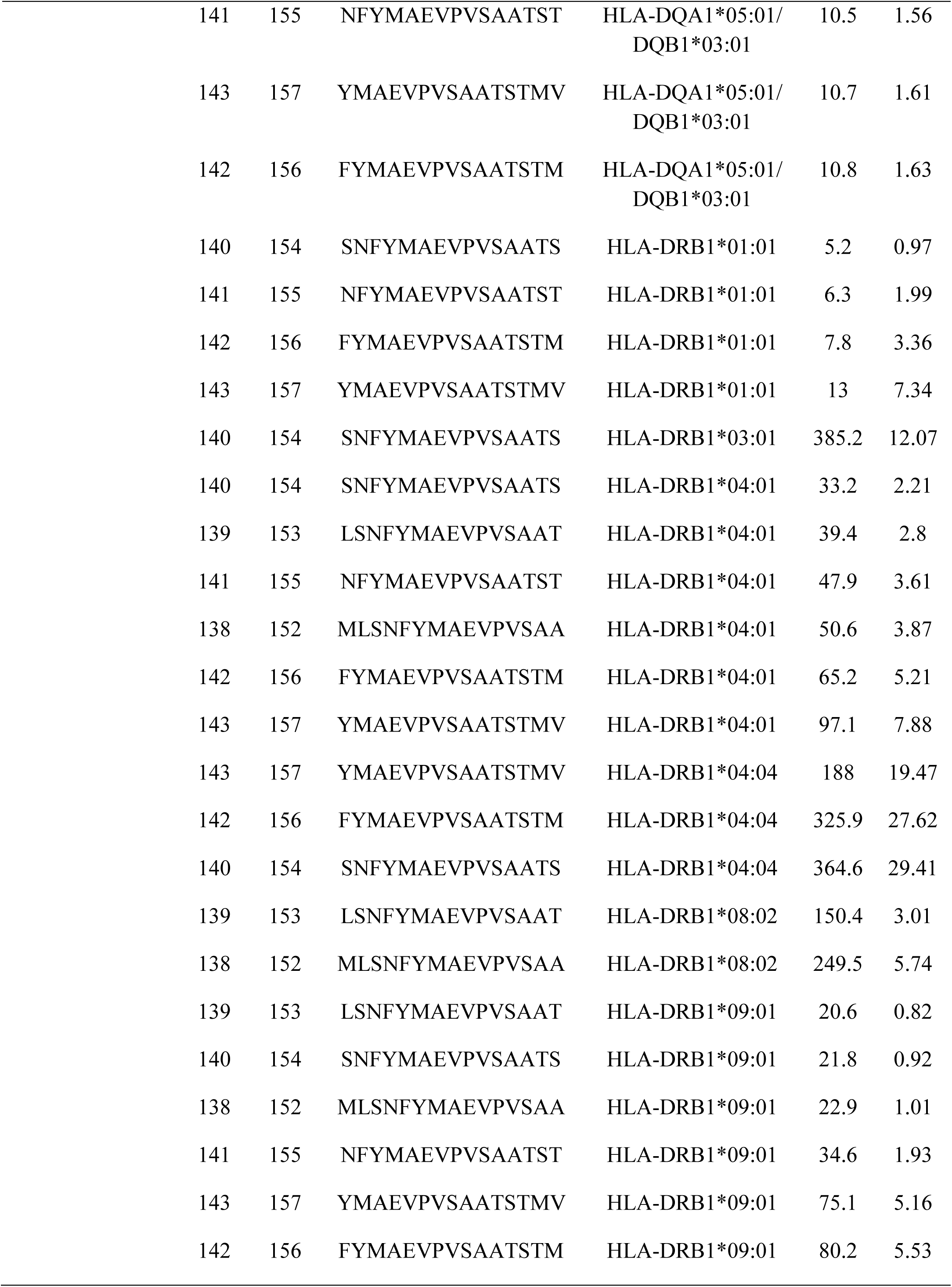
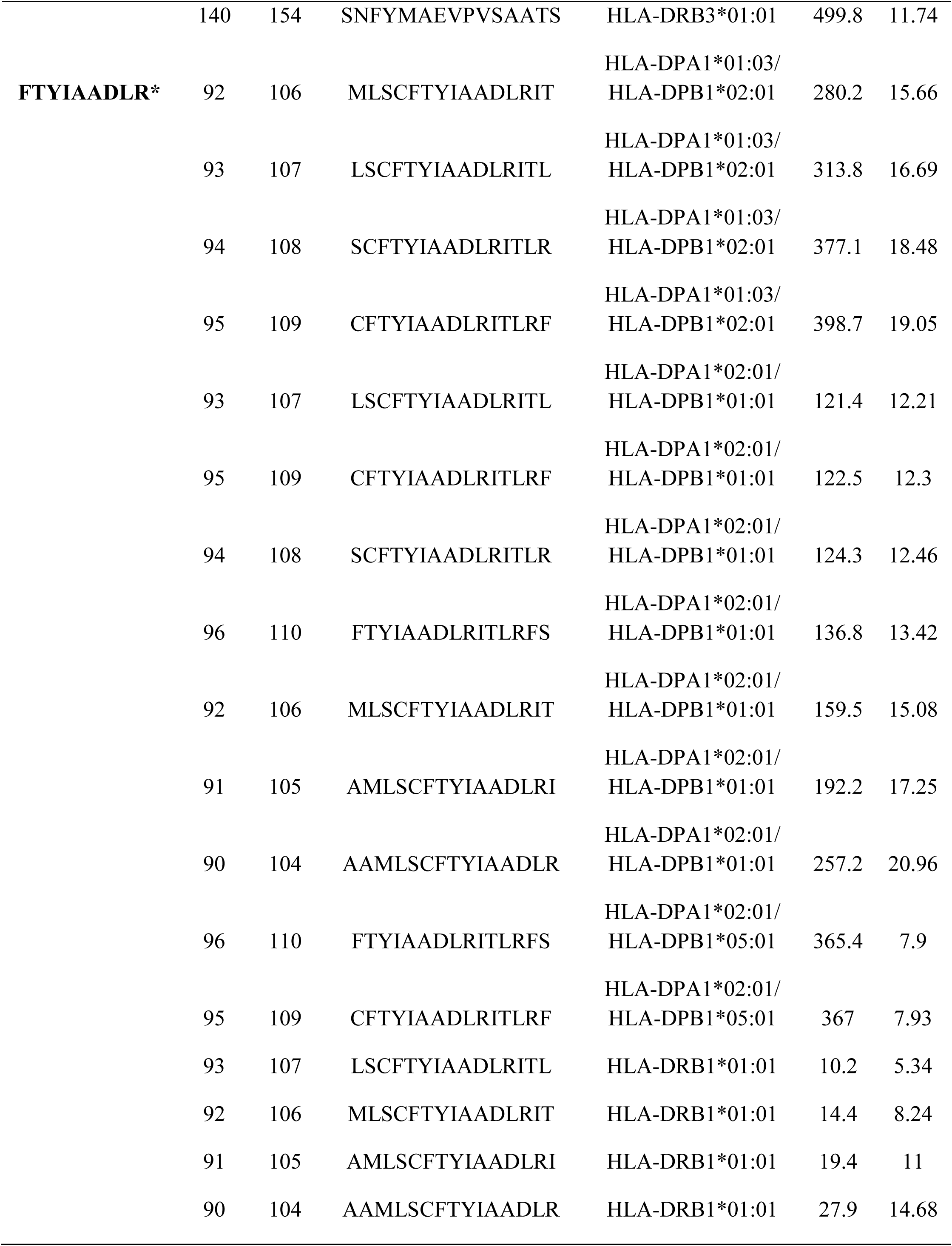
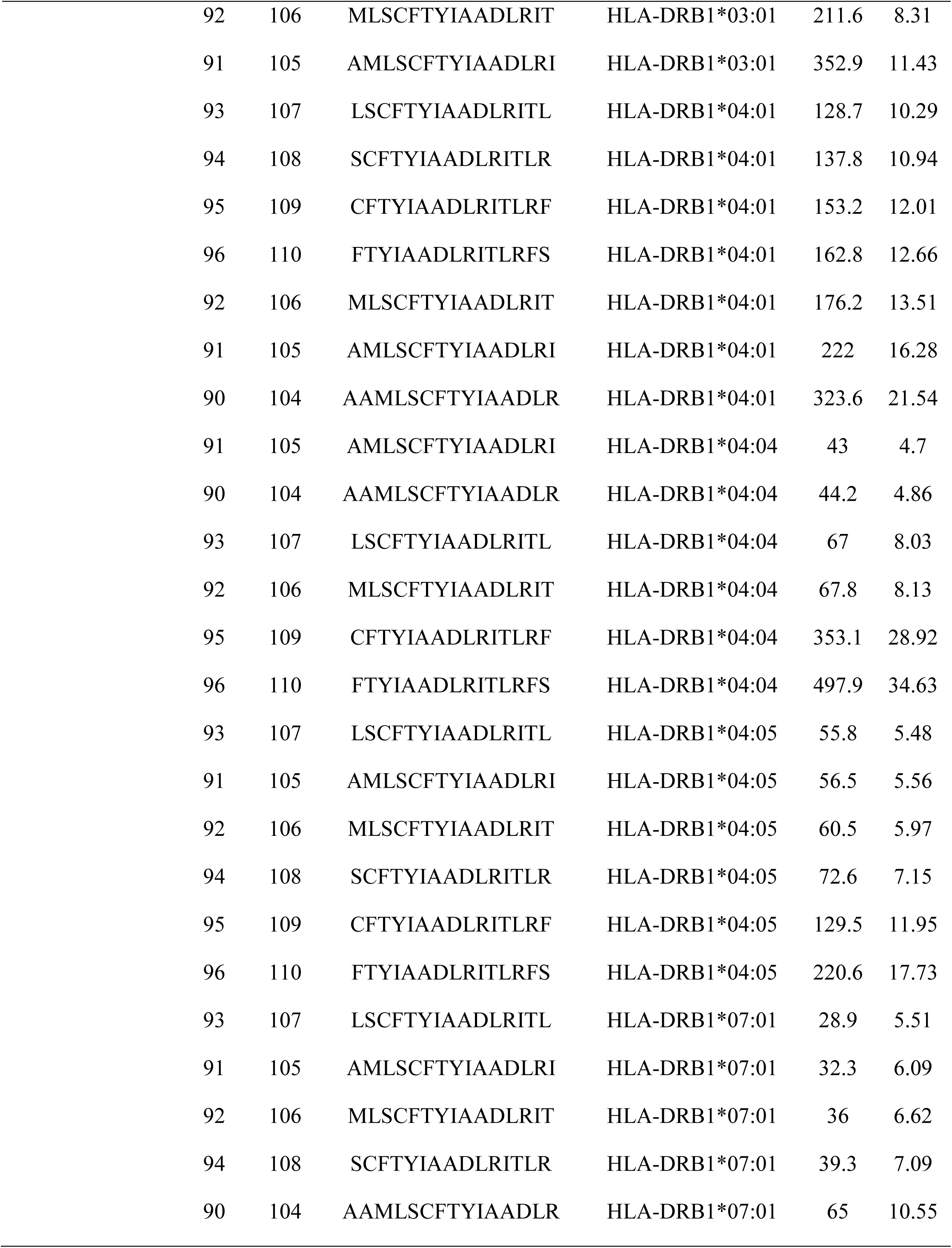
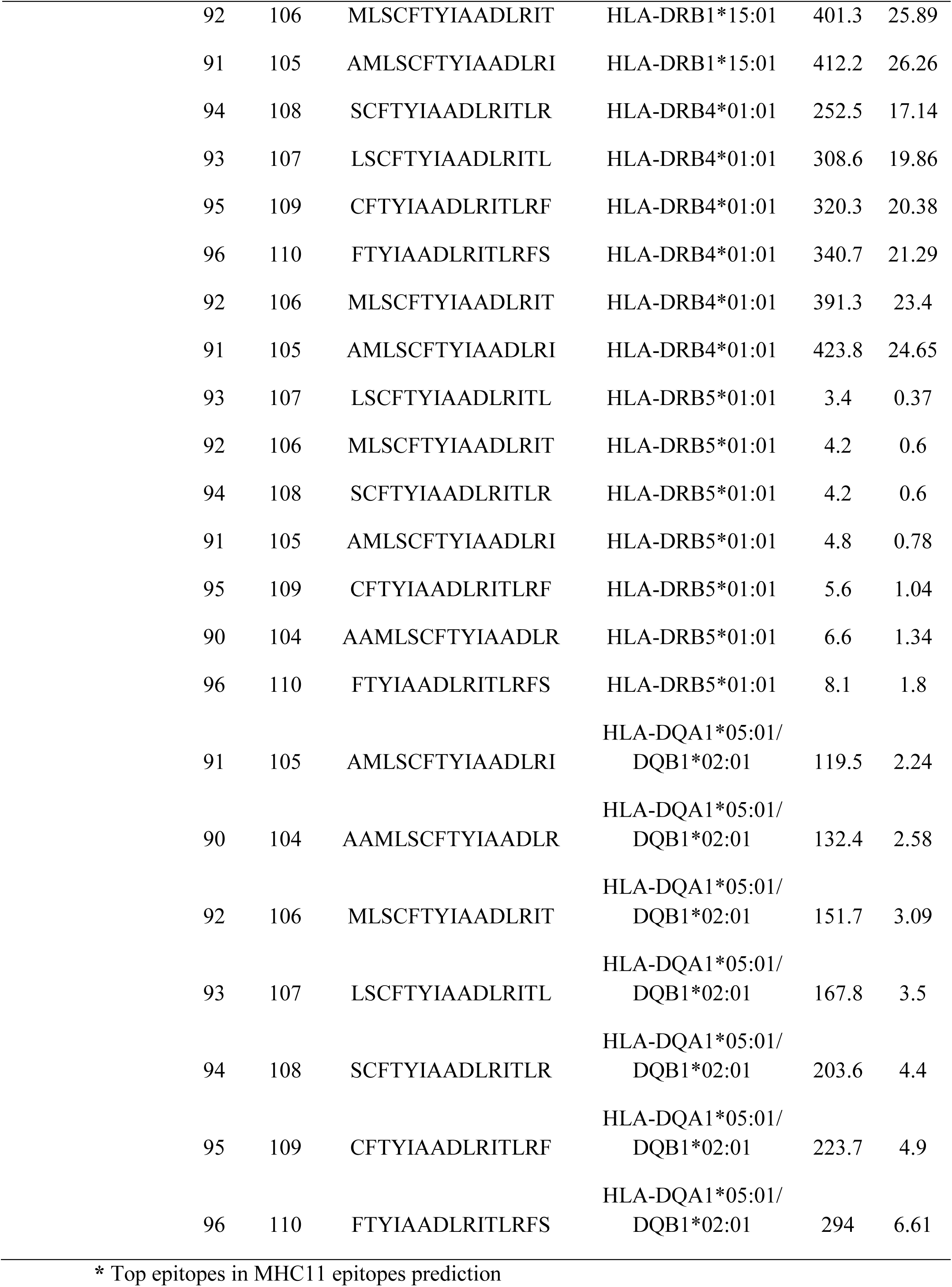
Some of the results of predicted peptides that interact with MHC 11; remaining data as an extra file.

**Figure (10):**
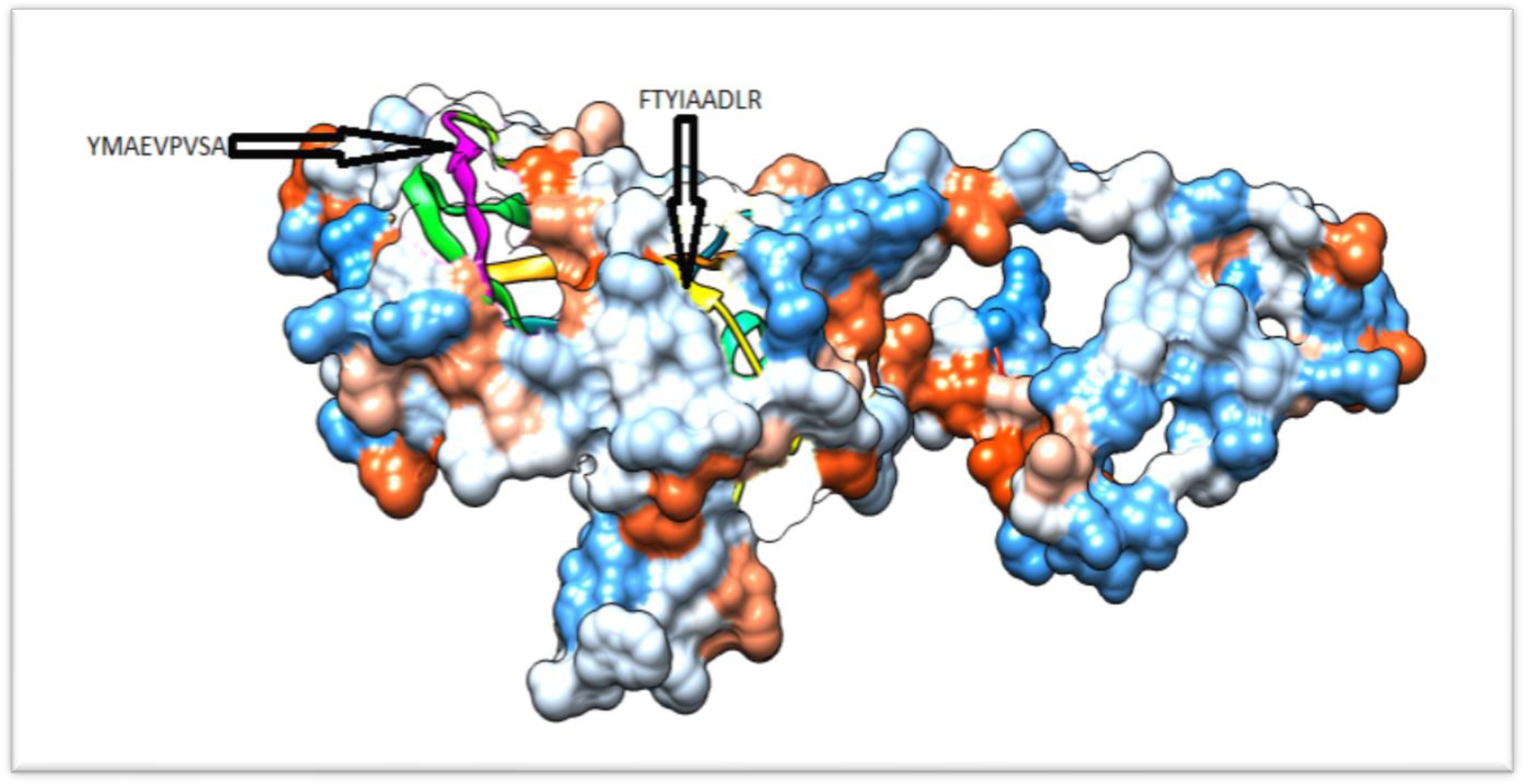
3D structure of T cell top epitopes interacts with MCH1, using chimera.

**Table (5):**
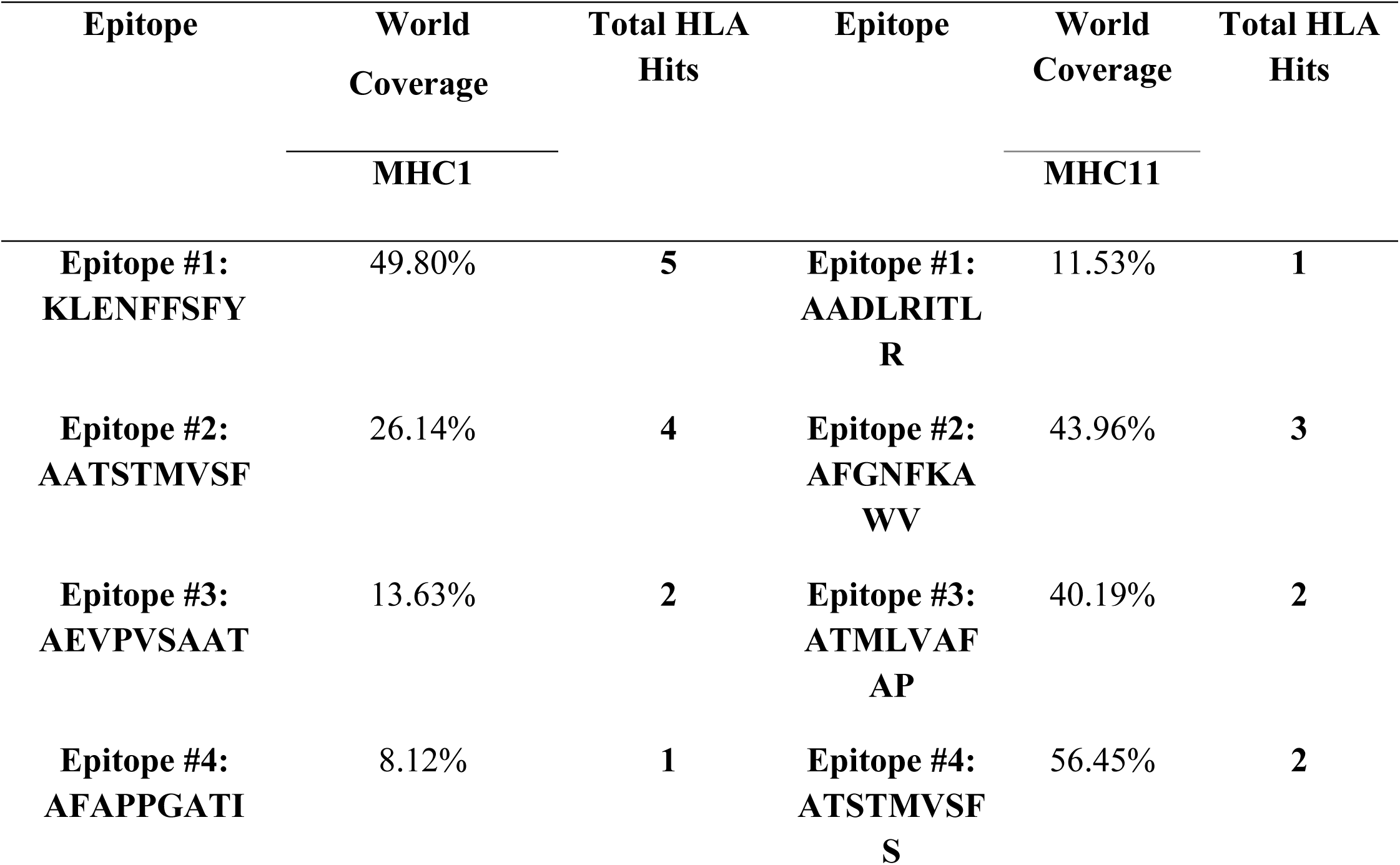
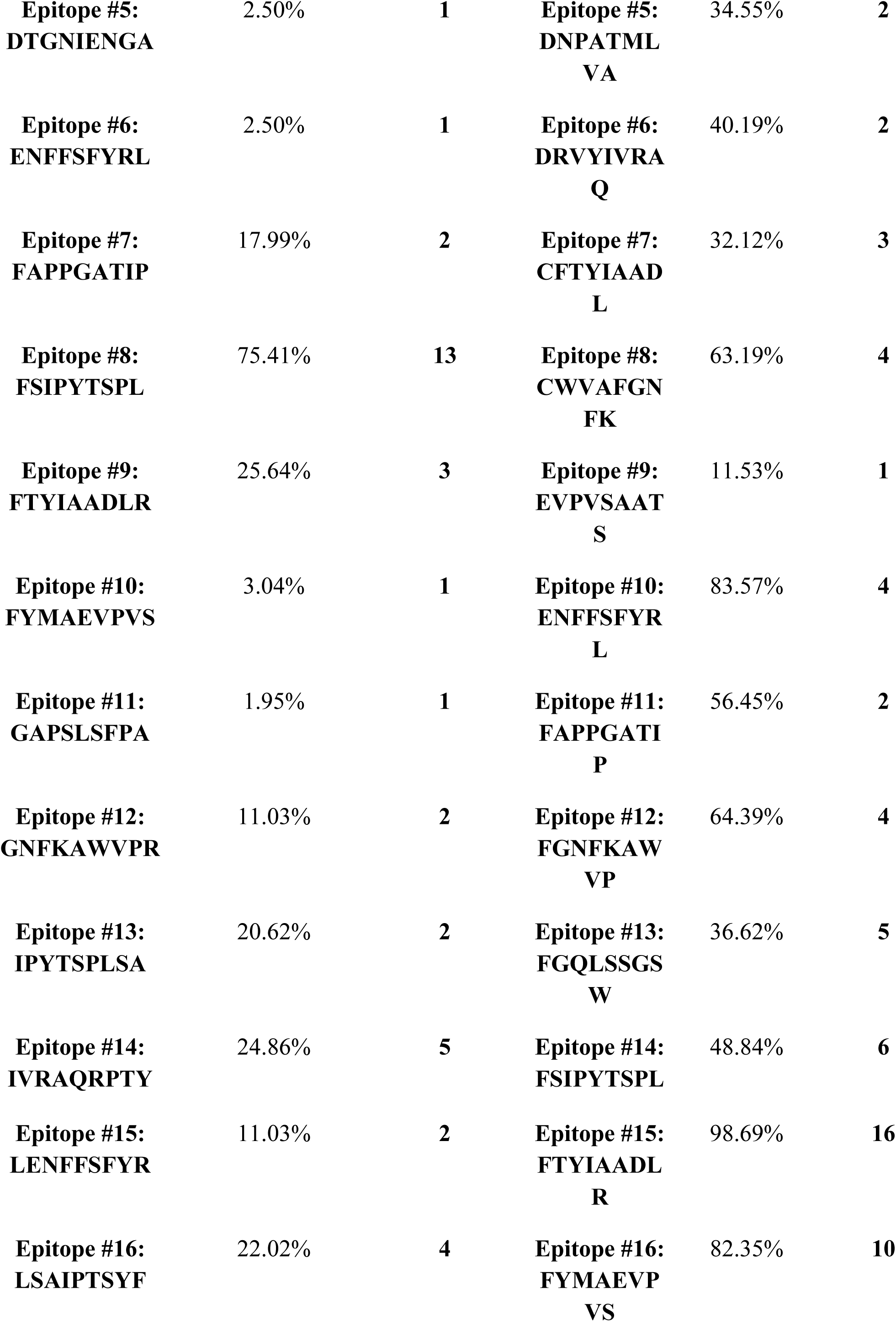
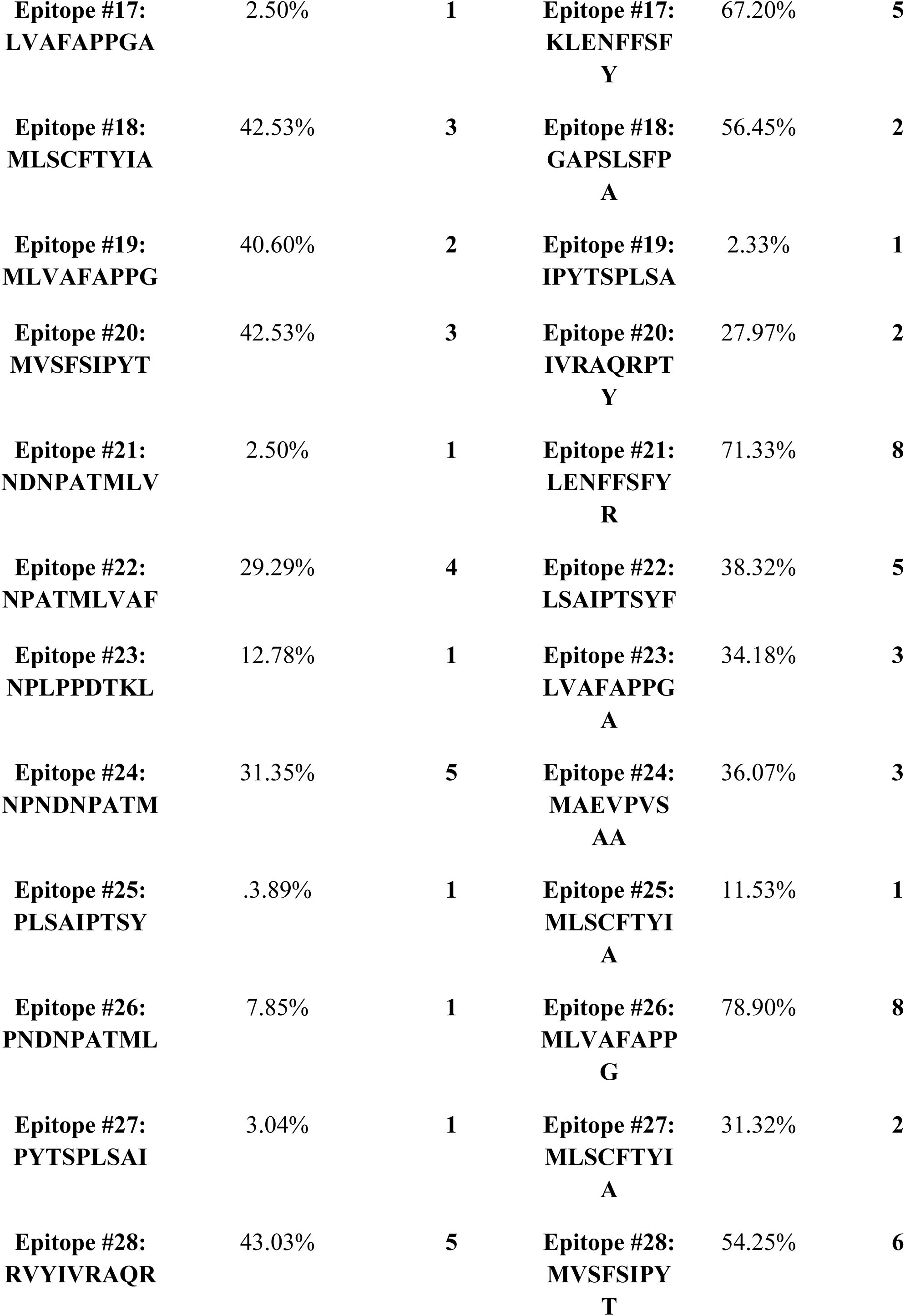
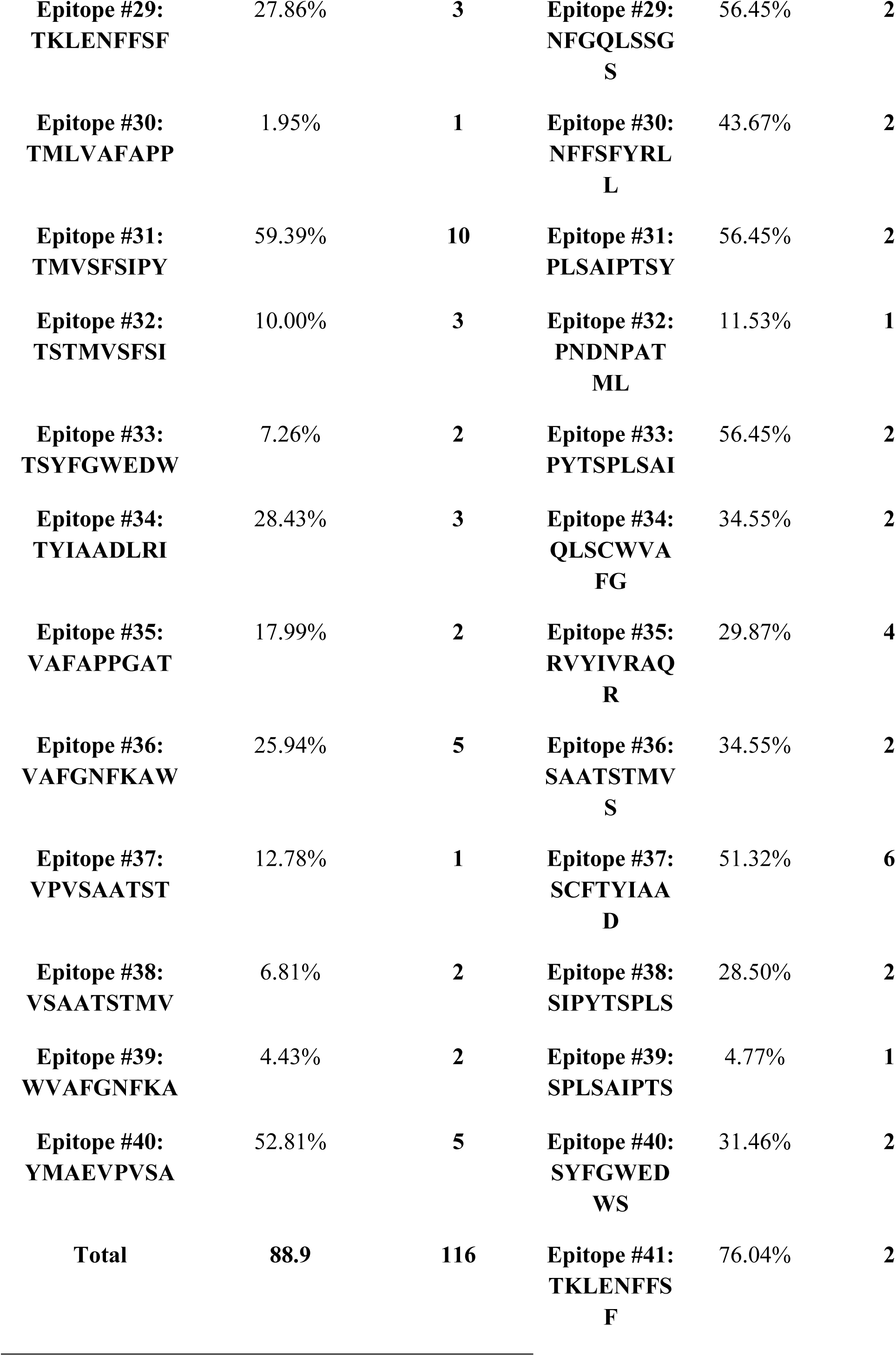
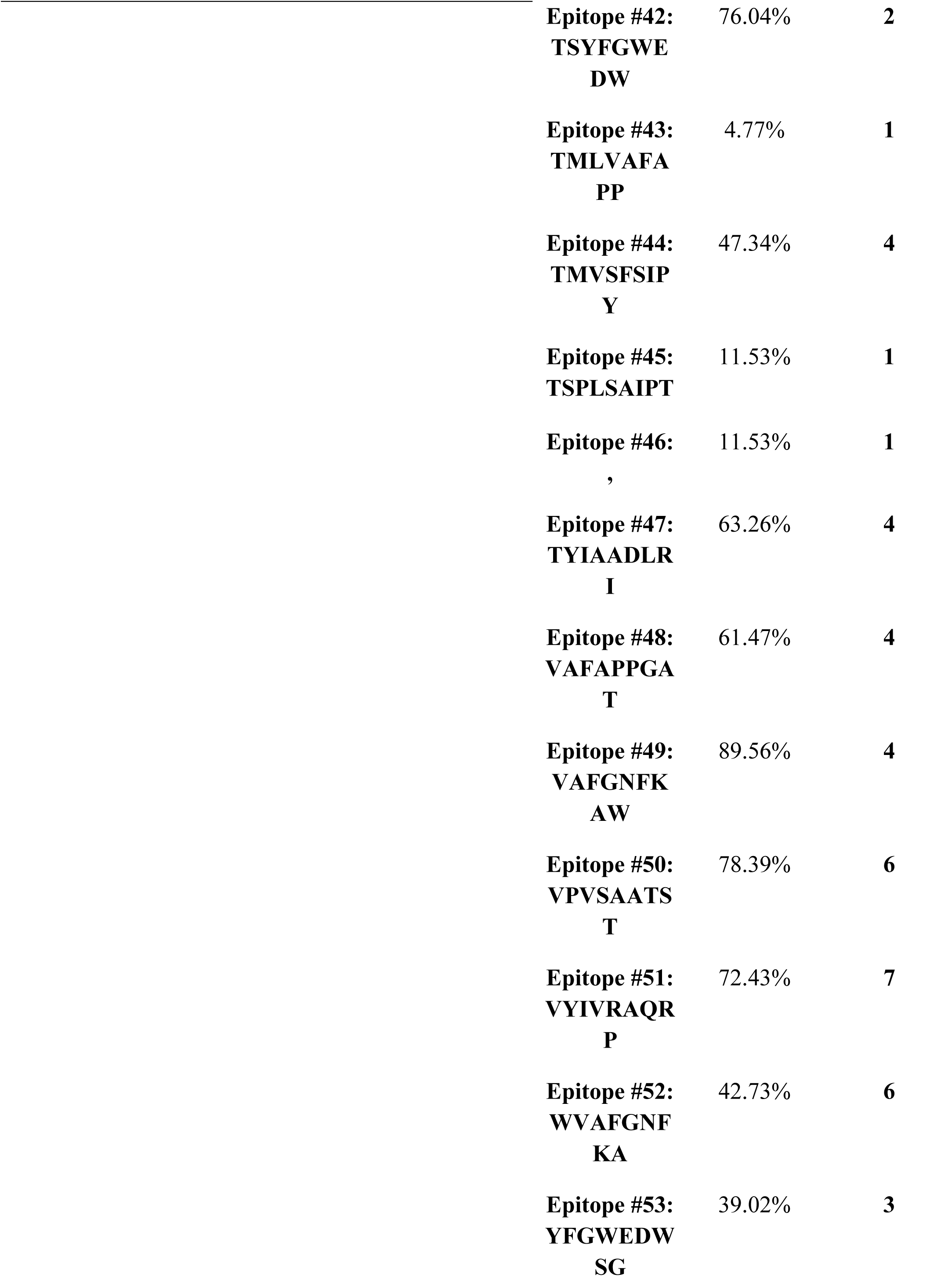
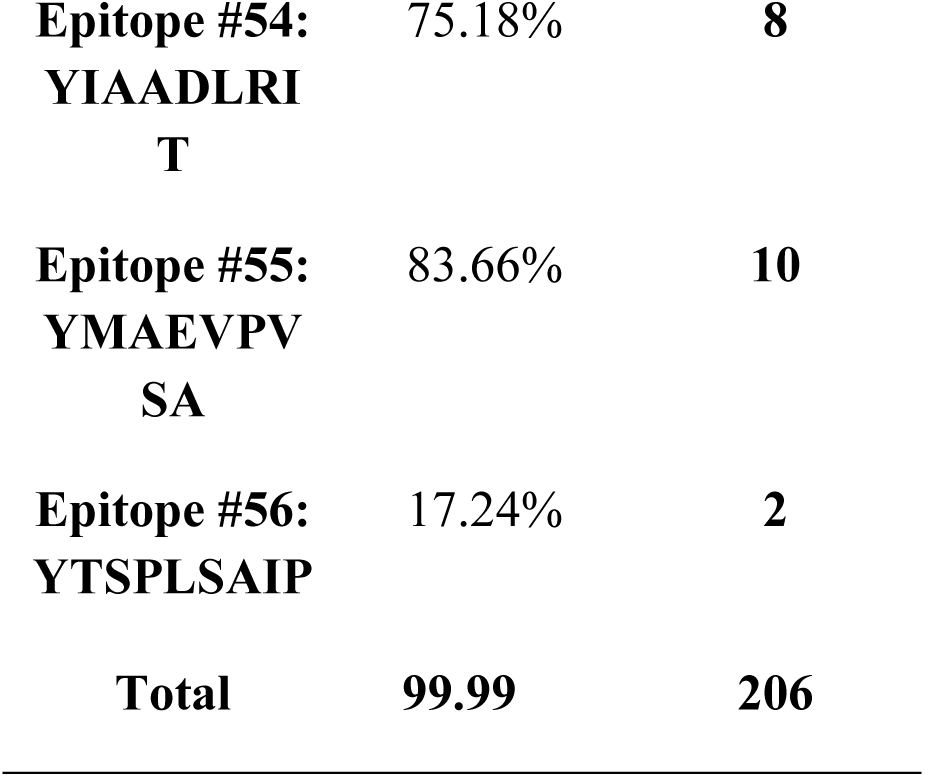
results of population coverage of all peptides in both MHC I and MHC II in the world:

**Table (6):**
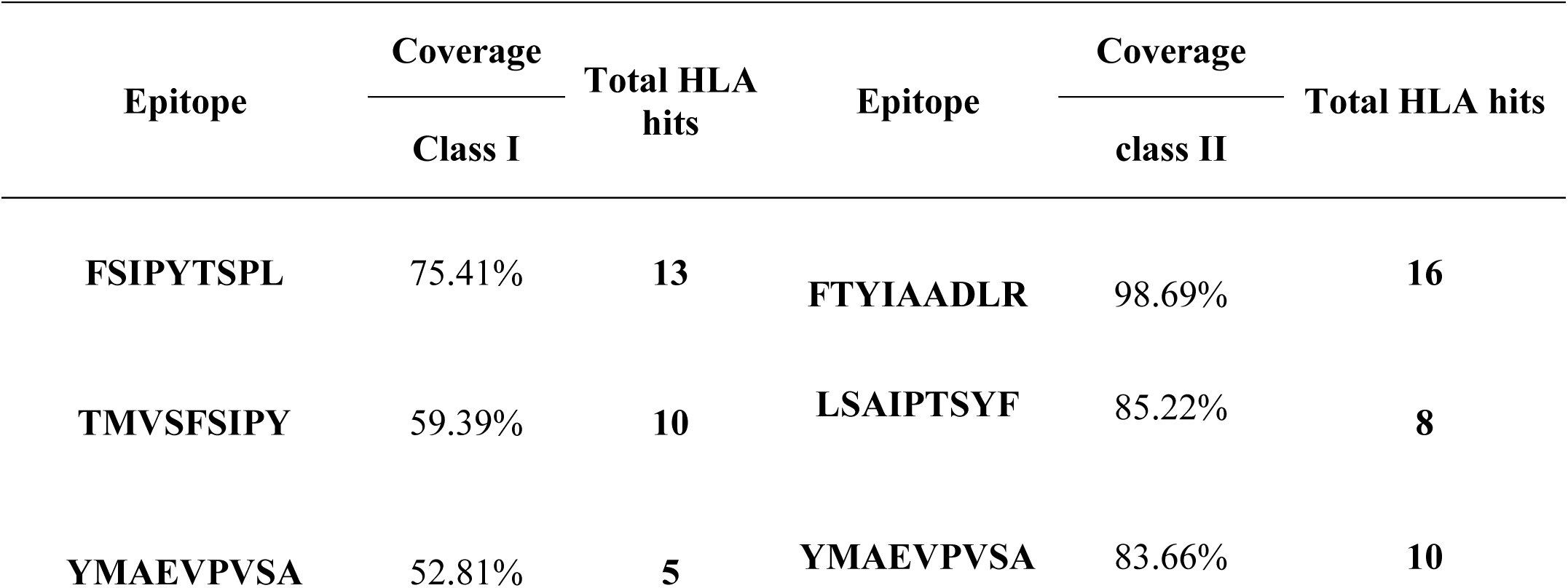
population coverage of top epitopes in both MHC I and MHC II in the world:

## 4. Discussion

In the current study, an immunoinformatic-driven approach used to screen emergent immunogen against Aichi virus. B-cell immunity is given the priority to design vaccine but T-cell was also shown to induce strong immune response (36) According to the prediction result of IEDB the peptides (PLPPDT, PPLPTP, and LPPLPTP) were passed Bepipred linear epitope prediction test, Emini surface accessibility test and Kolaskar and Tongaonkar antigenicity test, there were 14 conserved epitopes that have the binding affinity to B cell while there are 17 epitopes from different windows size was predicted to be on the surface and antigenic according to Emini surface accessibility and Tongaonkar antigenicity test. According to T cell, epitopes are typically peptide fragments and their responses are exquisitely antigen-specific, and they are important as antibodies in defending against infection (37, 38).

T cell immune response is long-lasting immunity as foreign particles and can avoid the effect of memory produced via an immune system. In the prediction results of IEDB, the peptides that have good affinity with HLA molecules were FSIPYTSPL and TMVSFSIPY for MHC1. FTYIAADLR and YMAEVPVSA for MHC class II.

We installed threshold associated with all epitopes in both MHC1 and MHC11 by reformulating the peptides bind with an IC50 value below 500 nM, this allowed computing the number of true negatives, true positives, false negatives, and false positives.

MHC1 binding prediction was analyzed using IEDB Based on Artificial neural network (ANN) with half-maximal inhibitory concentration (IC50) ≤ 500; 41 conserved peptides were predicted to interact with different MHC-1 alleles. The peptide FSIPYTSPL from 159 to 167 had higher affinity to interact with 13 alleles (HLA-A*02:01, HLA-A*02:06, HLA-A*68:02, HLA-B*15:01, HLA-B*35:01, HLA-B*39:01, HLA-B*46:01, HLA-B*58:01, HLA-C*03:03, HLA-C*07:02, HLA-C*12:03, HLA-C*14:02, HLA-C*15:02),followed by TMVSFSIPY from155 to 163 that binds 10 alleles (HLA-A*03:01, HLA-A*11:01, HLA-A*29:02, HLA-A*30:02, HLA-A*32:01, HLA-A*68:01, HLA-B*15:01, HLA-B*15:02, HLA-B*35:01, HLA-C*12:03). World population coverage results for total epitopes binding to MHC1 alleles was 98.9% and for the most promising peptides (FSIPYTSPL, TMVSFSIPY) was 75.41% and 59.39% respectively. these epitopes would possibly be the best vaccine candidates primarily based on the fact that an epitope needs to be as conservative as possible to provide extensive protection among specific virus isolates. These epitopes were also identified as nontoxic to humans depend on the antigenicity test. All the predicted epitopes were placed on the surface of the viral protein1, representing the accessibility for the entered virus.

MHC II binding prediction was also analyzed based on Artificial neural network (NN-align) with half-maximal inhibitory concentration (IC50) ≤ 500 .54 conserved epitopes found to interact with MHC-II alleles. The most promising peptides was FTYIAADLR from 96 to 104 with 9-mers which interact with 16 alleles (HLA-DRB5*01:01, HLA-DRB4*01:01, HLA-DRB1*15:01, HLA-DRB1*07:01, HLA-DRB1*04:05, HLA-DRB1*04:04, HLA-DRB1*04:01, HLA-DRB1*03:01, HLA-DRB1*01:01, HLA-DQA1*05:01, HLA-DQB1*02:01, HLA-DPA1*02:01, HLA-DPB1*05:01, HLA-DPA1*02:01, HLA-DPB1*01:01, HLA-DPA1*01:03, HLA-DPB1*02:01) followed by YMAEVPVSA from (143) to (151) which interact with 10 alleles (HLA-DRB1*09:01, HLA-DRB1*08:02, HLA-DRB1*04:04, HLA-DRB1*04:01, HLA-DRB1*03:01, HLA-DRB1*01:01, HLA-DQA1*05:01, HLA-DQB1*03:01, HLA-DQA1*05:01, HLA-DQB1*02:01). The world population coverage results for all epitopes that have binding affinity to MHC11 alleles was 99.99% while world population coverage of the most promising three epitopes FTYIAADLR and YMAEVPVSA was 98.69% and 83.66% respectively.

An overarching approach to gain most protection against viral infections is to design a successful peptide-based vaccine following the identification of essential epitopes by using the immunoinformatic approach combined with an effective adjuvant choice. Computational immunology is now regarded to contribute to vaccine design in the way of computational chemistry contributes to drug design, before the wet lab confirmation, an advance bioinformatics software should be employed to predict these properties (37, 39).

Immunoinformatic focuses mostly on small peptides ranging from 8 to 11residues, just one epitope per protein can be sufficient to create an immune response in the host(40–42). Bioinformatic techniques to search for epitopes are well understood and available, however can sometimes lead to high false positive rates(43). Despite this drawback, epitope predictors are successful of identifying weak or even strong epitope motifs that have been experimentally ignored (44).

With the advent of next-generation sequencing (NGS) methods, an extraordinary wealth of information has become available that requires more-advanced immunoinformatic tools. this has allowed new opportunities for translational applications of epitope prediction, such as epitope-based design of prophylactic and therapeutic vaccines (45).

## 5. Conclusion

World population coverage results for total epitopes binding for both MHC1 and 11 alleles was 99.8% and the most promising T-cell peptides was FSIPYTSPL from 159 to 167 that considered as a unique domain which successfully interacted with both MHC1 and MHC11 alleles together, and it can be binding with 19 distinctive alleles and provided the highest population coverage epitope set (87.42%) this region is probably promising and This peptide should be considered as a viable peptide vaccine for Aichi virus.

## 6. Recommendation

We recommend Further in vitro and in vivo studies to undertake the effectiveness of these predicted epitopes as peptide vaccine. and also, to do further studies in other strains, there will be a possibility to find common conserved promising epitopes for multiple strains. this work considered for further investigation.

## 7. Acknowledgment

This research was supported by Africa city of technology, for whom the authors would like to show their gratitude for their provided insight and expertise that greatly assisted the research.

## Confect of interest

the authors declare that there is no conflict of interest regarding the publication of this paper and the authors declare that they have no competing interests.

## Data availability

All relevant data used to support the findings of this study are included within the manuscript and supplementary information files.

