## supplemental Table (4) for "Multi Epitope Vaccine Prediction Against Aichi Virus using Immunoinformatic Approach"

Table (4)**: Results of predicted peptides that interact with MHC 11.**

| Core sequence | Start | End | Peptide sequence | Allele | IC50 | Rank |
|---|---|---|---|---|---|---|
| **AADLRITLR** | 99 | 113 | IAADLRITLRFSNPN | HLA-DRB1*01:01 | 390.2 | 52.67 |
| **AFGNFKAWV** | 214 | 228 | LSCWVAFGNFKAWVP | HLA-DPA1*01/ HLA-DPB1*04:01 | 214.6 | 8.93 |
| | 213 | 227 | QLSCWVAFGNFKAWV | HLA-DPA1*01/ HLA-DPB1*04:01 | 216.9 | 8.99 |
| | 216 | 230 | CWVAFGNFKAWVPRP | HLA-DPA1*01/ HLA-DPB1*04:01 | 269.1 | 10.27 |
| | 215 | 229 | SCWVAFGNFKAWVPR | HLA-DRB1*15:01 | 25.4 | 2.18 |
| | 216 | 230 | CWVAFGNFKAWVPRP | HLA-DRB1*15:01 | 28.9 | 2.61 |
| | 214 | 228 | LSCWVAFGNFKAWVP | HLA-DRB1*15:01 | 29 | 2.62 |
| | 217 | 231 | WVAFGNFKAWVPRPP | HLA-DRB1*15:01 | 50 | 5.09 |
| | 218 | 232 | VAFGNFKAWVPRPPP | HLA-DRB1*15:01 | 114 | 11.1 |
| **ATMLVAFAP** | 112 | 126 | PNDNPATMLVAFAPP | HLA-DQA1*03:01/ DQB1*03:02 | 405.1 | 6.83 |
| | 113 | 127 | NDNPATMLVAFAPPG | HLA-DQA1*03:01/ DQB1*03:02 | 472.4 | 8.16 |
| | 111 | 125 | NPNDNPATMLVAFAP | HLA-DQA1*03:01/ DQB1*03:02 | 493.5 | 8.57 |
| **ATSTMVSFS** | 150 | 164 | SAATSTMVSFSIPYT | HLA-DQA1*05:01/ DQB1*03:01 | 216.1 | 23.56 |
| **DNPATMLVA** | 111 | 125 | NPNDNPATMLVAFAP | HLA-DQA1*01:02/ DQB1*06:02 | 20.8 | 0.62 |
| | 112 | 126 | PNDNPATMLVAFAPP | HLA-DQA1*01:02/ DQB1*06:02 | 24.8 | 0.89 |
| | 110 | 124 | SNPNDNPATMLVAFA | HLA-DQA1*01:02/ DQB1*06:02 | 25.2 | 0.92 |
| | 113 | 127 | NDNPATMLVAFAPPG | HLA-DQA1*01:02/ DQB1*06:02 | 32.1 | 1.43 |
| | 109 | 123 | FSNPNDNPATMLVAF | HLA-DQA1*01:02/ DQB1*06:02 | 44.4 | 2.46 |
| | 114 | 128 | DNPATMLVAFAPPGA | HLA-DQA1*01:02/ DQB1*06:02 | 51.6 | 3.09 |
| | 108 | 122 | RFSNPNDNPATMLVA | HLA-DQA1*01:02/ DQB1*06:02 | 111.7 | 8.05 |
| **DRVYIVRAQ** | 262 | 276 | VDPDDRVYIVRAQRP | HLA-DQA1*03:01/ DQB1*03:02 | 400.2 | 6.73 |
| | 261 | 275 | DVDPDDRVYIVRAQR | HLA-DQA1*03:01/ DQB1*03:02 | 400.5 | 6.74 |
| | 263 | 277 | DPDDRVYIVRAQRPT | HLA-DQA1*03:01/ DQB1*03:02 | 469.2 | 8.1 |
| **CFTYIAADL** | 92 | 106 | MLSCFTYIAADLRIT | HLA-DPA1*03:01/ HLA-DPB1*04:02 | 133 | 11.62 |
| | 91 | 105 | AMLSCFTYIAADLRI | HLA-DPA1*03:01/ HLA-DPB1*04:02 | 340 | 19.59 |
| | 92 | 106 | MLSCFTYIAADLRIT | HLA-DRB1*09:01 | 67 | 4.54 |
| | 93 | 107 | LSCFTYIAADLRITL | HLA-DRB1*09:01 | 79.4 | 5.46 |
| | 91 | 105 | AMLSCFTYIAADLRI | HLA-DRB1*09:01 | 81.6 | 5.63 |
| | 94 | 108 | SCFTYIAADLRITLR | HLA-DRB1*09:01 | 92.3 | 6.4 |
| | 90 | 104 | AAMLSCFTYIAADLR | HLA-DRB1*09:01 | 103.2 | 7.12 |
| | 95 | 109 | CFTYIAADLRITLRF | HLA-DRB1*09:01 | 123.8 | 8.47 |
| | 89 | 103 | IAAMLSCFTYIAADL | HLA-DRB1*09:01 | 143.6 | 9.71 |
| **CWVAFGNFK** | 212 | 226 | FQLSCWVAFGNFKAW | HLA-DPA1*01/ HLA-DPB1*04:01 | 168.7 | 7.62 |
| | 215 | 229 | SCWVAFGNFKAWVPR | HLA-DPA1*01/ HLA-DPB1*04:01 | 209 | 8.78 |
| | 213 | 227 | QLSCWVAFGNFKAWV | HLA-DPA1*02:01/ HLA-DPB1*05:01 | 498.1 | 10.34 |
| **EVPVSAATS** | 144 | 158 | MAEVPVSAATSTMVS | HLA-DRB1*01:01 | 64.6 | 24.41 |
| **ENFFSFYRL** | 38 | 52 | PPDTKLENFFSFYRL | HLA-DPA1*01:03/ HLA-DPB1*02:01 | 54.2 | 5.45 |

| | | | | | | |
|---|---|---|---|---|---|---|
| | 40 | 54 | DTKLENFFSFYRLLP | HLA-DQA1*01:01/ DQB1*05:01 | 268.5 | 5.57 |
| | 39 | 53 | PDTKLENFFSFYRLL | HLA-DQA1*01:01/ DQB1*05:01 | 458.8 | 8.49 |
| **FAPPGATIP** | 122 | 136 | AFAPPGATIPLKPTR | HLA-DQA1*05:01/ DQB1*03:01 | 76.3 | 12.34 |
| | 123 | 137 | FAPPGATIPLKPTRQ | HLA-DQA1*05:01/ DQB1*03:01 | 84.7 | 13.29 |
| **FGNFKAWVP** | 217 | 231 | WVAFGNFKAWVPRPP | HLA-DQA1*05:01/ DQB1*03:01 | 157.7 | 19.7 |
| | 218 | 232 | VAFGNFKAWVPRPPP | HLA-DQA1*05:01/ DQB1*03:01 | 189.7 | 21.91 |
| | 216 | 230 | CWVAFGNFKAWVPRP | HLA-DQA1*05:01/ DQB1*03:01 | 198 | 22.44 |
| | 215 | 229 | SCWVAFGNFKAWVPR | HLA-DQA1*05:01/ DQB1*03:01 | 212.9 | 23.38 |
| | 219 | 233 | AFGNFKAWVPRPPPP | HLA-DQA1*05:01/ DQB1*03:01 | 263.9 | 26.2 |
| | 214 | 228 | LSCWVAFGNFKAWVP | HLA-DRB1*07:01 | 261.3 | 24.52 |
| | 215 | 229 | SCWVAFGNFKAWVPR | HLA-DRB1*07:01 | 369.7 | 29.03 |
| | 217 | 231 | WVAFGNFKAWVPRPP | HLA-DRB5*01:01 | 233.7 | 23.71 |
| **FGQLSSGSW** | 182 | 196 | GTNFGQLSSGSWGNL | HLA-DRB1*01:01 | 60.2 | 23.53 |
| | 183 | 197 | TNFGQLSSGSWGNLM | HLA-DRB1*01:01 | 100.2 | 30.21 |
| | 184 | 198 | NFGQLSSGSWGNLML | HLA-DRB1*01:01 | 190.8 | 40.1 |
| | 181 | 195 | SGTNFGQLSSGSWGN | HLA-DRB1*01:01 | 193.1 | 40.3 |
| | 185 | 199 | FGQLSSGSWGNLMLI | HLA-DRB1*01:01 | 210.9 | 41.81 |
| | 180 | 194 | WSGTNFGQLSSGSWG | HLA-DRB1*01:01 | 336.5 | 50.14 |
| | 181 | 195 | SGTNFGQLSSGSWGN | HLA-DRB1*04:01 | 172.5 | 13.28 |
| | 182 | 196 | GTNFGQLSSGSWGNL | HLA-DRB1*04:01 | 176.4 | 13.52 |
| | 180 | 194 | WSGTNFGQLSSGSWG | HLA-DRB1*04:01 | 215.4 | 15.89 |
| | 179 | 193 | DWSGTNFGQLSSGSW | HLA-DRB1*04:01 | 240.1 | 17.28 |
| | 183 | 197 | TNFGQLSSGSWGNLM | HLA-DRB1*04:01 | 266.2 | 18.68 |
| | 184 | 198 | NFGQLSSGSWGNLML | HLA-DRB1*04:01 | 421.1 | 25.78 |
| | 181 | 195 | SGTNFGQLSSGSWGN | HLA-DRB1*09:01 | 62.7 | 4.23 |
| | 183 | 197 | TNFGQLSSGSWGNLM | HLA-DRB1*09:01 | 67.4 | 4.56 |
| | 180 | 194 | WSGTNFGQLSSGSWG | HLA-DRB1*09:01 | 80.8 | 5.57 |
| | 179 | 193 | DWSGTNFGQLSSGSW | HLA-DRB1*09:01 | 89.6 | 6.22 |
| | 184 | 198 | NFGQLSSGSWGNLML | HLA-DRB1*09:01 | 103.5 | 7.14 |
| | 185 | 199 | FGQLSSGSWGNLMLI | HLA-DRB1*09:01 | 156.7 | 10.48 |
| | 182 | 196 | GTNFGQLSSGSWGNL | HLA-DRB1*11:01 | 351.2 | 26.61 |
| | 182 | 196 | GTNFGQLSSGSWGNL | HLA-DRB5*01:01 | 100.9 | 15.37 |
| | 179 | 193 | DWSGTNFGQLSSGSW | HLA-DRB5*01:01 | 107.4 | 15.92 |
| | 181 | 195 | SGTNFGQLSSGSWGN | HLA-DRB5*01:01 | 120.9 | 16.97 |
| | 180 | 194 | WSGTNFGQLSSGSWG | HLA-DRB5*01:01 | 122.1 | 17.06 |
| | 183 | 197 | TNFGQLSSGSWGNLM | HLA-DRB5*01:01 | 198.8 | 21.91 |
| | 184 | 198 | NFGQLSSGSWGNLML | HLA-DRB5*01:01 | 349.6 | 28.42 |
| **FSIPYTSPL** | 156 | 170 | MVSFSIPYTSPLSAI | HLA-DRB1*01:01 | 28.7 | 14.99 |
| | 155 | 169 | TMVSFSIPYTSPLSA | HLA-DRB1*01:01 | 71.9 | 25.77 |
| | 157 | 171 | VSFSIPYTSPLSAIP | HLA-DRB1*01:01 | 74.3 | 26.2 |
| | 154 | 168 | STMVSFSIPYTSPLS | HLA-DRB1*01:01 | 119.2 | 32.7 |
| | 156 | 170 | MVSFSIPYTSPLSAI | HLA-DRB1*04:01 | 175.5 | 13.47 |
| | 156 | 170 | MVSFSIPYTSPLSAI | HLA-DRB1*04:04 | 43.5 | 4.77 |

| | | | | | | |
|---|---|---|---|---|---|---|
| | 159 | 173 | FSIPYTSPLSAIPTS | HLA-DRB1*04:04 | 78.9 | 9.51 |
| | 158 | 172 | SFSIPYTSPLSAIPT | HLA-DRB1*04:04 | 83.3 | 10.03 |
| | 157 | 171 | VSFSIPYTSPLSAIP | HLA-DRB1*04:04 | 103.9 | 12.27 |
| | 154 | 168 | STMVSFSIPYTSPLS | HLA-DRB1*04:05 | 451 | 27.66 |
| | 153 | 167 | TSTMVSFSIPYTSPL | HLA-DRB1*07:01 | 32.4 | 6.11 |
| | 154 | 168 | STMVSFSIPYTSPLS | HLA-DRB1*07:01 | 36.5 | 6.69 |
| | 155 | 169 | TMVSFSIPYTSPLSA | HLA-DRB1*07:01 | 39.9 | 7.18 |
| | 156 | 170 | MVSFSIPYTSPLSAI | HLA-DRB1*07:01 | 45.4 | 8.01 |
| | 157 | 171 | VSFSIPYTSPLSAIP | HLA-DRB1*07:01 | 65.9 | 10.66 |
| | 158 | 172 | SFSIPYTSPLSAIPT | HLA-DRB1*07:01 | 94.5 | 13.66 |
| | 159 | 173 | FSIPYTSPLSAIPTS | HLA-DRB1*07:01 | 137.3 | 17.29 |
| | 157 | 171 | VSFSIPYTSPLSAIP | HLA-DRB1*09:01 | 81.2 | 5.6 |
| | 158 | 172 | SFSIPYTSPLSAIPT | HLA-DRB1*09:01 | 82.5 | 5.7 |
| | 156 | 170 | MVSFSIPYTSPLSAI | HLA-DRB1*09:01 | 88.3 | 6.13 |
| | 155 | 169 | TMVSFSIPYTSPLSA | HLA-DRB1*09:01 | 109.7 | 7.55 |
| | 154 | 168 | STMVSFSIPYTSPLS | HLA-DRB1*09:01 | 128.2 | 8.74 |
| | 153 | 167 | TSTMVSFSIPYTSPL | HLA-DRB1*09:01 | 161.8 | 10.77 |
| **FTYIAADLR** | 92 | 106 | MLSCFTYIAADLRIT | HLA-DPA1*01:03/ HLA-DPB1*02:01 | 280.2 | 15.66 |
| | 93 | 107 | LSCFTYIAADLRITL | HLA-DPA1*01:03/ HLA-DPB1*02:01 | 313.8 | 16.69 |
| | 94 | 108 | SCFTYIAADLRITLR | HLA-DPA1*01:03/ HLA-DPB1*02:01 | 377.1 | 18.48 |
| | 95 | 109 | CFTYIAADLRITLRF | HLA-DPA1*01:03/ HLA-DPB1*02:01 | 398.7 | 19.05 |
| | 93 | 107 | LSCFTYIAADLRITL | HLA-DPA1*02:01/ HLA-DPB1*01:01 | 121.4 | 12.21 |
| | 95 | 109 | CFTYIAADLRITLRF | HLA-DPA1*02:01/ HLA-DPB1*01:01 | 122.5 | 12.3 |
| | 94 | 108 | SCFTYIAADLRITLR | HLA-DPA1*02:01/ HLA-DPB1*01:01 | 124.3 | 12.46 |
| | 96 | 110 | FTYIAADLRITLRFS | HLA-DPA1*02:01/ HLA-DPB1*01:01 | 136.8 | 13.42 |
| | 92 | 106 | MLSCFTYIAADLRIT | HLA-DPA1*02:01/ HLA-DPB1*01:01 | 159.5 | 15.08 |
| | 91 | 105 | AMLSCFTYIAADLRI | HLA-DPA1*02:01/ HLA-DPB1*01:01 | 192.2 | 17.25 |
| | 90 | 104 | AAMLSCFTYIAADLR | HLA-DPA1*02:01/ HLA-DPB1*01:01 | 257.2 | 20.96 |
| | 96 | 110 | FTYIAADLRITLRFS | HLA-DPA1*02:01/ HLA-DPB1*05:01 | 365.4 | 7.9 |
| | 95 | 109 | CFTYIAADLRITLRF | HLA-DPA1*02:01/ HLA-DPB1*05:01 | 367 | 7.93 |
| | 93 | 107 | LSCFTYIAADLRITL | HLA-DRB1*01:01 | 10.2 | 5.34 |
| | 92 | 106 | MLSCFTYIAADLRIT | HLA-DRB1*01:01 | 14.4 | 8.24 |

| | | | | | |
|---|---|---|---|---|---|
| 91 | 105 | AMLSCFTYIAADLRI | HLA-DRB1*01:01 | 19.4 | 11 |
| 90 | 104 | AAMLSCFTYIAADLR | HLA-DRB1*01:01 | 27.9 | 14.68 |
| 92 | 106 | MLSCFTYIAADLRIT | HLA-DRB1*03:01 | 211.6 | 8.31 |
| 91 | 105 | AMLSCFTYIAADLRI | HLA-DRB1*03:01 | 352.9 | 11.43 |
| 93 | 107 | LSCFTYIAADLRITL | HLA-DRB1*04:01 | 128.7 | 10.29 |
| 94 | 108 | SCFTYIAADLRITLR | HLA-DRB1*04:01 | 137.8 | 10.94 |
| 95 | 109 | CFTYIAADLRITLRF | HLA-DRB1*04:01 | 153.2 | 12.01 |
| 96 | 110 | FTYIAADLRITLRFS | HLA-DRB1*04:01 | 162.8 | 12.66 |
| 92 | 106 | MLSCFTYIAADLRIT | HLA-DRB1*04:01 | 176.2 | 13.51 |
| 91 | 105 | AMLSCFTYIAADLRI | HLA-DRB1*04:01 | 222 | 16.28 |
| 90 | 104 | AAMLSCFTYIAADLR | HLA-DRB1*04:01 | 323.6 | 21.54 |
| 91 | 105 | AMLSCFTYIAADLRI | HLA-DRB1*04:04 | 43 | 4.7 |
| 90 | 104 | AAMLSCFTYIAADLR | HLA-DRB1*04:04 | 44.2 | 4.86 |
| 93 | 107 | LSCFTYIAADLRITL | HLA-DRB1*04:04 | 67 | 8.03 |
| 92 | 106 | MLSCFTYIAADLRIT | HLA-DRB1*04:04 | 67.8 | 8.13 |
| 95 | 109 | CFTYIAADLRITLRF | HLA-DRB1*04:04 | 353.1 | 28.92 |
| 96 | 110 | FTYIAADLRITLRFS | HLA-DRB1*04:04 | 497.9 | 34.63 |
| 93 | 107 | LSCFTYIAADLRITL | HLA-DRB1*04:05 | 55.8 | 5.48 |
| 91 | 105 | AMLSCFTYIAADLRI | HLA-DRB1*04:05 | 56.5 | 5.56 |
| 92 | 106 | MLSCFTYIAADLRIT | HLA-DRB1*04:05 | 60.5 | 5.97 |
| 94 | 108 | SCFTYIAADLRITLR | HLA-DRB1*04:05 | 72.6 | 7.15 |
| 95 | 109 | CFTYIAADLRITLRF | HLA-DRB1*04:05 | 129.5 | 11.95 |
| 96 | 110 | FTYIAADLRITLRFS | HLA-DRB1*04:05 | 220.6 | 17.73 |
| 93 | 107 | LSCFTYIAADLRITL | HLA-DRB1*07:01 | 28.9 | 5.51 |
| 91 | 105 | AMLSCFTYIAADLRI | HLA-DRB1*07:01 | 32.3 | 6.09 |
| 92 | 106 | MLSCFTYIAADLRIT | HLA-DRB1*07:01 | 36 | 6.62 |
| 94 | 108 | SCFTYIAADLRITLR | HLA-DRB1*07:01 | 39.3 | 7.09 |
| 90 | 104 | AAMLSCFTYIAADLR | HLA-DRB1*07:01 | 65 | 10.55 |
| 92 | 106 | MLSCFTYIAADLRIT | HLA-DRB1*15:01 | 401.3 | 25.89 |
| 91 | 105 | AMLSCFTYIAADLRI | HLA-DRB1*15:01 | 412.2 | 26.26 |
| 94 | 108 | SCFTYIAADLRITLR | HLA-DRB4*01:01 | 252.5 | 17.14 |
| 93 | 107 | LSCFTYIAADLRITL | HLA-DRB4*01:01 | 308.6 | 19.86 |
| 95 | 109 | CFTYIAADLRITLRF | HLA-DRB4*01:01 | 320.3 | 20.38 |
| 96 | 110 | FTYIAADLRITLRFS | HLA-DRB4*01:01 | 340.7 | 21.29 |
| 92 | 106 | MLSCFTYIAADLRIT | HLA-DRB4*01:01 | 391.3 | 23.4 |
| 91 | 105 | AMLSCFTYIAADLRI | HLA-DRB4*01:01 | 423.8 | 24.65 |
| 93 | 107 | LSCFTYIAADLRITL | HLA-DRB5*01:01 | 3.4 | 0.37 |
| 92 | 106 | MLSCFTYIAADLRIT | HLA-DRB5*01:01 | 4.2 | 0.6 |
| 94 | 108 | SCFTYIAADLRITLR | HLA-DRB5*01:01 | 4.2 | 0.6 |
| 91 | 105 | AMLSCFTYIAADLRI | HLA-DRB5*01:01 | 4.8 | 0.78 |
| 95 | 109 | CFTYIAADLRITLRF | HLA-DRB5*01:01 | 5.6 | 1.04 |
| 90 | 104 | AAMLSCFTYIAADLR | HLA-DRB5*01:01 | 6.6 | 1.34 |
| 96 | 110 | FTYIAADLRITLRFS | HLA-DRB5*01:01 | 8.1 | 1.8 |

| | | | | | | |
|---|---|---|---|---|---|---|
| | 91 | 105 | AMLSCFTYIAADLRI | HLA-DQA1*05:01/ DQB1*02:01 | 119.5 | 2.24 |
| | 90 | 104 | AAMLSCFTYIAADLR | HLA-DQA1*05:01/ DQB1*02:01 | 132.4 | 2.58 |
| | 92 | 106 | MLSCFTYIAADLRIT | HLA-DQA1*05:01/ DQB1*02:01 | 151.7 | 3.09 |
| | 93 | 107 | LSCFTYIAADLRITL | HLA-DQA1*05:01/ DQB1*02:01 | 167.8 | 3.5 |
| | 94 | 108 | SCFTYIAADLRITLR | HLA-DQA1*05:01/ DQB1*02:01 | 203.6 | 4.4 |
| | 95 | 109 | CFTYIAADLRITLRF | HLA-DQA1*05:01/ DQB1*02:01 | 223.7 | 4.9 |
| | 96 | 110 | FTYIAADLRITLRFS | HLA-DQA1*05:01/ DQB1*02:01 | 294 | 6.61 |
| **FYMAEVPVS** | 139 | 153 | LSNFYMAEVPVSAAT | HLA-DQA1*04:01/ DQB1*04:02 | 213 | 2.76 |
| | 140 | 154 | SNFYMAEVPVSAATS | HLA-DQA1*04:01/ DQB1*04:02 | 236.4 | 3.18 |
| | 138 | 152 | MLSNFYMAEVPVSAA | HLA-DQA1*04:01/ DQB1*04:02 | 278.3 | 3.91 |
| | 141 | 155 | NFYMAEVPVSAATST | HLA-DQA1*04:01/ DQB1*04:02 | 311.1 | 4.49 |
| | 136 | 150 | RQMLSNFYMAEVPVS | HLA-DQA1*04:01/ DQB1*04:02 | 317.4 | 4.6 |
| | 137 | 151 | QMLSNFYMAEVPVSA | HLA-DQA1*04:01/ DQB1*04:02 | 321.7 | 4.68 |
| | 142 | 156 | FYMAEVPVSAATSTM | HLA-DQA1*04:01/ DQB1*04:02 | 369 | 5.51 |
| | 139 | 153 | LSNFYMAEVPVSAAT | HLA-DQA1*05:01/ DQB1*03:01 | 11.3 | 1.74 |
| | 138 | 152 | MLSNFYMAEVPVSAA | HLA-DQA1*05:01/ DQB1*03:01 | 13.9 | 2.35 |
| | 137 | 151 | QMLSNFYMAEVPVSA | HLA-DQA1*05:01/ DQB1*03:01 | 15.3 | 2.66 |
| | 136 | 150 | RQMLSNFYMAEVPVS | HLA-DQA1*05:01/ DQB1*03:01 | 62.9 | 10.72 |
| | 139 | 153 | LSNFYMAEVPVSAAT | HLA-DRB1*01:01 | 5 | 0.79 |
| | 138 | 152 | MLSNFYMAEVPVSAA | HLA-DRB1*01:01 | 5.7 | 1.43 |
| | 137 | 151 | QMLSNFYMAEVPVSA | HLA-DRB1*01:01 | 7.3 | 2.91 |
| | 136 | 150 | RQMLSNFYMAEVPVS | HLA-DRB1*01:01 | 10.9 | 5.88 |
| | 137 | 151 | QMLSNFYMAEVPVSA | HLA-DRB1*04:01 | 72.2 | 5.82 |
| | 136 | 150 | RQMLSNFYMAEVPVS | HLA-DRB1*04:01 | 211.1 | 15.66 |
| | 139 | 153 | LSNFYMAEVPVSAAT | HLA-DRB1*04:05 | 37.8 | 3.45 |
| | 138 | 152 | MLSNFYMAEVPVSAA | HLA-DRB1*04:05 | 38.5 | 3.54 |
| | 137 | 151 | QMLSNFYMAEVPVSA | HLA-DRB1*04:05 | 40.2 | 3.73 |
| | 136 | 150 | RQMLSNFYMAEVPVS | HLA-DRB1*04:05 | 44.7 | 4.26 |
| | 140 | 154 | SNFYMAEVPVSAATS | HLA-DRB1*04:05 | 52 | 5.07 |
| | 141 | 155 | NFYMAEVPVSAATST | HLA-DRB1*04:05 | 89.9 | 8.76 |
| | 142 | 156 | FYMAEVPVSAATSTM | HLA-DRB1*04:05 | 151.3 | 13.47 |
| | 136 | 150 | RQMLSNFYMAEVPVS | HLA-DRB1*07:01 | 457.5 | 32.02 |
| | 139 | 153 | LSNFYMAEVPVSAAT | HLA-DRB1*11:01 | 227.9 | 21.68 |
| | 138 | 152 | MLSNFYMAEVPVSAA | HLA-DRB1*11:01 | 331.4 | 25.92 |
| | 140 | 154 | SNFYMAEVPVSAATS | HLA-DRB1*11:01 | 385.8 | 27.77 |
| | 139 | 153 | LSNFYMAEVPVSAAT | HLA-DRB3*01:01 | 349.6 | 9.52 |
| | 138 | 152 | MLSNFYMAEVPVSAA | HLA-DRB3*01:01 | 367.9 | 9.81 |
| | 137 | 151 | QMLSNFYMAEVPVSA | HLA-DRB3*01:01 | 402.3 | 10.34 |
| **KLENFFSFY** | 38 | 52 | PPDTKLENFFSFYRL | HLA-DPA1*01/ HLA-DPB1*04:01 | 106.2 | 5.54 |
| | 37 | 51 | LPPDTKLENFFSFYR | HLA-DPA1*01/ HLA-DPB1*04:01 | 183.6 | 8.08 |
| | 36 | 50 | PLPPDTKLENFFSFY | HLA-DPA1*01/ HLA-DPB1*04:01 | 244.8 | 9.68 |
| | 36 | 50 | PLPPDTKLENFFSFY | HLA-DPA1*02:01/ HLA-DPB1*01:01 | 266.5 | 21.43 |

| | | | | | | |
|---|---|---|---|---|---|---|
| | 40 | 54 | DTKLENFFSFYRLLP | HLA-DPA1*02:01/ HLA-DPB1*05:01 | 26 | 0.25 |
| | 41 | 55 | TKLENFFSFYRLLPM | HLA-DPA1*02:01/ HLA-DPB1*05:01 | 26.8 | 0.26 |
| | 39 | 53 | PDTKLENFFSFYRLL | HLA-DPA1*02:01/ HLA-DPB1*05:01 | 27.9 | 0.28 |
| | 38 | 52 | PPDTKLENFFSFYRL | HLA-DPA1*02:01/ HLA-DPB1*05:01 | 36.5 | 0.45 |
| | 42 | 56 | KLENFFSFYRLLPMG | HLA-DPA1*02:01/ HLA-DPB1*05:01 | 37.3 | 0.47 |
| | 37 | 51 | LPPDTKLENFFSFYR | HLA-DPA1*02:01/ HLA-DPB1*05:01 | 51.9 | 0.8 |
| | 36 | 50 | PLPPDTKLENFFSFY | HLA-DPA1*02:01/ HLA-DPB1*05:01 | 143.7 | 3.09 |
| **GAPSLSFPA** | 57 | 71 | GSGAPSLSFPADEGT | HLA-DQA1*05:01/ DQB1*03:01 | 59.9 | 10.34 |
| | 58 | 72 | SGAPSLSFPADEGTI | HLA-DQA1*05:01/ DQB1*03:01 | 110.4 | 15.84 |
| | 59 | 73 | GAPSLSFPADEGTII | HLA-DQA1*05:01/ DQB1*03:01 | 117.6 | 16.49 |
| **IPYTSPLSA** | 158 | 172 | SFSIPYTSPLSAIPT | HLA-DRB1*08:02 | 194.6 | 4.23 |
| | 156 | 170 | MVSFSIPYTSPLSAI | HLA-DRB1*08:02 | 200.8 | 4.41 |
| | 157 | 171 | VSFSIPYTSPLSAIP | HLA-DRB1*08:02 | 210.1 | 4.65 |
| | 159 | 173 | FSIPYTSPLSAIPTS | HLA-DRB1*08:02 | 253.7 | 5.84 |
| | 160 | 174 | SIPYTSPLSAIPTSY | HLA-DRB1*08:02 | 356.1 | 8.55 |
| **IVRAQRPTY** | 264 | 278 | PDDRVYIVRAQRPTY | HLA-DRB1*03:01 | 263.8 | 9.54 |
| | 264 | 278 | PDDRVYIVRAQRPTY | HLA-DRB1*04:01 | 370.7 | 23.67 |
| **LENFFSFYR** | 39 | 53 | PDTKLENFFSFYRLL | HLA-DPA1*02:01/ HLA-DPB1*01:01 | 9.9 | 0.43 |
| | 38 | 52 | PPDTKLENFFSFYRL | HLA-DPA1*02:01/ HLA-DPB1*01:01 | 24.3 | 2.12 |
| | 37 | 51 | LPPDTKLENFFSFYR | HLA-DPA1*02:01/ HLA-DPB1*01:01 | 61.5 | 6.57 |
| | 39 | 53 | PDTKLENFFSFYRLL | HLA-DPA1*03:01/ HLA-DPB1*04:02 | 77.6 | 8.12 |
| | 38 | 52 | PPDTKLENFFSFYRL | HLA-DPA1*03:01/ HLA-DPB1*04:02 | 234.4 | 16.13 |
| | 39 | 53 | PDTKLENFFSFYRLL | HLA-DRB1*04:05 | 326.2 | 22.91 |
| | 40 | 54 | DTKLENFFSFYRLLP | HLA-DRB1*04:05 | 338.4 | 23.4 |
| | 38 | 52 | PPDTKLENFFSFYRL | HLA-DRB1*04:05 | 360 | 24.3 |
| | 37 | 51 | LPPDTKLENFFSFYR | HLA-DRB1*04:05 | 426.6 | 26.81 |
| | 40 | 54 | DTKLENFFSFYRLLP | HLA-DRB1*07:01 | 420.3 | 30.77 |
| | 39 | 53 | PDTKLENFFSFYRLL | HLA-DRB1*07:01 | 457 | 32.01 |
| | 39 | 53 | PDTKLENFFSFYRLL | HLA-DRB1*15:01 | 22.7 | 1.85 |
| | 40 | 54 | DTKLENFFSFYRLLP | HLA-DRB1*15:01 | 22.8 | 1.86 |
| | 41 | 55 | TKLENFFSFYRLLPM | HLA-DRB1*15:01 | 24.1 | 2.02 |
| | 42 | 56 | KLENFFSFYRLLPMG | HLA-DRB1*15:01 | 28.9 | 2.61 |
| | 38 | 52 | PPDTKLENFFSFYRL | HLA-DRB1*15:01 | 32.7 | 3.06 |

| | | | | | | |
|---|---|---|---|---|---|---|
| | 37 | 51 | LPPDTKLENFFSFYR | HLA-DRB1*15:01 | 50.8 | 5.18 |
| | 43 | 57 | LENFFSFYRLLPMGG | HLA-DRB1*15:01 | 68.6 | 7.04 |
| | 42 | 56 | KLENFFSFYRLLPMG | HLA-DRB5*01:01 | 41.7 | 8.82 |
| | 38 | 52 | PPDTKLENFFSFYRL | HLA-DRB5*01:01 | 44 | 9.15 |
| | 43 | 57 | LENFFSFYRLLPMGG | HLA-DRB5*01:01 | 48 | 9.72 |
| | 37 | 51 | LPPDTKLENFFSFYR | HLA-DRB5*01:01 | 93.7 | 14.74 |
| **LSAIPTSYF** | 165 | 179 | SPLSAIPTSYFGWED | HLA-DRB1*01:01 | 168.9 | 38.07 |
| | 167 | 181 | LSAIPTSYFGWEDWS | HLA-DRB1*04:05 | 289.3 | 21.24 |
| | 161 | 175 | IPYTSPLSAIPTSYF | HLA-DRB1*07:01 | 76.6 | 11.85 |
| | 162 | 176 | PYTSPLSAIPTSYFG | HLA-DRB1*07:01 | 132.7 | 16.93 |
| | 163 | 177 | YTSPLSAIPTSYFGW | HLA-DRB1*07:01 | 182.8 | 20.41 |
| | 164 | 178 | TSPLSAIPTSYFGWE | HLA-DRB1*07:01 | 323.1 | 27.2 |
| | 164 | 178 | TSPLSAIPTSYFGWE | HLA-DRB1*08:02 | 138.9 | 2.69 |
| | 163 | 177 | YTSPLSAIPTSYFGW | HLA-DRB1*08:02 | 158 | 3.2 |
| | 165 | 179 | SPLSAIPTSYFGWED | HLA-DRB1*08:02 | 161.1 | 3.27 |
| | 161 | 175 | IPYTSPLSAIPTSYF | HLA-DRB1*08:02 | 214.1 | 4.77 |
| | 162 | 176 | PYTSPLSAIPTSYFG | HLA-DRB1*08:02 | 227.4 | 5.13 |
| | 166 | 180 | PLSAIPTSYFGWEDW | HLA-DRB1*08:02 | 291.7 | 6.85 |
| | 163 | 177 | YTSPLSAIPTSYFGW | HLA-DRB1*09:01 | 130.7 | 8.91 |
| | 164 | 178 | TSPLSAIPTSYFGWE | HLA-DRB1*09:01 | 163.3 | 10.88 |
| | 165 | 179 | SPLSAIPTSYFGWED | HLA-DRB1*09:01 | 247.4 | 15.28 |
| | 166 | 180 | PLSAIPTSYFGWEDW | HLA-DRB1*09:01 | 472.6 | 24.58 |
| **LVAFAPPGA** | 120 | 134 | LVAFAPPGATIPLKP | HLA-DRB1*01:01 | 60.4 | 23.57 |
| | 117 | 131 | ATMLVAFAPPGATIP | HLA-DRB1*09:01 | 11 | 0.22 |
| | 116 | 130 | PATMLVAFAPPGATI | HLA-DRB1*09:01 | 11.6 | 0.25 |
| | 118 | 132 | TMLVAFAPPGATIPL | HLA-DRB1*09:01 | 14.1 | 0.4 |
| | 115 | 129 | NPATMLVAFAPPGAT | HLA-DRB1*09:01 | 14.8 | 0.44 |
| | 114 | 128 | DNPATMLVAFAPPGA | HLA-DRB1*09:01 | 21.1 | 0.87 |
| | 119 | 133 | MLVAFAPPGATIPLK | HLA-DRB1*09:01 | 26.8 | 1.31 |
| | 120 | 134 | LVAFAPPGATIPLKP | HLA-DRB1*09:01 | 40 | 2.41 |
| | 117 | 131 | ATMLVAFAPPGATIP | HLA-DRB1*15:01 | 18.7 | 1.36 |
| | 116 | 130 | PATMLVAFAPPGATI | HLA-DRB1*15:01 | 21.8 | 1.75 |
| | 118 | 132 | TMLVAFAPPGATIPL | HLA-DRB1*15:01 | 22.4 | 1.82 |
| | 119 | 133 | MLVAFAPPGATIPLK | HLA-DRB1*15:01 | 28.6 | 2.58 |
| | 115 | 129 | NPATMLVAFAPPGAT | HLA-DRB1*15:01 | 36.9 | 3.57 |
| | 114 | 128 | DNPATMLVAFAPPGA | HLA-DRB1*15:01 | 60.7 | 6.24 |
| | 120 | 134 | LVAFAPPGATIPLKP | HLA-DRB1*15:01 | 92.5 | 9.28 |
| **MAEVPVSAA** | 141 | 155 | NFYMAEVPVSAATST | HLA-DRB1*08:02 | 91.5 | 1.37 |
| | 140 | 154 | SNFYMAEVPVSAATS | HLA-DRB1*08:02 | 93.5 | 1.42 |
| | 142 | 156 | FYMAEVPVSAATSTM | HLA-DRB1*08:02 | 146.1 | 2.89 |
| | 143 | 157 | YMAEVPVSAATSTMV | HLA-DRB1*08:02 | 158.7 | 3.22 |
| | 144 | 158 | MAEVPVSAATSTMVS | HLA-DRB1*08:02 | 304.5 | 7.21 |
| | 140 | 154 | SNFYMAEVPVSAATS | HLA-DQA1*01:02/ DQB1*06:02 | 33.3 | 1.53 |

| | | | | | | |
|---|---|---|---|---|---|---|
| | 139 | 153 | LSNFYMAEVPVSAAT | HLA-DQA1*01:02/ DQB1*06:02 | 34.4 | 1.63 |
| | 141 | 155 | NFYMAEVPVSAATST | HLA-DQA1*01:02/ DQB1*06:02 | 38.6 | 1.97 |
| | 138 | 152 | MLSNFYMAEVPVSAA | HLA-DQA1*01:02/ DQB1*06:02 | 38.9 | 1.99 |
| | 142 | 156 | FYMAEVPVSAATSTM | HLA-DQA1*01:02/ DQB1*06:02 | 53.1 | 3.21 |
| | 143 | 157 | YMAEVPVSAATSTMV | HLA-DQA1*01:02/ DQB1*06:02 | 65.7 | 4.29 |
| **MLSCFTYIA** | 89 | 103 | IAAMLSCFTYIAADL | HLA-DRB1*01:01 | 76.2 | 26.52 |
| **MLVAFAPPG** | 114 | 128 | DNPATMLVAFAPPGA | HLA-DQA1*05:01/ DQB1*03:01 | 195.4 | 22.28 |
| | 113 | 127 | NDNPATMLVAFAPPG | HLA-DQA1*05:01/ DQB1*03:01 | 309.3 | 28.47 |
| | 116 | 130 | PATMLVAFAPPGATI | HLA-DRB1*01:01 | 6.8 | 2.46 |
| | 117 | 131 | ATMLVAFAPPGATIP | HLA-DRB1*01:01 | 7.6 | 3.19 |
| | 115 | 129 | NPATMLVAFAPPGAT | HLA-DRB1*01:01 | 10.5 | 5.57 |
| | 118 | 132 | TMLVAFAPPGATIPL | HLA-DRB1*01:01 | 10.7 | 5.73 |
| | 114 | 128 | DNPATMLVAFAPPGA | HLA-DRB1*01:01 | 16.8 | 9.63 |
| | 119 | 133 | MLVAFAPPGATIPLK | HLA-DRB1*01:01 | 17.5 | 10.01 |
| | 113 | 127 | NDNPATMLVAFAPPG | HLA-DRB1*01:01 | 40.1 | 18.67 |
| | 116 | 130 | PATMLVAFAPPGATI | HLA-DRB1*04:01 | 80.6 | 6.53 |
| | 117 | 131 | ATMLVAFAPPGATIP | HLA-DRB1*04:01 | 93 | 7.54 |
| | 115 | 129 | NPATMLVAFAPPGAT | HLA-DRB1*04:01 | 125.7 | 10.07 |
| | 118 | 132 | TMLVAFAPPGATIPL | HLA-DRB1*04:01 | 155.7 | 12.18 |
| | 114 | 128 | DNPATMLVAFAPPGA | HLA-DRB1*04:01 | 166.7 | 12.91 |
| | 119 | 133 | MLVAFAPPGATIPLK | HLA-DRB1*04:01 | 262.4 | 18.48 |
| | 113 | 127 | NDNPATMLVAFAPPG | HLA-DRB1*04:01 | 427.2 | 26.03 |
| | 119 | 133 | MLVAFAPPGATIPLK | HLA-DRB1*04:04 | 120.5 | 13.92 |
| | 117 | 131 | ATMLVAFAPPGATIP | HLA-DRB1*08:02 | 90.8 | 1.34 |
| | 116 | 130 | PATMLVAFAPPGATI | HLA-DRB1*08:02 | 97.2 | 1.53 |
| | 118 | 132 | TMLVAFAPPGATIPL | HLA-DRB1*08:02 | 117.5 | 2.09 |
| | 115 | 129 | NPATMLVAFAPPGAT | HLA-DRB1*08:02 | 150.7 | 3.02 |
| | 119 | 133 | MLVAFAPPGATIPLK | HLA-DRB1*08:02 | 189.5 | 4.08 |
| | 114 | 128 | DNPATMLVAFAPPGA | HLA-DRB1*08:02 | 261.1 | 6.02 |
| | 117 | 131 | ATMLVAFAPPGATIP | HLA-DRB1*11:01 | 266.9 | 23.42 |
| | 116 | 130 | PATMLVAFAPPGATI | HLA-DRB1*11:01 | 270.4 | 23.56 |
| | 118 | 132 | TMLVAFAPPGATIPL | HLA-DRB1*11:01 | 380.8 | 27.61 |
| | 113 | 127 | NDNPATMLVAFAPPG | HLA-DRB1*15:01 | 388.6 | 25.45 |
| **MLSCFTYIA** | 90 | 104 | AAMLSCFTYIAADLR | HLA-DPA1*01/ HLA-DPB1*04:01 | 97.1 | 5.18 |
| | 89 | 103 | IAAMLSCFTYIAADL | HLA-DPA1*01/ HLA-DPB1*04:01 | 104.7 | 5.48 |
| | 91 | 105 | AMLSCFTYIAADLRI | HLA-DPA1*01/ HLA-DPB1*04:01 | 126.5 | 6.25 |
| | 88 | 102 | GIAAMLSCFTYIAAD | HLA-DPA1*01/ HLA-DPB1*04:01 | 133 | 6.47 |
| | 87 | 101 | SGIAAMLSCFTYIAA | HLA-DPA1*01/ HLA-DPB1*04:01 | 146.3 | 6.93 |
| | 92 | 106 | MLSCFTYIAADLRIT | HLA-DPA1*01/ HLA-DPB1*04:01 | 193.5 | 8.35 |
| **MVSFSIPYT** | 153 | 167 | TSTMVSFSIPYTSPL | HLA-DQA1*01:01/ DQB1*05:01 | 184.9 | 4.04 |
| | 154 | 168 | STMVSFSIPYTSPLS | HLA-DQA1*01:01/ DQB1*05:01 | 196.3 | 4.26 |
| | 152 | 166 | ATSTMVSFSIPYTSP | HLA-DQA1*01:01/ DQB1*05:01 | 235.5 | 4.98 |
| | 155 | 169 | TMVSFSIPYTSPLSA | HLA-DQA1*01:01/ DQB1*05:01 | 238.5 | 5.04 |

| | | | | | | |
|---|---|---|---|---|---|---|
| | 151 | 165 | AATSTMVSFSIPYTS | HLA-DQA1*01:01/ DQB1*05:01 | 347.6 | 6.85 |
| | 153 | 167 | TSTMVSFSIPYTSPL | HLA-DRB1*01:01 | 173.8 | 38.54 |
| | 152 | 166 | ATSTMVSFSIPYTSP | HLA-DRB1*01:01 | 316.4 | 49.06 |
| | 153 | 167 | TSTMVSFSIPYTSPL | HLA-DRB1*04:05 | 467.8 | 28.24 |
| | 155 | 169 | TMVSFSIPYTSPLSA | HLA-DRB1*08:02 | 178 | 3.77 |
| | 154 | 168 | STMVSFSIPYTSPLS | HLA-DRB1*08:02 | 257.5 | 5.93 |
| | 153 | 167 | TSTMVSFSIPYTSPL | HLA-DRB1*08:02 | 300.1 | 7.09 |
| | 152 | 166 | ATSTMVSFSIPYTSP | HLA-DRB1*08:02 | 415.4 | 10.07 |
| | 153 | 167 | TSTMVSFSIPYTSPL | HLA-DRB1*15:01 | 43.5 | 4.34 |
| | 154 | 168 | STMVSFSIPYTSPLS | HLA-DRB1*15:01 | 47 | 4.74 |
| | 152 | 166 | ATSTMVSFSIPYTSP | HLA-DRB1*15:01 | 52 | 5.3 |
| | 155 | 169 | TMVSFSIPYTSPLSA | HLA-DRB1*15:01 | 54.1 | 5.53 |
| | 151 | 165 | AATSTMVSFSIPYTS | HLA-DRB1*15:01 | 62.4 | 6.42 |
| | 150 | 164 | SAATSTMVSFSIPYT | HLA-DRB1*15:01 | 83.4 | 8.46 |
| | 156 | 170 | MVSFSIPYTSPLSAI | HLA-DRB1*15:01 | 106.7 | 10.49 |
| **NFGQLSSGS** | 182 | 196 | GTNFGQLSSGSWGNL | HLA-DQA1*05:01/ DQB1*03:01 | 249.2 | 25.42 |
| **NFFSFYRLL** | 43 | 57 | LENFFSFYRLLPMGG | HLA-DPA1*02:01/ HLA-DPB1*05:01 | 128.1 | 2.7 |
| **PLSAIPTSY** | 161 | 175 | IPYTSPLSAIPTSYF | HLA-DQA1*05:01/ DQB1*03:01 | 75.9 | 12.3 |
| | 162 | 176 | PYTSPLSAIPTSYFG | HLA-DQA1*05:01/ DQB1*03:01 | 76.4 | 12.35 |
| | 163 | 177 | YTSPLSAIPTSYFGW | HLA-DQA1*05:01/ DQB1*03:01 | 77.5 | 12.48 |
| | 160 | 174 | SIPYTSPLSAIPTSY | HLA-DQA1*05:01/ DQB1*03:01 | 79.5 | 12.71 |
| | 164 | 178 | TSPLSAIPTSYFGWE | HLA-DQA1*05:01/ DQB1*03:01 | 128.4 | 17.41 |
| | 165 | 179 | SPLSAIPTSYFGWED | HLA-DQA1*05:01/ DQB1*03:01 | 265.7 | 26.29 |
| | 166 | 180 | PLSAIPTSYFGWEDW | HLA-DQA1*05:01/ DQB1*03:01 | 334 | 29.59 |
| **PNDNPATML** | 109 | 123 | FSNPNDNPATMLVAF | HLA-DRB1*01:01 | 499.6 | 56.72 |
| **PYTSPLSAI** | 159 | 173 | FSIPYTSPLSAIPTS | HLA-DQA1*05:01/ DQB1*03:01 | 160.5 | 19.9 |
| | 158 | 172 | SFSIPYTSPLSAIPT | HLA-DQA1*05:01/ DQB1*03:01 | 168.2 | 20.46 |
| **QLSCWVAFG** | 209 | 223 | PFDFQLSCWVAFGNF | HLA-DQA1*01:02/ DQB1*06:02 | 473.8 | 26.41 |
| **RVYIVRAQR** | 264 | 278 | PDDRVYIVRAQRPTY | HLA-DRB1*08:02 | 101.5 | 1.65 |
| | 263 | 277 | DPDDRVYIVRAQRPT | HLA-DRB1*08:02 | 289.7 | 6.8 |
| | 262 | 276 | VDPDDRVYIVRAQRP | HLA-DRB1*08:02 | 456.1 | 11.08 |
| | 264 | 278 | PDDRVYIVRAQRPTY | HLA-DRB1*11:01 | 15.3 | 2.55 |
| | 263 | 277 | DPDDRVYIVRAQRPT | HLA-DRB1*11:01 | 29.6 | 5.44 |
| | 262 | 276 | VDPDDRVYIVRAQRP | HLA-DRB1*11:01 | 60.3 | 9.8 |
| | 264 | 278 | PDDRVYIVRAQRPTY | HLA-DRB1*15:01 | 140.8 | 13.15 |
| | 263 | 277 | DPDDRVYIVRAQRPT | HLA-DRB1*15:01 | 313.8 | 22.52 |
| | 262 | 276 | VDPDDRVYIVRAQRP | HLA-DRB1*15:01 | 493.5 | 28.86 |
| | 264 | 278 | PDDRVYIVRAQRPTY | HLA-DRB5*01:01 | 129.9 | 17.64 |
| | 263 | 277 | DPDDRVYIVRAQRPT | HLA-DRB5*01:01 | 245.2 | 24.27 |
| **SAATSTMVS** | 147 | 161 | VPVSAATSTMVSFSI | HLA-DQA1*01:02/ DQB1*06:02 | 23.8 | 0.82 |
| | 148 | 162 | PVSAATSTMVSFSIP | HLA-DQA1*01:02/ DQB1*06:02 | 32.4 | 1.46 |
| | 146 | 160 | EVPVSAATSTMVSFS | HLA-DQA1*01:02/ DQB1*06:02 | 37.4 | 1.87 |

| | | | | | | |
|---|---|---|---|---|---|---|
| | 149 | 163 | VSAATSTMVSFSIPY | HLA-DQA1*01:02/ DQB1*06:02 | 42.7 | 2.32 |
| | 145 | 159 | AEVPVSAATSTMVSF | HLA-DQA1*01:02/ DQB1*06:02 | 47.2 | 2.71 |
| | 144 | 158 | MAEVPVSAATSTMVS | HLA-DQA1*01:02/ DQB1*06:02 | 57.3 | 3.57 |
| | 150 | 164 | SAATSTMVSFSIPYT | HLA-DQA1*01:02/ DQB1*06:02 | 65.5 | 4.27 |
| **SCFTYIAAD** | 89 | 103 | IAAMLSCFTYIAADL | HLA-DQA1*01:02/ DQB1*06:02 | 46 | 2.6 |
| | 90 | 104 | AAMLSCFTYIAADLR | HLA-DQA1*01:02/ DQB1*06:02 | 49.6 | 2.91 |
| | 91 | 105 | AMLSCFTYIAADLRI | HLA-DQA1*01:02/ DQB1*06:02 | 54.3 | 3.32 |
| | 88 | 102 | GIAAMLSCFTYIAAD | HLA-DQA1*01:02/ DQB1*06:02 | 67.7 | 4.46 |
| | 92 | 106 | MLSCFTYIAADLRIT | HLA-DQA1*01:02/ DQB1*06:02 | 70.4 | 4.69 |
| | 93 | 107 | LSCFTYIAADLRITL | HLA-DQA1*01:02/ DQB1*06:02 | 93.2 | 6.57 |
| | 94 | 108 | SCFTYIAADLRITLR | HLA-DQA1*01:02/ DQB1*06:02 | 349.9 | 21.57 |
| | 91 | 105 | AMLSCFTYIAADLRI | HLA-DQA1*04:01/ DQB1*04:02 | 374.1 | 5.6 |
| | 90 | 104 | AAMLSCFTYIAADLR | HLA-DQA1*04:01/ DQB1*04:02 | 465.6 | 7.21 |
| | 92 | 106 | MLSCFTYIAADLRIT | HLA-DQA1*04:01/ DQB1*04:02 | 467.4 | 7.24 |
| | 94 | 108 | SCFTYIAADLRITLR | HLA-DRB1*04:04 | 88.1 | 10.58 |
| | 90 | 104 | AAMLSCFTYIAADLR | HLA-DRB1*04:05 | 76.9 | 7.56 |
| | 89 | 103 | IAAMLSCFTYIAADL | HLA-DRB1*04:05 | 206.6 | 16.95 |
| | 88 | 102 | GIAAMLSCFTYIAAD | HLA-DRB1*04:05 | 294.3 | 21.48 |
| **SIPYTSPLS** | 157 | 171 | VSFSIPYTSPLSAIP | HLA-DRB1*04:01 | 190.3 | 14.4 |
| | 157 | 171 | VSFSIPYTSPLSAIP | HLA-DRB1*15:01 | 193.4 | 16.58 |
| | 158 | 172 | SFSIPYTSPLSAIPT | HLA-DRB1*15:01 | 262.9 | 20.22 |
| | 159 | 173 | FSIPYTSPLSAIPTS | HLA-DRB1*15:01 | 379.4 | 25.12 |
| **SPLSAIPTS** | 162 | 176 | PYTSPLSAIPTSYFG | HLA-DRB1*04:04 | 102.4 | 12.12 |
| | 163 | 177 | YTSPLSAIPTSYFGW | HLA-DRB1*04:04 | 105.3 | 12.44 |
| | 161 | 175 | IPYTSPLSAIPTSYF | HLA-DRB1*04:04 | 106.3 | 12.54 |
| | 164 | 178 | TSPLSAIPTSYFGWE | HLA-DRB1*04:04 | 111 | 13 |
| | 165 | 179 | SPLSAIPTSYFGWED | HLA-DRB1*04:04 | 173.4 | 18.34 |
| | 160 | 174 | SIPYTSPLSAIPTSY | HLA-DRB1*04:04 | 175.9 | 18.53 |
| **SYFGWEDWS** | 170 | 184 | IPTSYFGWEDWSGTN | HLA-DQA1*01:01/ DQB1*05:01 | 50.7 | 1 |
| | 171 | 185 | PTSYFGWEDWSGTNF | HLA-DQA1*01:01/ DQB1*05:01 | 52.5 | 1.05 |
| | 172 | 186 | TSYFGWEDWSGTNFG | HLA-DQA1*01:01/ DQB1*05:01 | 56.6 | 1.15 |
| | 169 | 183 | AIPTSYFGWEDWSGT | HLA-DQA1*01:01/ DQB1*05:01 | 58.8 | 1.21 |
| | 168 | 182 | SAIPTSYFGWEDWSG | HLA-DQA1*01:01/ DQB1*05:01 | 82.5 | 1.79 |
| | 173 | 187 | SYFGWEDWSGTNFGQ | HLA-DQA1*01:01/ DQB1*05:01 | 110.7 | 2.44 |
| | 167 | 181 | LSAIPTSYFGWEDWS | HLA-DQA1*01:01/ DQB1*05:01 | 156.1 | 3.45 |
| **TKLENFFSF** | 37 | 51 | LPPDTKLENFFSFYR | HLA-DPA1*01:03/ HLA-DPB1*02:01 | 125.1 | 9.64 |
| | 36 | 50 | PLPPDTKLENFFSFY | HLA-DPA1*01:03/ HLA-DPB1*02:01 | 276 | 15.53 |
| **TSYFGWEDW** | 168 | 182 | SAIPTSYFGWEDWSG | HLA-DPA1*01:03/ HLA-DPB1*02:01 | 81.9 | 7.32 |
| | 167 | 181 | LSAIPTSYFGWEDWS | HLA-DPA1*01:03/ HLA-DPB1*02:01 | 132.6 | 10.01 |
| | 166 | 180 | PLSAIPTSYFGWEDW | HLA-DPA1*01:03/ HLA- | 234.3 | 14.13 |

| | | | | | | |
|---|---|---|---|---|---|---|
| | | | | DPB1*02:01 | | |
| **TMLVAFAPP** | 116 | 130 | PATMLVAFAPPGATI | HLA-DRB1*04:04 | 23.5 | 1.89 |
| | 115 | 129 | NPATMLVAFAPPGAT | HLA-DRB1*04:04 | 25.5 | 2.19 |
| | 117 | 131 | ATMLVAFAPPGATIP | HLA-DRB1*04:04 | 26.4 | 2.31 |
| | 118 | 132 | TMLVAFAPPGATIPL | HLA-DRB1*04:04 | 30.6 | 2.89 |
| | 114 | 128 | DNPATMLVAFAPPGA | HLA-DRB1*04:04 | 32 | 3.11 |
| | 113 | 127 | NDNPATMLVAFAPPG | HLA-DRB1*04:04 | 67.2 | 8.05 |
| | 112 | 126 | PNDNPATMLVAFAPP | HLA-DRB1*04:04 | 127.1 | 14.55 |
| **TMVSFSIPY** | 152 | 166 | ATSTMVSFSIPYTSP | HLA-DRB1*04:01 | 367.3 | 23.52 |
| | 155 | 169 | TMVSFSIPYTSPLSA | HLA-DRB1*04:04 | 18.8 | 1.21 |
| | 153 | 167 | TSTMVSFSIPYTSPL | HLA-DRB1*04:04 | 28 | 2.55 |
| | 154 | 168 | STMVSFSIPYTSPLS | HLA-DRB1*04:04 | 29.1 | 2.7 |
| | 152 | 166 | ATSTMVSFSIPYTSP | HLA-DRB1*04:04 | 30.3 | 2.86 |
| | 151 | 165 | AATSTMVSFSIPYTS | HLA-DRB1*04:04 | 32.8 | 3.22 |
| | 150 | 164 | SAATSTMVSFSIPYT | HLA-DRB1*04:04 | 38.4 | 4.06 |
| | 149 | 163 | VSAATSTMVSFSIPY | HLA-DRB1*04:04 | 130.2 | 14.82 |
| | 150 | 164 | SAATSTMVSFSIPYT | HLA-DRB1*07:01 | 427.2 | 31.02 |
| | 151 | 165 | AATSTMVSFSIPYTS | HLA-DRB1*07:01 | 460.9 | 32.12 |
| | 149 | 163 | VSAATSTMVSFSIPY | HLA-DRB1*15:01 | 258.8 | 20.04 |
| **TSPLSAIPT** | 161 | 175 | IPYTSPLSAIPTSYF | HLA-DRB1*01:01 | 10.8 | 5.8 |
| | 160 | 174 | SIPYTSPLSAIPTSY | HLA-DRB1*01:01 | 15.5 | 8.9 |
| | 162 | 176 | PYTSPLSAIPTSYFG | HLA-DRB1*01:01 | 18.1 | 10.33 |
| | 163 | 177 | YTSPLSAIPTSYFGW | HLA-DRB1*01:01 | 21.4 | 11.97 |
| | 159 | 173 | FSIPYTSPLSAIPTS | HLA-DRB1*01:01 | 34 | 16.82 |
| | 158 | 172 | SFSIPYTSPLSAIPT | HLA-DRB1*01:01 | 62.7 | 24.03 |
| **TYIAADLRI** | 95 | 109 | CFTYIAADLRITLRF | HLA-DRB1*07:01 | 59.9 | 9.95 |
| | 96 | 110 | FTYIAADLRITLRFS | HLA-DRB1*07:01 | 78.7 | 12.09 |
| | 97 | 111 | TYIAADLRITLRFSN | HLA-DRB1*07:01 | 197.4 | 21.26 |
| | 93 | 107 | LSCFTYIAADLRITL | HLA-DRB1*15:01 | 274.6 | 20.78 |
| | 94 | 108 | SCFTYIAADLRITLR | HLA-DRB1*15:01 | 291.5 | 21.55 |
| | 94 | 108 | SCFTYIAADLRITLR | HLA-DPA1*02:01/ HLA-DPB1*05:01 | 395 | 8.46 |
| **VAFAPPGAT** | 121 | 135 | VAFAPPGATIPLKPT | HLA-DRB1*01:01 | 416.2 | 53.75 |
| | 118 | 132 | TMLVAFAPPGATIPL | HLA-DRB5*01:01 | 445.6 | 31.51 |
| | 118 | 132 | TMLVAFAPPGATIPL | HLA-DQA1*05:01/ DQB1*03:01 | 6.5 | 0.64 |
| | 119 | 133 | MLVAFAPPGATIPLK | HLA-DQA1*05:01/ DQB1*03:01 | 6.5 | 0.64 |
| | 117 | 131 | ATMLVAFAPPGATIP | HLA-DQA1*05:01/ DQB1*03:01 | 6.9 | 0.73 |
| | 120 | 134 | LVAFAPPGATIPLKP | HLA-DQA1*05:01/ DQB1*03:01 | 7 | 0.75 |
| | 121 | 135 | VAFAPPGATIPLKPT | HLA-DQA1*05:01/ DQB1*03:01 | 7.2 | 0.79 |
| | 116 | 130 | PATMLVAFAPPGATI | HLA-DQA1*05:01/ DQB1*03:01 | 8.6 | 1.12 |
| | 115 | 129 | NPATMLVAFAPPGAT | HLA-DQA1*05:01/ DQB1*03:01 | 10.6 | 1.58 |
| **VAFGNFKAW** | 213 | 227 | QLSCWVAFGNFKAWV | HLA-DPA1*01:03/ HLA-DPB1*02:01 | 81.2 | 7.27 |

| | | | | | | |
|---|---|---|---|---|---|---|
| | 214 | 228 | LSCWVAFGNFKAWVP | HLA-DPA1*01:03/ HLA-DPB1*02:01 | 86.8 | 7.61 |
| | 215 | 229 | SCWVAFGNFKAWVPR | HLA-DPA1*01:03/ HLA-DPB1*02:01 | 97.6 | 8.23 |
| | 216 | 230 | CWVAFGNFKAWVPRP | HLA-DPA1*01:03/ HLA-DPB1*02:01 | 153.9 | 10.99 |
| | 217 | 231 | WVAFGNFKAWVPRPP | HLA-DPA1*01:03/ HLA-DPB1*02:01 | 279.5 | 15.64 |
| | 218 | 232 | VAFGNFKAWVPRPPP | HLA-DPA1*01:03/ HLA-DPB1*02:01 | 388.8 | 18.79 |
| | 214 | 228 | LSCWVAFGNFKAWVP | HLA-DQA1*05:01/ DQB1*03:01 | 231.8 | 24.46 |
| | 213 | 227 | QLSCWVAFGNFKAWV | HLA-DQA1*05:01/ DQB1*03:01 | 340.6 | 29.88 |
| | 212 | 226 | FQLSCWVAFGNFKAW | HLA-DQA1*05:01/ DQB1*03:01 | 342.3 | 29.96 |
| **VPVSAATST** | 147 | 161 | VPVSAATSTMVSFSI | HLA-DRB1*04:04 | 187.4 | 19.43 |
| | 146 | 160 | EVPVSAATSTMVSFS | HLA-DRB1*04:04 | 213.8 | 21.26 |
| | 145 | 159 | AEVPVSAATSTMVSF | HLA-DRB1*04:04 | 234.8 | 22.69 |
| | 144 | 158 | MAEVPVSAATSTMVS | HLA-DRB1*04:04 | 284.7 | 25.47 |
| | 141 | 155 | NFYMAEVPVSAATST | HLA-DRB1*04:04 | 290.9 | 25.79 |
| | 153 | 167 | TSTMVSFSIPYTSPL | HLA-DQA1*01:02/ DQB1*06:02 | 184.3 | 13.04 |
| | 152 | 166 | ATSTMVSFSIPYTSP | HLA-DQA1*01:02/ DQB1*06:02 | 191.6 | 13.49 |
| | 151 | 165 | AATSTMVSFSIPYTS | HLA-DQA1*01:02/ DQB1*06:02 | 207.8 | 14.45 |
| | 154 | 168 | STMVSFSIPYTSPLS | HLA-DQA1*01:02/ DQB1*06:02 | 286.6 | 18.67 |
| | 154 | 168 | STMVSFSIPYTSPLS | HLA-DQA1*05:01/ DQB1*03:01 | 78.5 | 12.6 |
| | 157 | 171 | VSFSIPYTSPLSAIP | HLA-DQA1*05:01/ DQB1*03:01 | 85.3 | 13.35 |
| | 153 | 167 | TSTMVSFSIPYTSPL | HLA-DQA1*05:01/ DQB1*03:01 | 86.9 | 13.52 |
| | 155 | 169 | TMVSFSIPYTSPLSA | HLA-DQA1*05:01/ DQB1*03:01 | 94.7 | 14.32 |
| | 156 | 170 | MVSFSIPYTSPLSAI | HLA-DQA1*05:01/ DQB1*03:01 | 95.9 | 14.43 |
| | 152 | 166 | ATSTMVSFSIPYTSP | HLA-DQA1*05:01/ DQB1*03:01 | 97.7 | 14.6 |
| | 151 | 165 | AATSTMVSFSIPYTS | HLA-DQA1*05:01/ DQB1*03:01 | 117 | 16.44 |
| | 155 | 169 | TMVSFSIPYTSPLSA | HLA-DRB1*04:01 | 155.5 | 12.17 |
| | 154 | 168 | STMVSFSIPYTSPLS | HLA-DRB1*04:01 | 156.6 | 12.24 |
| | 153 | 167 | TSTMVSFSIPYTSPL | HLA-DRB1*04:01 | 203.7 | 15.22 |
| **VYIVRAQRP** | 264 | 278 | PDDRVYIVRAQRPTY | HLA-DQA1*05:01/ DQB1*03:01 | 308.1 | 28.41 |
| | 263 | 277 | DPDDRVYIVRAQRPT | HLA-DQA1*05:01/ DQB1*03:01 | 488.4 | 35.4 |
| | 264 | 278 | PDDRVYIVRAQRPTY | HLA-DRB1*01:01 | 21.1 | 11.83 |
| | 263 | 277 | DPDDRVYIVRAQRPT | HLA-DRB1*01:01 | 49.7 | 21.17 |
| | 262 | 276 | VDPDDRVYIVRAQRP | HLA-DRB1*01:01 | 106.2 | 31.04 |
| | 264 | 278 | PDDRVYIVRAQRPTY | HLA-DRB1*04:05 | 494.2 | 29.11 |
| | 264 | 278 | PDDRVYIVRAQRPTY | HLA-DRB1*07:01 | 183.5 | 20.46 |
| | 263 | 277 | DPDDRVYIVRAQRPT | HLA-DRB1*07:01 | 204.8 | 21.65 |
| | 262 | 276 | VDPDDRVYIVRAQRP | HLA-DRB1*07:01 | 343.7 | 28.05 |
| | 264 | 278 | PDDRVYIVRAQRPTY | HLA-DRB1*13:02 | 334.1 | 13.63 |
| | 262 | 276 | VDPDDRVYIVRAQRP | HLA-DRB1*13:02 | 334.6 | 13.64 |
| | 263 | 277 | DPDDRVYIVRAQRPT | HLA-DRB1*13:02 | 443.2 | 16.19 |

| | | | | | | |
|---|---|---|---|---|---|---|
| | 264 | 278 | PDDRVYIVRAQRPTY | HLA-DRB4*01:01 | 46 | 3.35 |
| | 263 | 277 | DPDDRVYIVRAQRPT | HLA-DRB4*01:01 | 70 | 5.43 |
| | 262 | 276 | VDPDDRVYIVRAQRP | HLA-DRB4*01:01 | 154 | 11.51 |
| **WVAFGNFKA** | 213 | 227 | QLSCWVAFGNFKAWV | HLA-DRB1*01:01 | 14.9 | 8.54 |
| | 214 | 228 | LSCWVAFGNFKAWVP | HLA-DRB1*01:01 | 17.1 | 9.79 |
| | 212 | 226 | FQLSCWVAFGNFKAW | HLA-DRB1*01:01 | 22.2 | 12.33 |
| | 215 | 229 | SCWVAFGNFKAWVPR | HLA-DRB1*01:01 | 34.9 | 17.11 |
| | 211 | 225 | DFQLSCWVAFGNFKA | HLA-DRB1*01:01 | 40.6 | 18.81 |
| | 216 | 230 | CWVAFGNFKAWVPRP | HLA-DRB1*01:01 | 66.6 | 24.79 |
| | 217 | 231 | WVAFGNFKAWVPRPP | HLA-DRB1*01:01 | 96.9 | 29.75 |
| | 214 | 228 | LSCWVAFGNFKAWVP | HLA-DRB1*04:01 | 283 | 19.56 |
| | 211 | 225 | DFQLSCWVAFGNFKA | HLA-DRB1*04:01 | 355.6 | 23.02 |
| | 212 | 226 | FQLSCWVAFGNFKAW | HLA-DRB1*04:01 | 431 | 26.18 |
| | 213 | 227 | QLSCWVAFGNFKAWV | HLA-DRB1*04:01 | 442.3 | 26.62 |
| | 215 | 229 | SCWVAFGNFKAWVPR | HLA-DRB1*04:01 | 448.4 | 26.86 |
| | 217 | 231 | WVAFGNFKAWVPRPP | HLA-DRB1*04:04 | 82.7 | 9.96 |
| | 215 | 229 | SCWVAFGNFKAWVPR | HLA-DRB1*04:04 | 99.5 | 11.83 |
| | 216 | 230 | CWVAFGNFKAWVPRP | HLA-DRB1*04:04 | 142.7 | 15.91 |
| | 212 | 226 | FQLSCWVAFGNFKAW | HLA-DRB1*04:05 | 348.4 | 23.83 |
| | 211 | 225 | DFQLSCWVAFGNFKA | HLA-DRB1*04:05 | 355.8 | 24.13 |
| | 211 | 225 | DFQLSCWVAFGNFKA | HLA-DRB1*09:01 | 88 | 6.11 |
| | 212 | 226 | FQLSCWVAFGNFKAW | HLA-DRB1*09:01 | 122.5 | 8.38 |
| | 214 | 228 | LSCWVAFGNFKAWVP | HLA-DRB1*09:01 | 195 | 12.68 |
| | 215 | 229 | SCWVAFGNFKAWVPR | HLA-DRB1*09:01 | 228.2 | 14.37 |
| | 213 | 227 | QLSCWVAFGNFKAWV | HLA-DRB1*09:01 | 258.7 | 15.83 |
| | 216 | 230 | CWVAFGNFKAWVPRP | HLA-DRB1*09:01 | 333.1 | 19.23 |
| | 214 | 228 | LSCWVAFGNFKAWVP | HLA-DRB1*11:01 | 131.3 | 16.13 |
| | 217 | 231 | WVAFGNFKAWVPRPP | HLA-DRB1*11:01 | 164.6 | 18.3 |
| | 215 | 229 | SCWVAFGNFKAWVPR | HLA-DRB1*11:01 | 175.8 | 18.97 |
| | 213 | 227 | QLSCWVAFGNFKAWV | HLA-DRB1*11:01 | 201.8 | 20.39 |
| | 216 | 230 | CWVAFGNFKAWVPRP | HLA-DRB1*11:01 | 222.1 | 21.4 |
| | 212 | 226 | FQLSCWVAFGNFKAW | HLA-DRB1*11:01 | 298.5 | 24.7 |
| | 211 | 225 | DFQLSCWVAFGNFKA | HLA-DRB1*11:01 | 437 | 29.32 |
| **YFGWEDWSG** | 171 | 185 | PTSYFGWEDWSGTNF | HLA-DRB1*04:01 | 377.5 | 23.98 |
| | 170 | 184 | IPTSYFGWEDWSGTN | HLA-DPA1*01/ HLA-DPB1*04:01 | 159.5 | 7.34 |
| | 174 | 188 | YFGWEDWSGTNFGQL | HLA-DPA1*01/ HLA-DPB1*04:01 | 176.8 | 7.87 |
| | 168 | 182 | SAIPTSYFGWEDWSG | HLA-DPA1*01/ HLA-DPB1*04:01 | 432.1 | 13.69 |
| **YIAADLRIT** | 95 | 109 | CFTYIAADLRITLRF | HLA-DPA1*03:01/ HLA-DPB1*04:02 | 41.7 | 4.91 |
| | 96 | 110 | FTYIAADLRITLRFS | HLA-DPA1*03:01/ HLA-DPB1*04:02 | 47.7 | 5.53 |
| | 94 | 108 | SCFTYIAADLRITLR | HLA-DPA1*03:01/ HLA-DPB1*04:02 | 53.5 | 6.1 |

| | | | | | | |
|---|---|---|---|---|---|---|
| | 93 | 107 | LSCFTYIAADLRITL | HLA-DPA1*03:01/ HLA-DPB1*04:02 | 77.3 | 8.1 |
| | 97 | 111 | TYIAADLRITLRFSN | HLA-DPA1*03:01/ HLA-DPB1*04:02 | 80.2 | 8.31 |
| | 98 | 112 | YIAADLRITLRFSNP | HLA-DPA1*03:01/ HLA-DPB1*04:02 | 138.3 | 11.89 |
| | 95 | 109 | CFTYIAADLRITLRF | HLA-DRB1*01:01 | 11.1 | 6.03 |
| | 96 | 110 | FTYIAADLRITLRFS | HLA-DRB1*01:01 | 15.8 | 9.07 |
| | 97 | 111 | TYIAADLRITLRFSN | HLA-DRB1*01:01 | 38.3 | 18.14 |
| | 95 | 109 | CFTYIAADLRITLRF | HLA-DRB1*11:01 | 16 | 2.7 |
| | 96 | 110 | FTYIAADLRITLRFS | HLA-DRB1*11:01 | 20.4 | 3.66 |
| | 94 | 108 | SCFTYIAADLRITLR | HLA-DRB1*11:01 | 24.9 | 4.57 |
| | 93 | 107 | LSCFTYIAADLRITL | HLA-DRB1*11:01 | 44.3 | 7.76 |
| | 92 | 106 | MLSCFTYIAADLRIT | HLA-DRB1*11:01 | 99.6 | 13.69 |
| | 95 | 109 | CFTYIAADLRITLRF | HLA-DRB3*01:01 | 171.1 | 6.25 |
| | 97 | 111 | TYIAADLRITLRFSN | HLA-DRB5*01:01 | 455.2 | 31.8 |
| | 95 | 109 | CFTYIAADLRITLRF | HLA-DQA1*05:01/ DQB1*03:01 | 59.7 | 10.31 |
| | 94 | 108 | SCFTYIAADLRITLR | HLA-DQA1*05:01/ DQB1*03:01 | 60.6 | 10.43 |
| | 93 | 107 | LSCFTYIAADLRITL | HLA-DQA1*05:01/ DQB1*03:01 | 71.6 | 11.8 |
| | 96 | 110 | FTYIAADLRITLRFS | HLA-DQA1*05:01/ DQB1*03:01 | 71.8 | 11.82 |
| **YMAEVPVSA** | 137 | 151 | QMLSNFYMAEVPVSA | HLA-DQA1*05:01/ DQB1*02:01 | 247 | 5.47 |
| | 138 | 152 | MLSNFYMAEVPVSAA | HLA-DQA1*05:01/ DQB1*02:01 | 299 | 6.72 |
| | 139 | 153 | LSNFYMAEVPVSAAT | HLA-DQA1*05:01/ DQB1*02:01 | 393.3 | 8.89 |
| | 140 | 154 | SNFYMAEVPVSAATS | HLA-DQA1*05:01/ DQB1*03:01 | 10 | 1.44 |
| | 141 | 155 | NFYMAEVPVSAATST | HLA-DQA1*05:01/ DQB1*03:01 | 10.5 | 1.56 |
| | 143 | 157 | YMAEVPVSAATSTMV | HLA-DQA1*05:01/ DQB1*03:01 | 10.7 | 1.61 |
| | 142 | 156 | FYMAEVPVSAATSTM | HLA-DQA1*05:01/ DQB1*03:01 | 10.8 | 1.63 |
| | 140 | 154 | SNFYMAEVPVSAATS | HLA-DRB1*01:01 | 5.2 | 0.97 |
| | 141 | 155 | NFYMAEVPVSAATST | HLA-DRB1*01:01 | 6.3 | 1.99 |
| ` | 142 | 156 | FYMAEVPVSAATSTM | HLA-DRB1*01:01 | 7.8 | 3.36 |
| | 143 | 157 | YMAEVPVSAATSTMV | HLA-DRB1*01:01 | 13 | 7.34 |
| | 140 | 154 | SNFYMAEVPVSAATS | HLA-DRB1*03:01 | 385.2 | 12.07 |
| | 140 | 154 | SNFYMAEVPVSAATS | HLA-DRB1*04:01 | 33.2 | 2.21 |
| | 139 | 153 | LSNFYMAEVPVSAAT | HLA-DRB1*04:01 | 39.4 | 2.8 |
| | 141 | 155 | NFYMAEVPVSAATST | HLA-DRB1*04:01 | 47.9 | 3.61 |
| | 138 | 152 | MLSNFYMAEVPVSAA | HLA-DRB1*04:01 | 50.6 | 3.87 |
| | 142 | 156 | FYMAEVPVSAATSTM | HLA-DRB1*04:01 | 65.2 | 5.21 |
| | 143 | 157 | YMAEVPVSAATSTMV | HLA-DRB1*04:01 | 97.1 | 7.88 |
| | 143 | 157 | YMAEVPVSAATSTMV | HLA-DRB1*04:04 | 188 | 19.47 |
| | 142 | 156 | FYMAEVPVSAATSTM | HLA-DRB1*04:04 | 325.9 | 27.62 |
| | 140 | 154 | SNFYMAEVPVSAATS | HLA-DRB1*04:04 | 364.6 | 29.41 |
| | 139 | 153 | LSNFYMAEVPVSAAT | HLA-DRB1*08:02 | 150.4 | 3.01 |
| | 138 | 152 | MLSNFYMAEVPVSAA | HLA-DRB1*08:02 | 249.5 | 5.74 |

| | | | | | | |
|---|---|---|---|---|---|---|
| | 139 | 153 | LSNFYMAEVPVSAAT | HLA-DRB1*09:01 | 20.6 | 0.82 |
| | 140 | 154 | SNFYMAEVPVSAATS | HLA-DRB1*09:01 | 21.8 | 0.92 |
| | 138 | 152 | MLSNFYMAEVPVSAA | HLA-DRB1*09:01 | 22.9 | 1.01 |
| | 141 | 155 | NFYMAEVPVSAATST | HLA-DRB1*09:01 | 34.6 | 1.93 |
| | 143 | 157 | YMAEVPVSAATSTMV | HLA-DRB1*09:01 | 75.1 | 5.16 |
| | 142 | 156 | FYMAEVPVSAATSTM | HLA-DRB1*09:01 | 80.2 | 5.53 |
| | 140 | 154 | SNFYMAEVPVSAATS | HLA-DRB3*01:01 | 499.8 | 11.74 |
| **YTSPLSAIP** | 160 | 174 | SIPYTSPLSAIPTSY | HLA-DRB1*04:01 | 213.4 | 15.79 |
| | 161 | 175 | IPYTSPLSAIPTSYF | HLA-DRB1*04:01 | 244 | 17.49 |
| | 162 | 176 | PYTSPLSAIPTSYFG | HLA-DRB1*04:01 | 279.5 | 19.37 |
| | 159 | 173 | FSIPYTSPLSAIPTS | HLA-DRB1*04:01 | 287.5 | 19.79 |
| | 158 | 172 | SFSIPYTSPLSAIPT | HLA-DRB1*04:01 | 302.8 | 20.56 |
| | 163 | 177 | YTSPLSAIPTSYFGW | HLA-DRB1*04:01 | 309.7 | 20.89 |
| | 160 | 174 | SIPYTSPLSAIPTSY | HLA-DRB1*09:01 | 63 | 4.25 |
| | 161 | 175 | IPYTSPLSAIPTSYF | HLA-DRB1*09:01 | 66.9 | 4.53 |
| | 159 | 173 | FSIPYTSPLSAIPTS | HLA-DRB1*09:01 | 86.2 | 5.97 |
| | 162 | 176 | PYTSPLSAIPTSYFG | HLA-DRB1*09:01 | 96.2 | 6.65 |
